# Tissue Memory Score measures transcriptional retention in solid tumors and predicts survival in clear cell renal cell carcinoma

**DOI:** 10.64898/2026.04.22.718749

**Authors:** Somar Khalil, Marwa Hamdan

## Abstract

Solid tumors retain transcriptional programs inherited from their tissue of origin, but the degree of this retention has not been quantified as a continuous metric, nor has its prognostic relevance been evaluated systematically. We developed the Tissue Memory Score (TMS) by projecting tumor bulk RNA-seq onto principal component axes constructed from 2,694 Genotype-Tissue Expression V8 samples across eight healthy tissues, and applied it to 3,900 primary tumors from eight Cancer Genome Atlas cohorts. Nearest-centroid classification recovered tissue identity in six of eight tumor types (80.1%–100.0% accuracy; chance 12.5%; all p < 10⁻⁵⁵). The two failures, gastric and endometrial carcinoma, reflected limitations of the reference atlas composition and not loss of tissue memory. Among cohorts passing pre-specified retention and dedifferentiation gates, TMS was independently associated with overall survival in clear cell renal cell carcinoma (ccRCC; multivariable HR = 0.59 per SD; 95% CI 0.46–0.77; p = 6.6 × 10⁻⁵) and accounted for 16.3% of explained survival variance by Shapley decomposition. The association held under purity residualization, rank transformation, stage stratification, and progression-free interval as an alternative endpoint. In hepatocellular carcinoma, TMS showed a protective univariable association in the same direction, which did not survive multivariable adjustment and is reported as exploratory. External application to CPTAC-ccRCC yielded a pooled hazard ratio of 0.71 in the same direction (I² = 0%). Tissue-of-origin transcriptional memory is quantifiable in solid tumors and independently prognostic in ccRCC, but did not extend to the remaining seven cohorts. Broader applicability depends on whether the reference atlas represents the cellular compartment from which a given carcinoma arises.

## Introduction

Solid tumors retain transcriptional programs inherited from their tissue of origin. This property is sufficiently pervasive that unsupervised molecular taxonomy of over 10,000 tumors from 33 cancer types clusters primarily by anatomical site rather than by shared oncogenic alterations or recurrent pathway mutations^1^. Tissue-of-origin signal dominates driver mutation signal in bulk transcriptomic data because lineage-specific transcription factor networks established during normal development persist through malignant transformation^2^. These lineage-dependent transcriptional circuits constrain the differentiation states and therapeutic vulnerabilities of each malignancy^3^. The cell type from which a tumor originates determines its molecular identity, clinical behavior, and treatment sensitivity. Yet no quantitative framework exists to measure the degree to which individual tumors retain their tissue-of-origin transcriptional identity, nor has the prognostic relevance of this retention been evaluated systematically in a pan-cancer setting.

The clinical evidence that loss of tissue-specific differentiation correlates with aggressive disease has been documented independently in virtually every solid tumor type for which histological grading systems exist. In clear cell renal cell carcinoma (ccRCC), nuclear grading systems introduced by Fuhrman *et al.*^4^ and refined by the World Health Organization (WHO)/International Society of Urological Pathology (ISUP)^5^ stratify patients on the basis of morphological features that reflect progressive loss of proximal tubular epithelial differentiation. In hepatocellular carcinoma (HCC), the Edmondson-Steiner system^6^ classifies tumors along a four-tier dedifferentiation axis that remains prognostically relevant six decades after its introduction. In prostate adenocarcinoma, the Gleason grading system^7,8^ constitutes one of the most direct dedifferentiation scores in clinical oncology: loss of glandular architecture defines high-grade disease with elevated metastatic potential and cancer-specific mortality. Analogous dedifferentiation-outcome associations have been documented in colorectal^9^, gastric^10^, endometrial^11^, and thyroid carcinoma^12^. That dedifferentiation predicts poor outcome in each of these organ systems supports the hypothesis: that the degree of tissue-of-origin transcriptional identity retained by a tumor is a continuous, measurable property with prognostic information content independent of clinical stage, tumor purity, and mutational burden. A quantitative score capturing tissue identity retention could complement existing grading systems by providing a molecular, continuous, and reproducible metric that is not subject to interobserver variability and can be computed from standard RNA sequencing data.

Several computational approaches have addressed related questions. Malta *et al.*^13^ developed machine-learning-derived stemness indices (mRNAsi, mDNAsi) by training a one-class logistic regression classifier on pluripotent stem cell expression signatures, quantifying the degree to which a tumor resembles stem cells. This property is related to but not equivalent to tissue identity retention: stemness measures what tumors gain along the pluripotency axis, whereas tissue identity retention measures what tumors preserve of their differentiated program. Ben-Porath *et al.*^14^ demonstrated enrichment of embryonic stem cell gene expression signatures in poorly differentiated breast cancers, providing early evidence that transcriptional stemness correlates with clinical aggressiveness. Gene expression profiling for tumors of unknown primary^15^ addresses a categorical classification problem but does not quantify the degree of identity retention as a continuous variable whose contribution to survival variance can be formally quantified. Organ-specific molecular profiling studies have demonstrated that lineage-associated epigenomic and transcriptomic signatures carry prognostic information in individual cancer types^16^ but do not generalize to a pan-cancer framework. No study has constructed a continuous, atlas-derived transcriptomic score from a matched healthy tissue reference and evaluated its independent prognostic contribution with formal variance attribution in a pan-cancer design.

The Genotype-Tissue Expression (GTEx) project^17,18^ provides the first adequately powered, uniformly processed, multi-tissue bulk RNA-seq resource capable of serving as a healthy baseline for this purpose. Its bulk RNA-seq format is directly compatible with TCGA tumor expression data^19^, eliminating the pseudo-bulk aggregation step required by single-cell atlases. Whether the resulting score is informative depends on how well GTEx tissue samples capture the specific cellular compartment from which a given tumor type arises; where the reference is dominated by non-epithelial populations, the principal component analysis (PCA) axes may not encode the transcriptional programs relevant to epithelial malignancy.

A second methodological gap motivates this study. The standard approach to evaluating a novel prognostic biomarker reports a hazard ratio from a multivariable Cox model without quantifying the fraction of survival variance the biomarker uniquely explains. Shapley decomposition of the model pseudo-*R*²^20,21^ provides an order-independent variance attribution by averaging each predictor’s marginal contribution over all possible predictor orderings. Complementing variance attribution, net reclassification improvement (NRI)^22,23^ and decision curve analysis (DCA)^24^ quantify whether adding a biomarker to an existing clinical model improves patient classification at actionable risk thresholds.

This study introduces the Tissue Memory Score (TMS), a continuous metric computed by projecting tumor bulk RNA-seq expression onto principal component axes derived from a matched healthy tissue reference (GTEx V8, eight tissues, 2,694 samples). TMS is defined as the negative Euclidean distance from a tumor’s projection in PCA space to the centroid of its matched healthy tissue, such that higher values denote greater retention of tissue-of-origin transcriptional identity. Four pre-specified claims are evaluated through a sequential decision gate locked prior to survival analysis: tissue identity retention (C1, nearest-centroid accuracy >60%), attenuation with dedifferentiation (C2, |Cohen *d*| > 0.3), independent prognostic value (C3, multivariable Cox regression with Shapley decomposition), and cross-cohort directional consistency (C4, random-effects meta-analysis^25^). Covariates include ordinal AJCC stage, ESTIMATE-derived tumor purity^26,27^, and tumor mutational burden from Multi-Center Mutation Calling in Multiple Cancers (MC3) somatic mutation calls^28^, with harmonized clinical endpoints from the TCGA Clinical Data Resource^29^. External validation is conducted by applying the locked PCA reference without parameter re-estimation. The study is applied to 3,900 primary tumors from eight TCGA projects (COAD, KIRC, LIHC, PRAD, STAD, UCEC, LUAD, THCA), representing eight tissue lineages with matched GTEx V8 healthy references.

## Materials and Methods

The study protocol, including decision gate criteria and primary model specifications, was finalized before execution of survival analyses; no post-hoc modifications were made (Figure 1).

**Figure 1.**
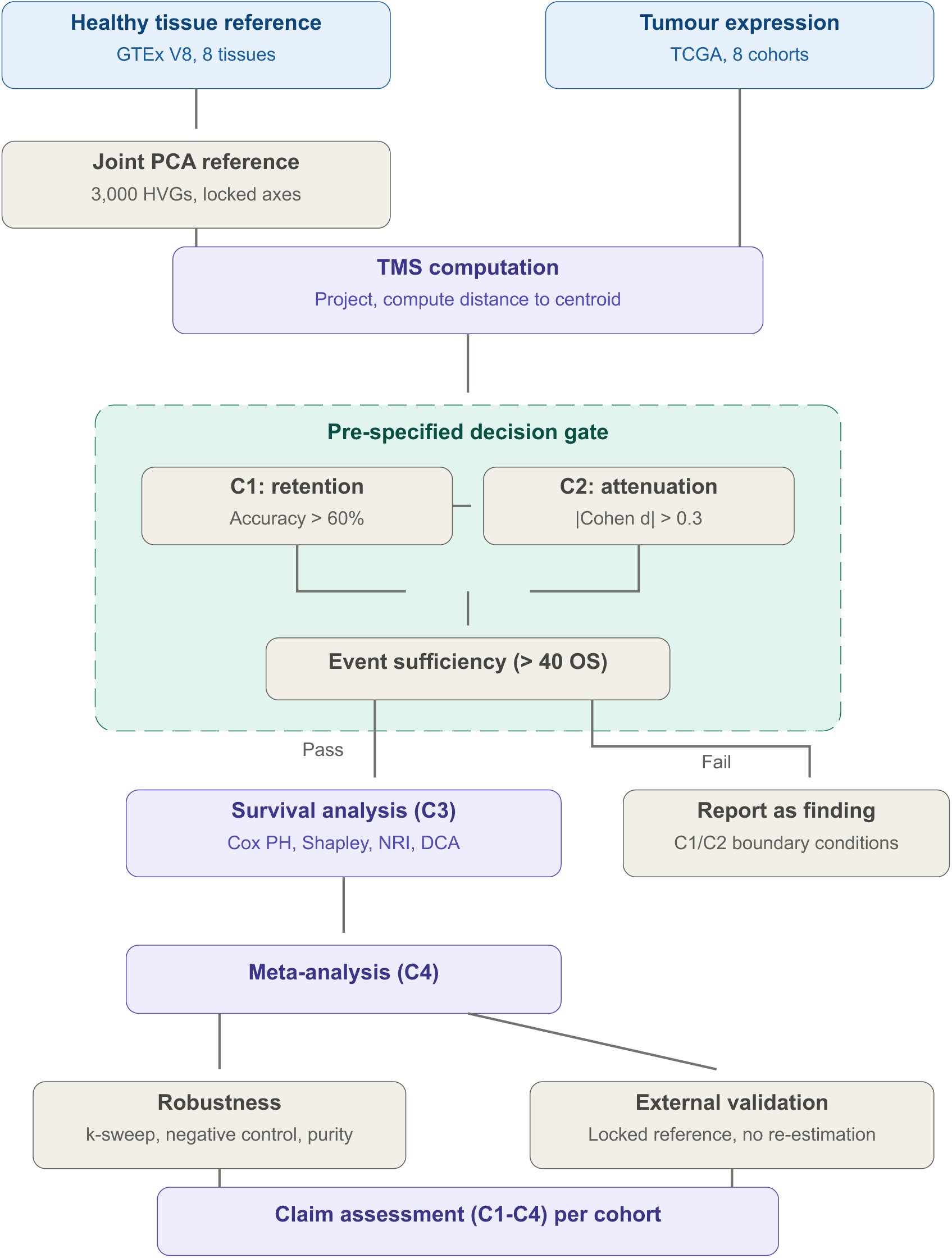
Pre-specified study design. GTEx V8 reference PCA (2,694 samples, eight tissues) was constructed and locked prior to tumor projection. TMS was computed as the negative Euclidean distance from each tumor’s PCA projection to its matched tissue centroid. A sequential decision gate classified cohorts as eligible (C1: nearest-centroid accuracy >60%; C2: stage- or grade-based |Cohen *d*| > 0.3; event suXiciency: >=40 OS events) or ineligible for survival analysis. Eligible cohorts proceeded to multivariable Cox regression with Shapley variance decomposition, clinical utility assessment, and meta-analysis. Cohorts failing C1 or C2 were reported as boundary condition findings. Robustness analyses and external validation using the locked reference without re-estimation were applied to eligible cohorts.

### Data sources

Gene-level read counts and sample metadata for the healthy tissue reference were obtained from the GTEx V8 release^17,18^ (STAR v2.5.3a alignment to GRCh38, RNA-SeQC v1.1.9 quantification). Eight tissues were selected to match the TCGA cohorts: Colon-Transverse, Kidney-Cortex, Liver, Prostate, Stomach, Uterus, Lung, and Thyroid (85--653 samples per tissue; 2,694 total). STAR-Counts gene expression matrices for eight TCGA projects^19^ (COAD, KIRC, LIHC, PRAD, STAD, UCEC, LUAD, THCA) were downloaded from the NCI Genomic Data Commons; only primary tumor samples were retained (3,900 total). Harmonized overall survival (OS), progression-free interval, and American Joint Committee on Cancer (AJCC) pathological stage were obtained from the TCGA Clinical Data Resource^29^. Histological grade was extracted from GDC clinical supplements where applicable (Supplementary Methods S2). ESTIMATE tumor purity was computed locally using a corrected single-sample gene set enrichment analysis (ssGSEA) implementation following Barbie *et al.*^26^ and Yoshihara *et al.*^27^ (Supplementary Methods S1). Tumor mutational burden (TMB) was computed from MC3 somatic mutation calls^28^ as non-silent mutations per megabase (log10-transformed). Two independent validation cohorts were used: Clinical Proteomic Tumor Analysis Consortium (CPTAC)-3 ccRCC (261 RNA-seq samples; 35 with OS annotation) and GSE76427 HCC (167 microarray samples; 23 events).

### Healthy tissue reference construction and TMS computation

GTEx read counts were normalized to log2(counts per million [CPM] + 1). The top 3,000 genes by variance were selected as highly variable genes (HVGs). Joint PCA on the 2,694 x 3,000 matrix yielded *k* = 8 retained components (75.5% cumulative variance). Bootstrap stability was confirmed by 200 donor-level resamples (mean subspace angle 3.30 degrees; threshold <15 degrees). Tissue centroids were computed as the arithmetic mean of projected coordinates per organ. All reference objects were serialized and permanently locked after construction. TCGA expression matrices were normalized identically, projected onto the locked PCA axes, and the TMS was computed as:

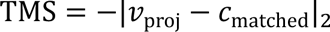

where 𝑣_proj_ is the tumor’s projection and 𝑐_matched_ is the matched healthy tissue centroid. Higher TMS denotes greater retention of tissue-of-origin transcriptional identity. TMS was standardized to zero mean and unit variance within each cohort (TMS_z_) for all downstream analyses.

### Statistical analysis

Nearest-centroid classification accuracy was computed per cohort and tested against the eight-tissue chance level (12.5%) by exact binomial test (C1). The 60% accuracy threshold was set as a conservative floor to accommodate expected classification heterogeneity arising from variable transcriptional distinctiveness of different tissue lineages in the joint PCA space, while remaining substantially above the 12.5% chance level. Stage-based C2 compared mean TMS between early-stage (I/II) and late-stage (III/IV) tumors by Welch t-test and Cohen *d*; grade-based C2 used Spearman correlation where histological grade was available. The |*d*| > 0.3 threshold was adopted from Cohen’s conventional benchmark for a small-to-medium effect size and was pre-specified prior to data inspection; it was not empirically calibrated to the distributional properties of TMS.

Spearman correlations between TMS and covariates (purity, TMB, stage) were computed per cohort; purity-residualized TMS was derived for cohorts with |ρTMS-purity| > 0.5 (Supplementary Methods S3). The decision gate summary, reporting per-cohort passage and failure, constitutes a primary study finding (Figure 1).

For gate-passing cohorts (KIRC, LIHC), univariable Cox proportional hazards models were fitted for TMS_z, ordinal stage, ESTIMATE purity, and log10 TMB using the lifelines Python package (v0.27). The primary significance threshold was Bonferroni-adjusted at *p* < 0.00625 (0.05/8 cohorts). This correction was applied to the eight univariable Cox tests (one per cohort) to control the family-wise error rate for the primary screening of TMS-OS associations. Multivariable models for gate-passing cohorts were evaluated at nominal α = 0.05, as the decision gate had already constrained the analytic space; sensitivity analyses (progression-free interval, stage-stratified, purity-residualized, and rank-normalized models) were likewise conducted at nominal α and interpreted as confirmatory within gate-passing cohorts.

A full multivariable model including all four covariates was fitted per cohort; collinearity was assessed by variance inflation factors (VIF) and model fit by Akaike information criterion (AIC) and concordance index. The proportional hazards assumption was evaluated by Spearman correlation of scaled Schoenfeld residuals with ranked event times; no violation was detected in either cohort. Shapley decomposition of the Royston-Sauerbrei pseudo-R-squared^20,21^ attributed explained survival variance to each covariate by evaluating all 2^4^ covariate subsets, with bootstrap confidence intervals from 500 case-level resamples. Kaplan-Meier analysis by TMS tertiles was performed with log-rank testing and restricted mean survival time (RMST) estimation.

Univariable TMS hazard ratios from the two eligible cohorts were assessed for directional consistency using DerSimonian-Laird random-effects pooling^25^, with heterogeneity quantified by I² and Cochran Q. With only two contributing cohorts, this procedure is presented as a directional concordance assessment; the I² point estimate is not meaningfully interpretable at this sample size (see Results).

For KIRC, clinical utility was assessed by category-free net reclassification improvement (NRI)^22,23^ and decision curve analysis (DCA)^24^ at three-year and five-year horizons, comparing stage-only versus stage + TMS models.

The testing hierarchy was pre-specified at four levels: gatekeeping (C1/C2, sequential without adjustment), primary inference (univariable Cox, Bonferroni-adjusted *p* < 0.00625), secondary inference (multivariable HRs, NRI, DCA; nominal alpha = 0.05), and exploratory (molecular subtype, grade-stratified, and validation subgroup analyses; hypothesis-generating).

### External validation

The locked PCA reference was applied to each validation cohort without re-estimation. TMS was computed by projection and distance to the matched centroid (Supplementary Methods S11). Replication was assessed by three pre-specified criteria: concordant HR direction (R1), univariable *p* < 0.05 (R2), and overlapping 95% CIs with discovery (R3). Fixed-effect meta-analysis pooled discovery and validation ccRCC cohorts.

### Software and data availability

All analyses were performed in Python 3.12 (scikit-learn v1.4, lifelines v0.27, SciPy v1.12, statsmodels v0.14). All source data are publicly available from the GTEx Portal, NCI GDC, and NCBI GEO.

## Results

### Healthy tissue reference and characterization (C1, C2)

Joint principal component analysis of 2,694 GTEx V8 samples spanning eight tissues (colon, kidney, liver, prostate, stomach, uterus, lung, thyroid) identified a stable tissue-identity coordinate system. The first eight principal components were retained (*k* = 8), collectively explaining 75.5% of total variance (Figure 2B). The first three components captured the majority of inter-tissue separation, with tissue-pair centroid distances exceeding within-tissue spread for most pairs (Figure 2A,C). Subject-level bootstrap resampling (200 iterations) yielded a mean subspace angle of 3.30° between resampled and reference PCA axes, well within the pre-specified 15° stability threshold (Figure 2D; Table S14). Thyroid, prostate, and liver formed the most distinct clusters in PC1-PC2 space, while colon and stomach exhibited partial overlap consistent with their shared gastrointestinal mucosal lineage (Figure 2A). Per-tissue PCA (Strategy B) confirmed that within-tissue variance structure was stable for all eight organs.

**Figure 2.**
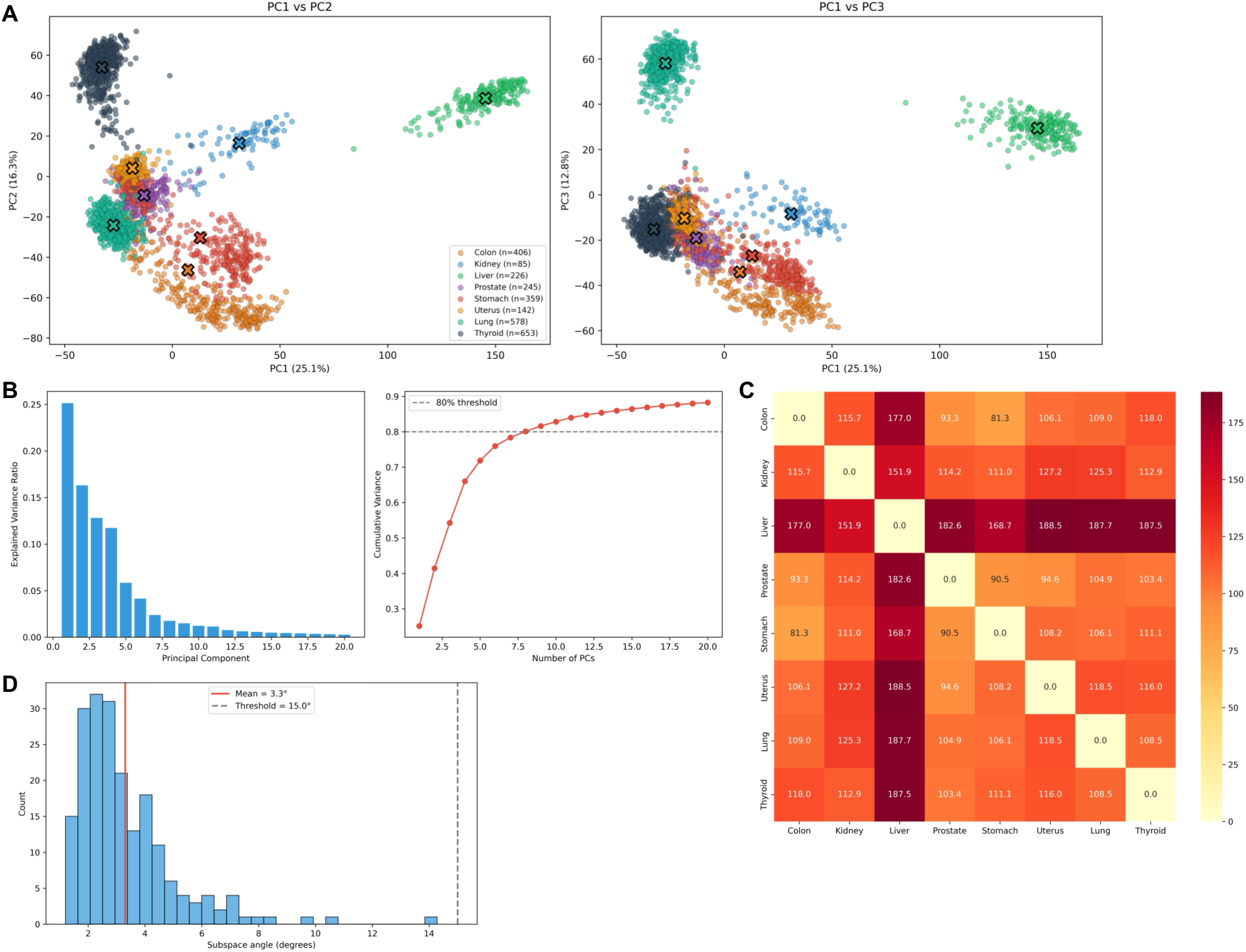
Healthy tissue reference construction. (A) Joint PCA of 2,694 GTEx V8 samples spanning eight tissues, shown in PC1 vs PC2 (left) and PC1 vs PC3 (right) space. Crosses denote tissue centroids. (B) Scree plot of per-component explained variance (left) and cumulative explained variance as a function of the number of retained PCs (right); dashed line indicates the 80% threshold. (C) Pairwise Euclidean centroid distance matrix among the eight tissue centroids in the full PCA space. (D) Distribution of principal subspace angles between 200 subject-level bootstrap PCA resamples and the reference PCA. Red line, observed mean (3.3°); dashed line, pre-specified 15° stability threshold.

Nearest-centroid classification of 3,900 primary tumors against the eight GTEx tissue centroids demonstrated tissue identity retention in six cohorts (Table 1; Figure 3A). PRAD achieved 100% accuracy (501/501; *p* < 10⁻³⁰⁰), followed by THCA (99.2%, 501/505), COAD (98.9%, 474/479), LIHC (96.0%, 356/371), KIRC (95.9%, 518/540), and LUAD (80.2%, 432/539); all six exceeded the 60% pre-specified threshold and were significant against the 12.5% chance level (all *p* < 10⁻⁵⁵). Projection of KIRC tumors onto the reference PCA axes confirmed dense clustering near the kidney centroid with minimal overlap into adjacent tissue regions (Figure 3C).

**Figure 3.**
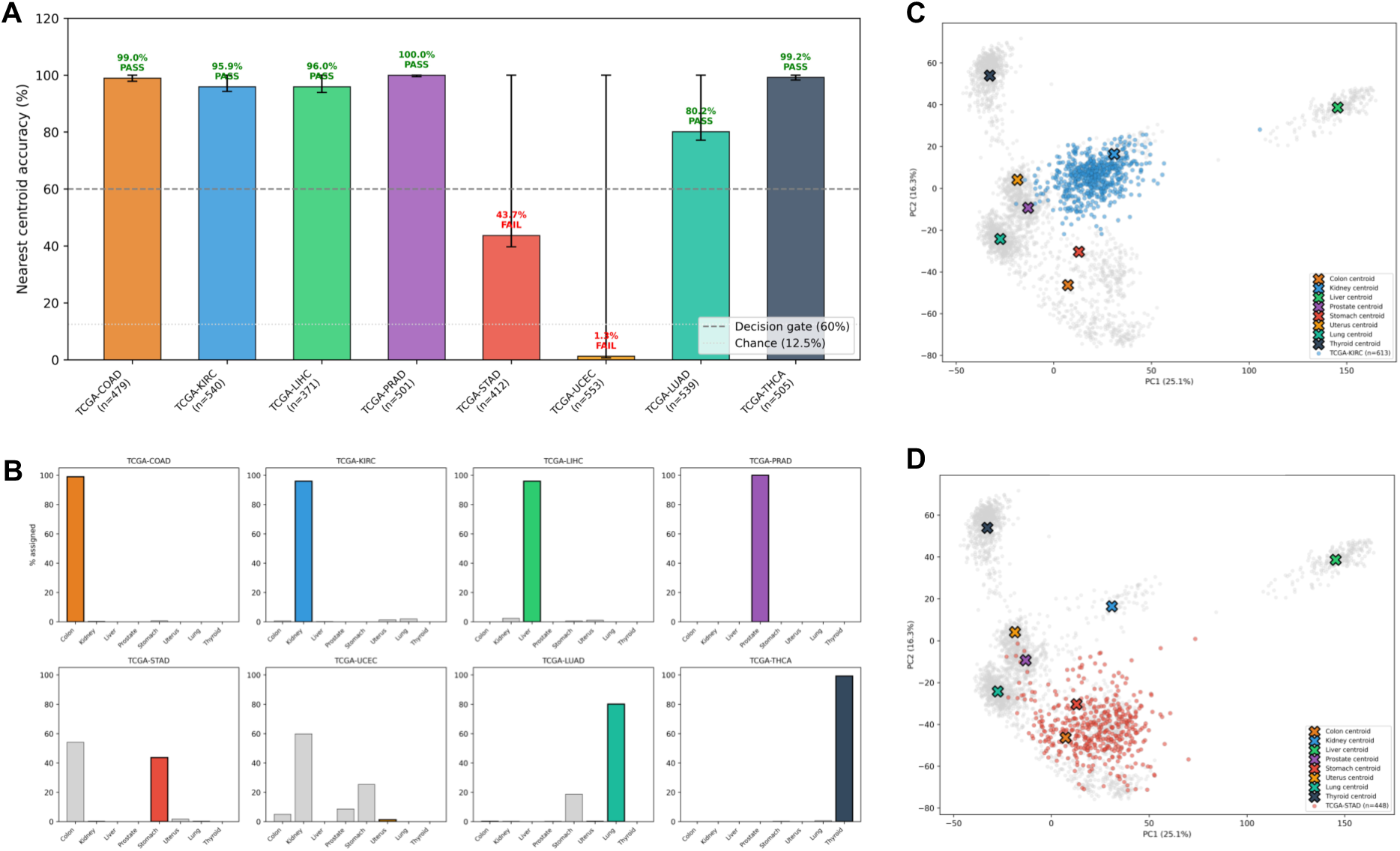
Tissue identity retention (C1). (A) Nearest-centroid classification accuracy for each of the eight TCGA cohorts. Dashed line, pre-specified 60% decision gate threshold; dotted line, 12.5% chance level. Error bars denote exact binomial 95% confidence intervals. (B) Per-cohort classification distribution barplots showing the proportion of tumors assigned to each of the eight tissue centroids. STAD tumors misclassify predominantly to the colon centroid; UCEC tumors distribute diffusely with no dominant assignment. (C) TCGA-KIRC primary tumors (blue) projected onto the GTEx reference PCA (grey), illustrating dense co-localization with the kidney centroid (C1 success). (D) TCGA-STAD primary tumors (red) projected onto the GTEx reference PCA, illustrating dispersion between the stomach and colon centroids (C1 failure).

**Table 1.** Nearest-centroid classification of tissue-of-origin identity (C1). Each primary tumor was classified to the nearest GTEx tissue centroid in the locked PCA space. Accuracy denotes the proportion assigned to the matched organ. P values from exact binomial test against the 12.5% chance level (1/8 tissues). C1 pass, accuracy exceeding the pre-specified 60% threshold.

| Cohort | Matched organ | N | Accuracy | 95% CI low | 95% CI high | *P* (binomial vs 12.5%) | C1 pass |
| --- | --- | --- | --- | --- | --- | --- | --- |
| COAD | Colon | 479 | 0.989 | 0.975 | >0.999 | <10<sup>-300</sup> | Yes |
| KIRC | Kidney | 540 | 0.959 | 0.938 | >0.999 | <10<sup>-300</sup> | Yes |
| **LIHC** | Liver | 371 | 0.960 | 0.936 | >0.999 | $6.9 \times 10^{-297}$ | **Yes** |
| **PRAD** | Prostate | 501 | 1.000 | 0.993 | >0.999 | $<10^{-300}$ | **Yes** |
| **STAD** | Stomach | 412 | 0.437 | 0.389 | >0.999 | $1.9 \times 10^{-55}$ | No |
| **UCEC** | Uterus | 553 | 0.013 | 0.005 | >0.999 | 1.000 | No |
| **LUAD** | Lung | 539 | 0.802 | 0.767 | >0.999 | $9.2 \times 10^{-282}$ | **Yes** |
| **THCA** | Thyroid | 505 | 0.992 | 0.980 | >0.999 | $<10^{-300}$ | **Yes** |

Two cohorts failed C1. STAD achieved 43.7% accuracy (180/412), with the majority of misclassified gastric tumors assigned to the colonic centroid (Figure 3B), consistent with the transcriptional overlap between gastric and colonic mucosal epithelium observed in the GTEx reference PCA. The STAD projection overlay illustrates this lineage-level blurring (Figure 3D). UCEC failed with 1.3% accuracy (7/553), a near-complete inability to assign endometrial tumors to the uterine centroid. This failure is attributable to the cellular composition of GTEx Uterus, which samples predominantly myometrial smooth muscle and endometrial stroma rather than the glandular epithelium from which endometrioid carcinomas arise. The resulting PCA axes capture myometrial transcriptional identity and do not represent the endometrial epithelial programs retained by UCEC tumors. These two failures define the boundary conditions of the framework: the tissue memory framework requires that the reference atlas faithfully represent the cellular compartment of tumor origin, and the STAD and UCEC results delineate the conditions under which this requirement is not met. Individual cohort projections for all eight tumor types are provided in Figure S1.

TCGA-matched normal tissue samples exhibited higher TMS than paired primary tumors in seven of eight cohorts, confirming the expected direction of tissue memory loss in malignancy (Supplementary Methods S4; Table S28). The largest effects were observed in LUAD (Cohen *d* = 2.50; *p* = 7.5 × 10⁻⁵⁶), KIRC (d = 1.69; *p* = 1.1 × 10⁻²⁶), THCA (d = 1.21; *p* = 6.7 × 10⁻¹²), and LIHC (d = 0.96; *p* = 1.9 × 10⁻²⁸). UCEC showed a particularly large effect (d = 4.11; *p* = 2.2 × 10⁻¹⁵), consistent with the dramatic separation between myometrial normals and endometrial tumors in PCA space. STAD was the sole exception: normal tissue TMS was lower than tumor TMS (*d* = −0.64; *p* = 0.025), an inversion that is attributable to the heterogeneous cellular composition of GTEx Stomach (dominated by muscularis propria, submucosa, and serosa, with limited representation of the foveolar and glandular epithelium from which gastric adenocarcinoma arises) relative to the mucosal epithelial origin of gastric adenocarcinoma.

Stage-based C2 testing revealed significant TMS attenuation in two cohorts (Figure 4B,C). LIHC showed the largest effect (Cohen *d* = 0.50; 95% CI 0.26-0.74; Welch *p* = 7.2 × 10⁻⁴), with late-stage tumors (stages III-IV; mean TMS = −45.8) scoring substantially lower than early-stage tumors (stages I-II; mean TMS = −34.4). KIRC followed with d = 0.42 (95% CI 0.25-0.60; p = 7.0 × 10⁻⁶), late-stage mean TMS = −49.7 versus early-stage mean TMS = −44.4. Both exceeded the pre-specified |d| > 0.3 threshold (Figure 4C; Table S30). The remaining C1-passing cohorts did not exhibit stage-dependent TMS attenuation: THCA (d = 0.17), LUAD (d = 0.15), COAD (d = −0.14), and PRAD (d = 0.01), none reaching the 0.3 threshold and none achieving statistical significance. Grade-based C2 testing was conducted for PRAD (ISUP Grade Group), STAD (WHO grade), and UCEC (International Federation of Gynecology and Obstetrics [FIGO] grade) (Figure S2B; Table S31). STAD and UCEC showed significant Spearman correlations between TMS and grade (*ρ* = 0.20, *p* = 2.7 × 10⁻⁴ and *ρ* = 0.23, *p* = 8.0 × 10⁻⁷, respectively), but both failed C1 and therefore did not proceed to survival analysis. PRAD showed no grade-TMS association (*ρ* = −0.003; *p* = 0.95). Per-cohort TMS distributions are shown in Figure 4A.

**Figure 4.**
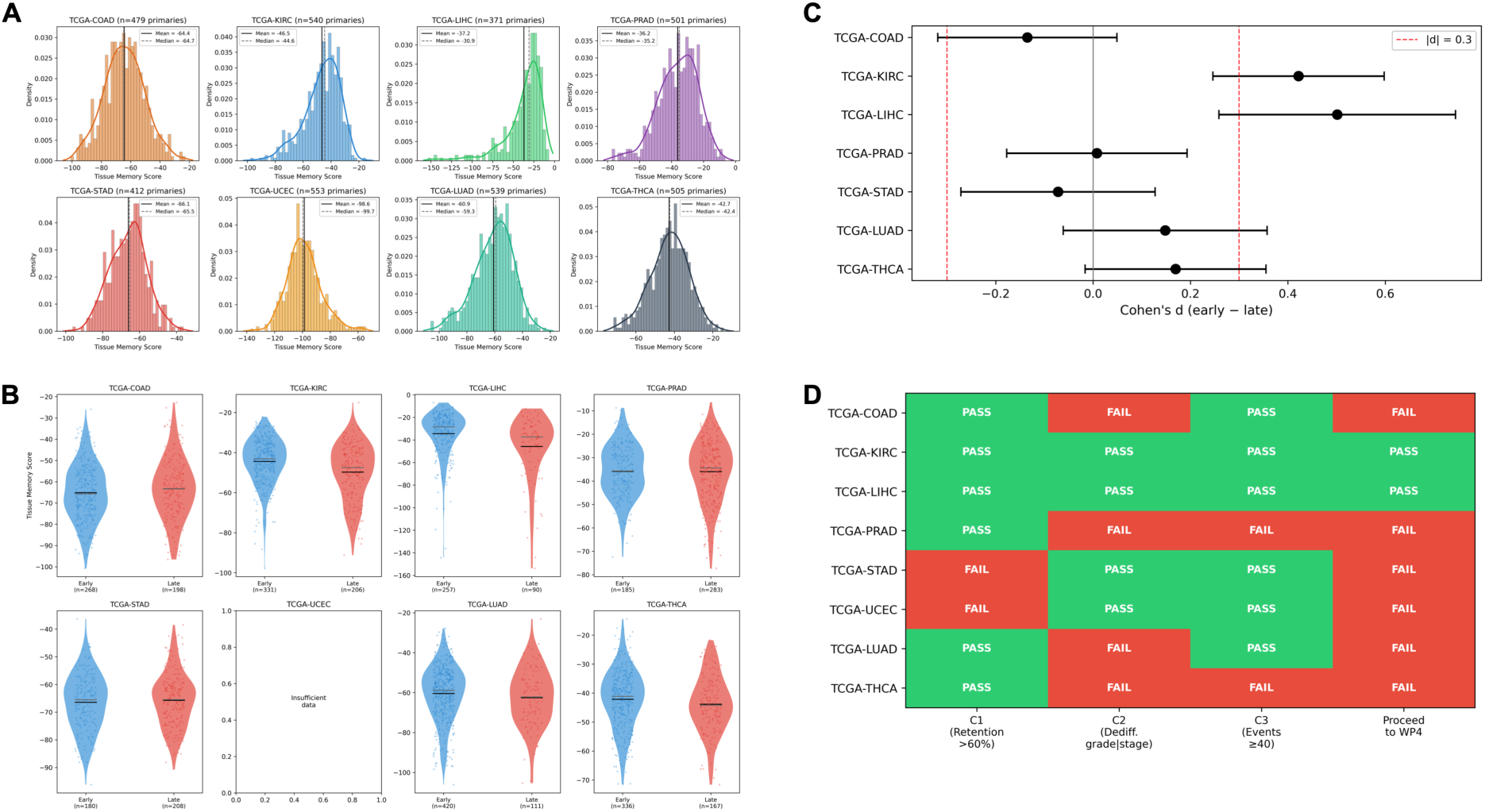
TMS characterization, attenuation, and decision gate (C2). (A) Per-cohort TMS density distributions for all eight TCGA cohorts. Vertical lines indicate cohort mean and median. (B) TMS distributions stratified by early (stages I–II) versus late (stages III–IV) AJCC pathological stage per cohort. UCEC is omitted due to insufficient stage annotation. (C) Forest plot of Cohen *d* effect sizes for the stage-based C2 test. Red dashed line, pre-specified |d| = 0.3 threshold. Error bars denote 95% confidence intervals. (D) Decision gate summary. Each cohort is classified as pass (green) or fail (red) for C1 (retention >60%), C2 (dedifferentiation |d| > 0.3), and event sufficiency (≥40 OS events), and eligibility to proceed to survival analysis. LIHC passed all gates but did not yield a confirmed C3 result (see Results).

TMS-purity correlations varied substantially by cohort (Figure S2A; Table S27). KIRC exhibited a strong positive correlation (Spearman *ρ* = 0.71; *p* = 7.5 × 10⁻⁸⁴), indicating that tumors with higher stromal/immune admixture (lower purity) tend to score closer to their matched healthy tissue centroid. LUAD showed a strong inverse correlation (*ρ* = −0.63; *p* = 1.5 × 10⁻⁶¹). Purity residualization in KIRC reduced the TMS-purity correlation from *ρ* = 0.71 to *ρ* = 0.02, after which the residualized TMS and purity were uncorrelated (Table S29). LUAD residualization similarly abolished the correlation (ρ = −0.63 to ρ = −0.02). TMS-TMB correlations were weak in all cohorts (|ρ| ≤ 0.23). TMS-stage correlations were modest, with the strongest in LIHC (ρ = −0.27) and KIRC (ρ = −0.20).

The pre-specified decision gate classified each cohort for survival analysis eligibility (Figure 4D; Table 2). Two cohorts met all criteria: KIRC (C1: 95.9%, pass; C2 stage: d = 0.42, pass; events: 175) and LIHC (C1: 96.0%, pass; C2 stage: d = 0.50, pass; events: 130). Six cohorts were excluded: four passed C1 but failed C2 (COAD, PRAD, LUAD, THCA), and two failed C1 (STAD, UCEC). THCA was additionally excluded a priori due to insufficient OS events (∼15). The decision gate outcomes constitute a primary result: tissue-of-origin identity retention is widespread (6/8), but its association with dedifferentiation status is restricted to tumor types where the primary differentiation axis maps onto the stage classification system.

**Table 2.** Per-cohort decision gate summary with univariable prognostic associations. C1 Acc., nearest-centroid classification accuracy; C1, passage of 60% retention threshold; C2 *d*, Cohen *d* for early- vs. late-stage TMS difference; C2, passage of |*d*| > 0.3 threshold; Univ. HR, univariable Cox hazard ratio per standard deviation increase in TMS (overall survival); Eligible, passage of both C1 and C2 gates with ≥40 OS events. *†Fewer than 20 OS events (PRAD: 10; THCA: 16); hazard ratio estimates are unstable and reported for completeness. ‡LIHC passed both C1 and C2 gates and proceeded to survival analysis; however, the univariable association did not meet the Bonferroni-corrected threshold (p = 0.008 vs. required p < 0.00625), and the multivariable association was non-significant (p = 0.11). LIHC results are classified as exploratory*.

| Cohort | Organ | N | C1 Acc. | C1 | C2 *d* | C2 | Univ. HR (95% CI) | Univ. *P* | Eligible‡ |
| --- | --- | --- | --- | --- | --- | --- | --- | --- | --- |
| COAD | Colon | 479 | 0.989 | Yes | −0.14 | No | 1.09 (0.90–1.32) | 0.373 | No |
| KIRC | Kidney | 540 | 0.959 | Yes | 0.42 | Yes | 0.71 (0.62–0.81) | $3.3 \times 10^{-7}$ | Yes |
| LIHC | Liver | 371 | 0.960 | Yes | 0.50 | Yes | 0.83 (0.72–0.95) | 0.008 | Yes |
| PRAD | Prostate | 501 | 1.000 | Yes | 0.01 | No | 0.62 (0.33–1.16) | 0.133† | No |
| STAD | Stomach | 412 | 0.437 | No | −0.07 | No | 1.14 (0.97–1.35) | 0.112 | No |
| UCEC | Uterus | 553 | 0.013 | No | — | No | 1.12 (0.91–1.39) | 0.271 | No |
| LUAD | Lung | 539 | 0.802 | Yes | 0.15 | No | 0.87 (0.76–1.00) | 0.051 | No |
| THCA | Thyroid | 505 | 0.992 | Yes | 0.17 | No | 1.02 (0.62–1.69) | 0.925† | No |

### Prognostic value of TMS (C3)

Univariable Cox regression demonstrated a significant association between TMS and OS in KIRC (HR = 0.71 per SD; 95% CI 0.62-0.81; *p* = 3.3 × 10⁻⁷; *n* = 538, 175 events; concordance = 0.62), surpassing the Bonferroni-corrected threshold of *p* < 0.00625 (Table S1). In the full multivariable model (TMS, stage, purity, TMB; *n* = 369, 97 events), TMS retained independent significance (HR = 0.59; 95% CI 0.46-0.77; *p* = 6.6 × 10⁻⁵; model concordance = 0.76; Figure 5B; Table 3; Table S2). All four covariates contributed significantly: stage (HR = 2.01; *p* = 1.4 × 10⁻¹³), purity (HR = 4.83; *p* = 0.008), and TMB (HR = 1.17; *p* = 2.3 × 10⁻⁴). VIF confirmed no problematic collinearity (VIF_TMS = 2.30, VIF_Purity = 2.33, VIF_Stage = 1.05, VIF_TMB = 1.01; Table S4).

**Figure 5.**
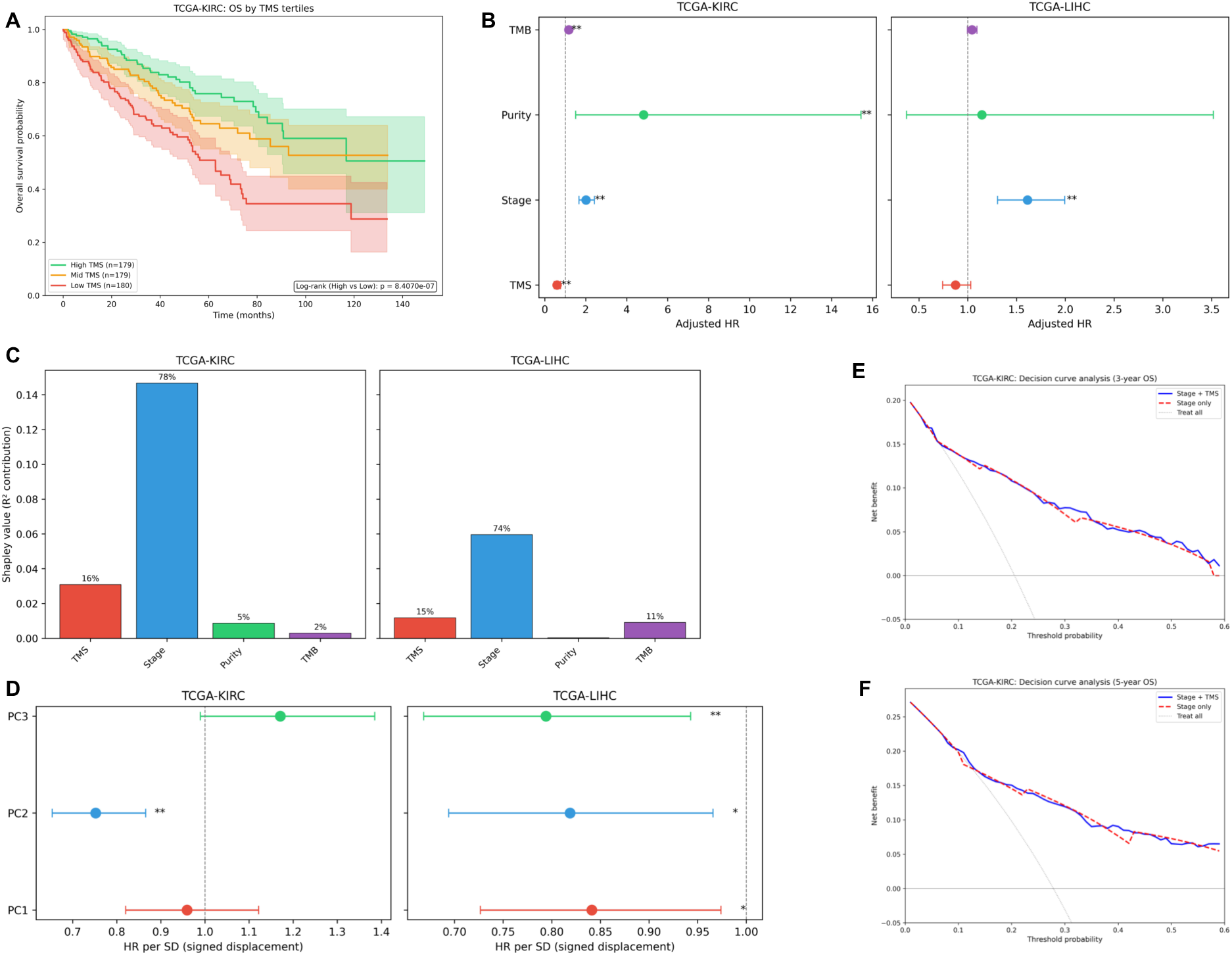
Prognostic value and mechanistic decomposition of TMS in ccRCC (C3). (A) Kaplan--Meier overall survival curves by TMS tertiles in TCGA-KIRC (*n* = 538; 175 events). Shaded regions denote 95% confidence intervals. (B) Multivariable Cox proportional hazards forest plot for TCGA-KIRC (left) and TCGA-LIHC (right). Hazard ratios are per standard deviation for continuous covariates and per unit increase for ordinal stage. (C) Shapley decomposition of the Royston--Sauerbrei pseudo-*R*² for the full four-covariate Cox model in KIRC (left) and LIHC (right). Percentages denote the proportion of total explained survival variance attributed to each covariate. (D) Per-principal-component univariable Cox hazard ratios (signed displacement, standardized) in KIRC (left) and LIHC (right). Asterisks denote significance (\**p* < 0.05, \*\**p* < 0.01). (E) Decision curve analysis at the three-year overall survival horizon in KIRC, comparing net benefit of stage-only versus stage + TMS models. (F) Decision curve analysis at the five-year horizon.

**Table 3.** Prognostic models for tissue memory score in clear cell renal cell carcinoma (TCGA-KIRC). Hazard ratios are per standard deviation increase in each continuous covariate and per unit increase for ordinal stage. Shapley % denotes the proportion of the Royston–Sauerbrei pseudo-*R*² attributed to each covariate by Shapley decomposition (500 bootstrap resamples). PFI, progression-free interval. All models use overall survival unless otherwise noted.

| Covariate | Univariable HR (95% CI) | *P* | Multivariable HR (95% CI) | *P* | Shapley % | *n* / events | C-index |
| --- | --- | --- | --- | --- | --- | --- | --- |
| TMS | 0.71 (0.62–0.81) | $3.3 \times 10^{-7}$ | 0.59 (0.46–0.77) | $6.6 \times 10^{-5}$ | 16.3% | 538 / 175 | |
| Stage | 1.89 (1.66–2.16) | $1.1 \times 10^{-21}$ | 2.01 (1.67–2.42) | $1.4 \times 10^{-13}$ | 77.6% | | |
| Purity | 0.45 (0.25–0.83) | 0.010 | 4.83 (1.51–15.4) | 0.008 | 4.6% |  |  |
| TMB | 1.12 (1.04–1.22) | 0.003 | 1.17 (1.08–1.27) | $2.3 \times 10^{-4}$ | 1.5% | | |
| **Full model** |  |  |  |  | **$R^2 = 0.189$** | **369 / 97** | **0.76** |
| **Sensitivity analyses** |  |  |  |  |  |  |  |
| PFI endpoint | — | — | 0.58 (0.45–0.76) | $4.7 \times 10^{-5}$ | — | 367 / 94 | 0.83 |
| Purity-residualized | 0.75 (0.67–0.84) | $1.1 \times 10^{-6}$ | 0.70 (0.59–0.84) | $6.3 \times 10^{-5}$ | — | 369 / 97 | 0.76 |
| Stage-stratified | — | — | 0.60 (0.46–0.77) | $8.0 \times 10^{-5}$ | — | 369 / 97 | 0.64 |
| TMS + mRNAsi | — | — | 0.61 (0.51–0.74) | $2.3 \times 10^{-7}$ | — | 518 / 168 | 0.76 |
| Intermediate stage (II–III) | 0.78 (0.62–0.99) | 0.039 | 0.56 (0.43–0.73) | $2.3 \times 10^{-5}$ | — | 181 / 62 | 0.68 |

The reversal of the purity hazard ratio from protective in the univariable model (HR = 0.45) to deleterious in the multivariable model (HR = 4.83) reflects the redistribution of shared variance between TMS and purity (Spearman *ρ* = 0.71) upon mutual adjustment. In the univariable model, purity absorbs the TMS-associated survival signal through their correlation; conditional on TMS, the residual purity effect captures a distinct, oppositely directed association. This is a standard consequence of collinearity in multivariable regression, not a sign of model instability, as confirmed by the VIF of 2.33, well below conventional thresholds.

Reciprocal residualization (purity regressed on TMS, OLS R² = 0.539) yielded a purity-residual HR of 2.19 (95% CI 0.86–5.63; p = 0.10) in the univariable model. In the multivariable model with residualized purity, the purity coefficient was 4.83 (95% CI 1.51–15.44; p = 0.008), identical to the primary model, with all VIFs near 1.0, confirming that the sign reversal is a symmetric consequence of shared variance redistribution (Table S16b).

Schoenfeld residual testing (Supplementary Methods S5) confirmed that the proportional hazards assumption was not violated for any covariate (TMS: *ρ* = −0.19, *p* = 0.064; all others *p* > 0.23; Figure S3A; Table S3). Time-varying coefficient analysis demonstrated stable TMS effects across early and late follow-up periods: at the median cutpoint (1,133 days), early-period HR = 0.73 (*p* = 0.005) and late-period HR = 0.67 (*p* = 0.016; Figure S3C; Table S19).

Kaplan-Meier analysis by TMS tertiles revealed a monotonic survival gradient (Figure 5A): five-year RMST was 42.9 months in the low tertile (180 patients, 83 events), 48.8 months in the mid tertile (179 patients, 51 events), and 52.7 months in the high tertile (179 patients, 41 events), yielding an RMST difference of 9.8 months between the extreme tertiles (Table S8).

In LIHC, TMS was associated with OS in the univariable model (HR = 0.83; 95% CI 0.72-0.95; *p* = 0.008; *n* = 365, 130 events; concordance = 0.62), but this association did not meet the Bonferroni-corrected threshold and did not retain significance in the multivariable model (HR = 0.87; 95% CI 0.74-1.03; *p* = 0.11; *n* = 331, 111 events; concordance = 0.65; Figure 5B). Post hoc power analysis based on the observed multivariable effect size (HR = 0.87) and event count (111 events) estimated approximately 35% power to detect this effect at α = 0.05; detection of an HR of 0.87 at 80% power would require approximately 350 events, substantially exceeding the available LIHC sample.

Stage was the dominant predictor in LIHC (multivariable HR = 1.61; *p* = 1.0 × 10⁻⁵), while purity and TMB were non-significant. Stage-stratified Cox regression partially recovered the TMS association (HR = 0.83; *p* = 0.032), consistent with mild confounding between TMS and stage in the unstratified model. The proportional hazards assumption was satisfied for all covariates (all *p* > 0.21; Figure S3B; Table S3). Kaplan-Meier analysis showed an RMST difference of 12.6 months between extreme TMS tertiles (low: 34.3 months; high: 46.9 months; Figure S6A), numerically larger than in KIRC but accompanied by wider confidence intervals due to fewer events per tertile. The PFI sensitivity analysis in LIHC yielded a directionally concordant univariable HR of 0.88 (95% CI 0.78–1.01; p = 0.071; n = 366, 180 PFI events; Table S32; Figure S6B), consistent with the OS result but non-significant, mirroring the attenuation observed in the OS multivariable model (PFI multivariable HR = 0.96; p = 0.635; Table S33).

Shapley decomposition of the Royston-Sauerbrei pseudo-*R*² quantified the contribution of each covariate to explained survival variance (Figure 5C). In KIRC (full model *R*² = 0.189), stage dominated at 77.6% (Shapley value = 0.147; bootstrap 95% CI 0.071-0.208), followed by TMS at 16.3% (0.031; CI 0.007-0.066), purity at 4.6% (0.009; CI 0.003-0.022), and TMB at 1.5% (full-sample point estimate 0.003; bootstrap mean 0.022, median 0.019; percentile 95% CI 0.004-0.071; Table S5). The full-sample point estimate falls below the bootstrap CI lower bound because the Shapley permutation path for TMB encountered a near-zero marginal *R*² allocation in the full dataset, a boundary condition that is resolved in the majority of bootstrap resamples where covariate perturbation yields non-degenerate TMB contributions. In LIHC (*R*² = 0.081), stage contributed 73.9% (0.060; CI 0.021-0.114), TMS 14.6% (0.012; CI 0.001-0.043), TMB 11.2% (0.009; CI 0.000-0.042), and purity 0.3% (0.000; CI 0.000-0.016). Bootstrap convergence was 100% (500/500) for both cohorts (Table S5). TMS thus accounts for approximately one-sixth of explained survival variance in both KIRC and LIHC, conditional on stage, purity, and TMB.

### Cross-cohort directional concordance and clinical utility (C4)

Random-effects meta-analysis of univariable TMS hazard ratios from KIRC and LIHC yielded a pooled HR of 0.76 (95% CI 0.65-0.89; z = −3.44; p = 5.8 × 10⁻⁴), with consistent protective direction in both cohorts. Between-study heterogeneity was moderate (I² = 60.4%; Q = 2.52, p = 0.11; τ² = 0.007), reflecting the attenuation of effect size from KIRC (HR = 0.71) to LIHC (HR = 0.83). The C4 pass criterion (significant pooled HR with I² < 75%) was met (Tables S6, S7). The fixed-effect estimate was similar (pooled HR = 0.76; p = 3.2 × 10⁻⁸). With only two contributing cohorts, the I² estimate is not meaningfully interpretable, and the DerSimonian-Laird machinery is underpowered to characterize between-study variance. This result is therefore presented as a directional consistency assessment rather than a formal meta-analytic estimate of a common effect: both eligible cohorts show protective TMS associations, but the magnitude differs by tissue type, and whether additional cohorts would converge toward the KIRC or LIHC estimate cannot be determined from the present data. A forest plot of univariable HRs for all eight cohorts is provided in Figure S5B.

Category-free net reclassification improvement at the five-year horizon yielded a total NRI of 0.33 (*n* = 535, 149 events), with event NRI = 0.09 and non-event NRI = 0.24, indicating that TMS primarily improved risk stratification among patients who did not experience an event (correctly down-classifying low-risk patients; Table S17). Decision curve analysis at the five-year horizon demonstrated that the stage + TMS model produced higher net benefit than the stage-only model across a clinically relevant range of threshold probabilities (approximately 5%-55%; Figure 5F). The augmented model yielded higher net benefit than the stage-only, treat-all, and treat-none strategies over the moderate-risk threshold range. The three-year DCA showed a concordant pattern (Figure 5E; Table S18).

### Robustness and sensitivity analyses

The KIRC prognostic association was stable from k = 2 to k = 10 retained PCs (univariable HR range 0.68-0.82; all p < 0.007; Table S12; Figure S4A). LIHC and other cohort k-sensitivity results are reported in Supplementary Methods S10. Sequential removal of each organ from the GTEx reference confirmed that no single tissue dominated the PCA structure (subspace angle range 0.84°-20.6°; Table S15).

One thousand iterations of random gene-set PCA and Cox modeling yielded false positive rates of 4.1%-5.2% for all eight cohorts, with permuted HR distributions centered on 1.00 (Figure S4B; Table S13). No cohort exceeded the nominal 5% null rate. The TMS-survival association is therefore specific to the tissue-identity gene set and is not attributable to generic properties of PCA-derived distance metrics.

Purity-residualized TMS retained significant prognostic value in KIRC. Univariable Cox with residualized TMS yielded HR = 0.75 (95% CI 0.67-0.84; p = 1.1 × 10⁻⁶; n = 538, 175 events). In the multivariable model excluding purity (TMS_residual + stage + TMB), residualized TMS remained independently significant (HR = 0.70; 95% CI 0.59-0.84; p = 6.3 × 10⁻⁵; concordance = 0.76). Shapley decomposition of the purity-residualized model attributed 14.6% of explained variance to TMS_residual, 76.7% to stage, and 8.7% to TMB, closely mirroring the primary model allocation (16.3%, 77.6%, 4.6%, and 1.5%, respectively; Table S16) and demonstrating that the KIRC prognostic signal is not attributable to tumor purity confounding. LUAD residualization abolished the C2 effect (d = −0.005), confirming that its raw TMS-stage association was purity-driven.

Rank-normalized and median-dichotomized TMS transformations confirmed the KIRC association in both univariable and multivariable models (Tables S10, S11). Stage-stratified Cox regression preserved the TMS association in KIRC (HR = 0.60; p = 8.0 × 10⁻⁵) and attenuated it in LIHC (HR = 0.83; p = 0.032; Table S9). All sensitivity models were directionally concordant with the primary analysis. Rank-normalized TMS also yielded nominally significant univariable associations in two gate-excluded cohorts: LUAD (HR = 0.58; p = 0.028) and THCA (HR = 0.28; p = 0.011; Table S11). These signals were not promoted to primary inference because the pre-specified decision gate requires passage of both C1 and C2 before survival analysis; the gate criteria were locked prior to execution and were not modified *post hoc*. Cohorts that fail C2 lack evidence that TMS tracks the dedifferentiation axis in that tissue, and any univariable survival association in such cohorts may reflect confounding by unmeasured covariates, particularly tumor purity or immune infiltration, rather than genuine tissue memory attenuation. The gate therefore functions as a biological plausibility filter, not merely a statistical significance filter.

### Comparison with mRNAsi stemness index

To evaluate whether TMS captures prognostic information independent of transcriptional stemness (Supplementary Methods S8), per-sample one-class logistic regression (OCLR)-derived mRNAsi scores from Malta et al.^13^ were merged with the TMS cohort data for KIRC (n = 521 matched) and LIHC (n = 362 matched). In KIRC, TMS and mRNAsi were weakly correlated (Spearman ρ = 0.26, p = 1.2 × 10⁻⁹). Three nested Cox models were compared: Model A (TMS + stage + purity; C = 0.752), Model B (TMS + mRNAsi + stage + purity; C = 0.760, AIC = 1775.0), and Model C (mRNAsi + stage + purity; C = 0.741). In Model B, TMS retained independent significance (HR = 0.62; 95% CI 0.51-0.74; p = 2.3 × 10⁻⁷), while mRNAsi was non-significant (HR = 0.95; p = 0.55; Tables S41, S42). Model C, which excluded TMS, showed no prognostic contribution from mRNAsi (HR = 1.00; p = 0.96). The ΔAIC of −77.4 (Model B AIC = 1775.0; Model A AIC = 1852.4) is not interpretable as a model improvement metric because Models A and B are fitted to different sample sizes (n = 535 vs. n = 518) due to incomplete mRNAsi availability; AIC values computed on different samples are not directly comparable. The relevant comparison is between Models B and C, which share the same sample (n = 518): Model B (with TMS) achieves AIC = 1775.0 versus Model C (without TMS) AIC = 1794.9 (ΔAIC = −19.9), confirming TMS as the primary contributor to model fit. TMS therefore subsumes the prognostic information contained in mRNAsi for ccRCC.

In LIHC, TMS and mRNAsi were orthogonal (ρ = −0.01, p = 0.80; Table S40). Both achieved significance in Model B (TMS: HR = 0.83, p = 0.029; mRNAsi: HR = 1.28, p = 0.008), with ΔAIC = −6.2 relative to Model A, consistent with non-redundant prognostic contributions from the two indices in HCC (Figure 6A-C).

**Figure 6.**
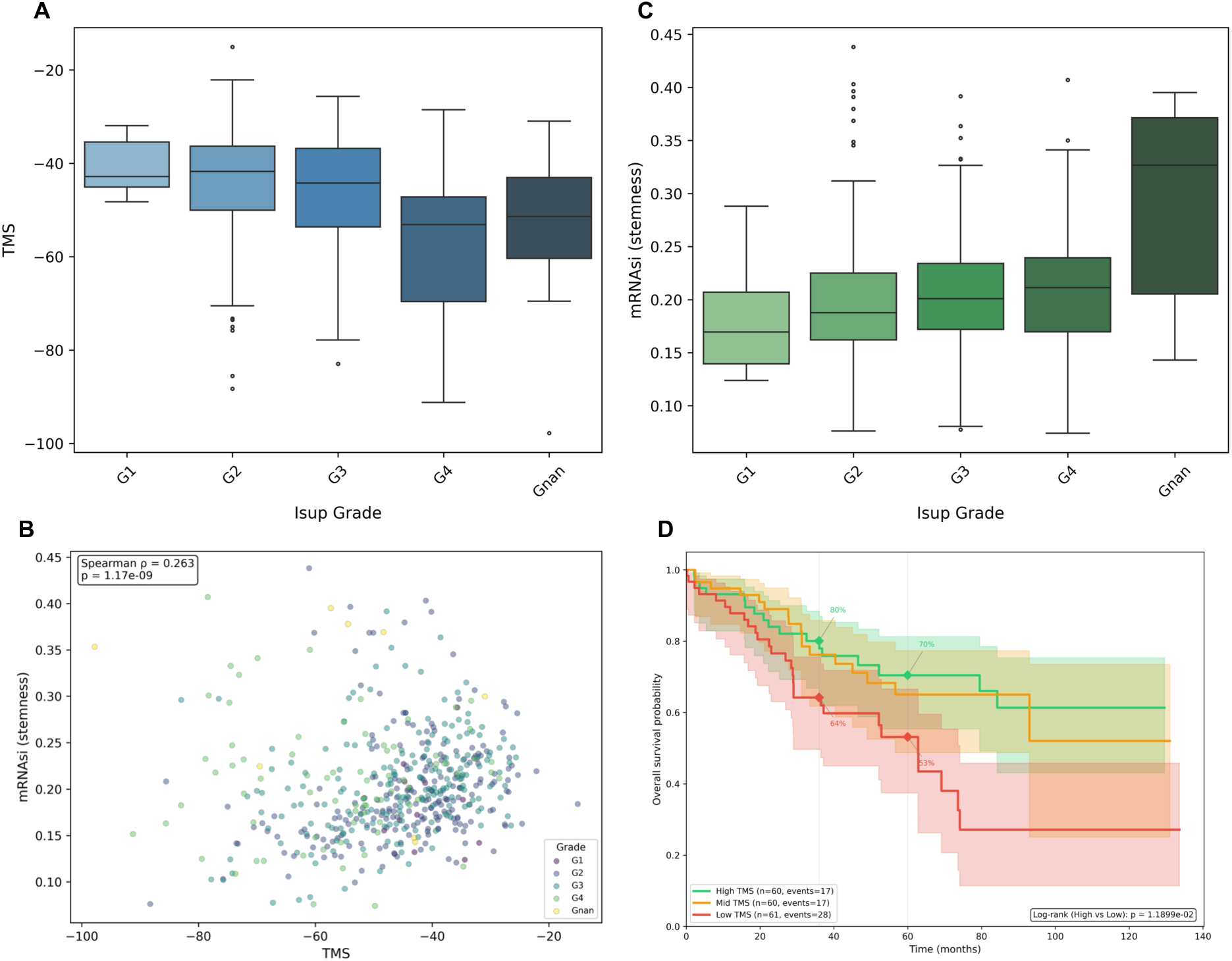
Stemness independence and clinical scenario. (A) TMS distribution by WHO/ISUP nuclear grade in TCGA-KIRC, demonstrating progressive decline with increasing grade. (B) Scatter plot of TMS versus OCLR-derived mRNAsi in KIRC, colored by ISUP grade (Spearman *ρ* = 0.26, *p* = 1.2 × 10⁻⁹). (C) mRNAsi distribution by ISUP grade, showing progressive increase with dediXerentiation. (D) Kaplan--Meier overall survival curves by TMS tertiles within intermediate-stage ccRCC (stages II--III; *n* = 181, 62 events). Annotated values denote three-year and five-year absolute survival probabilities per tertile.

### Clinical scenario: intermediate-stage ccRCC

Within AJCC stage II–III ccRCC (n = 181, 62 events; Supplementary Methods S9), TMS tertiles stratified five-year overall survival from 70.5% (high TMS) to 53.1% (low TMS), an absolute mortality difference of 17.3 percentage points (Table 4; Tables S43, S44; Figure 6D). The adjusted Cox model within this subset yielded HR = 0.56 (95% CI 0.43–0.73; *p* = 2.3 × 10⁻⁵), confirming that TMS discriminates prognosis beyond residual stage heterogeneity and purity within the intermediate-risk population.

**Table 4.** TMS-stratified absolute survival in intermediate-stage ccRCC (AJCC stage II–III; *n* = 181).

| TMS tertile | *n* | OS events | 3-year survival (95% CI) | 5-year survival (95% CI) | 3-year mortality | 5-year mortality |
| --- | --- | --- | --- | --- | --- | --- |
| High (best prognosis) | 60 | 17 | 80.1% (66.8–88.5) | 70.5% (55.4–81.3) | 20.0% | 29.5% |
| Mid | 60 | 17 | 76.2% (61.7–85.8) | 65.0% (48.7–77.3) | 23.8% | 35.0% |
| **Low (worst prognosis)** | **61** | **28** | **64.2% (49.6–75.6)** | **53.1% (37.4–66.6)** | **35.8%** | **46.9%** |
| Absolute risk difference (Low vs High) |  |  | +15.8 pp | +17.3 pp |  |  |
| Cox regression within subset |  |  |  |  |  |  |
| Univariable (TMS per SD) |  |  | HR = 0.78 (0.62–0.99) | *P* = 0.039 | C = 0.563 |  |
| Adjusted (TMS + stage + purity) |  |  | **HR = 0.56 (0.43–0.73)** | ***P* = 2.3 × 10<sup>−5</sup>** | C = 0.680 |  |

TMS tertile boundaries computed within the intermediate-stage subset. Survival probabilities from Kaplan–Meier estimates with 95% Greenwood confidence intervals. pp, percentage points.

### External validation

The locked PCA reference was applied to 261 CPTAC-ccRCC RNA-seq samples without re-estimation (Table S22). HVG overlap was 97.0%. C1 accuracy was 95.0%, closely matching the TCGA-KIRC discovery value (95.9%). TMS distribution parameters were concordant between validation (mean = −48.9, SD = 12.9) and discovery (mean = −46.5, SD = 12.9; Figure 7B), with a Kolmogorov-Smirnov (KS) D = 0.09 (*p* = 0.08; Table S26) and a mean shift of −2.3 (0.18 SD units). Projection of CPTAC tumors onto the locked PCA axes confirmed co-localization with TCGA-KIRC tumors near the kidney centroid (Figure 7A).

**Figure 7.**
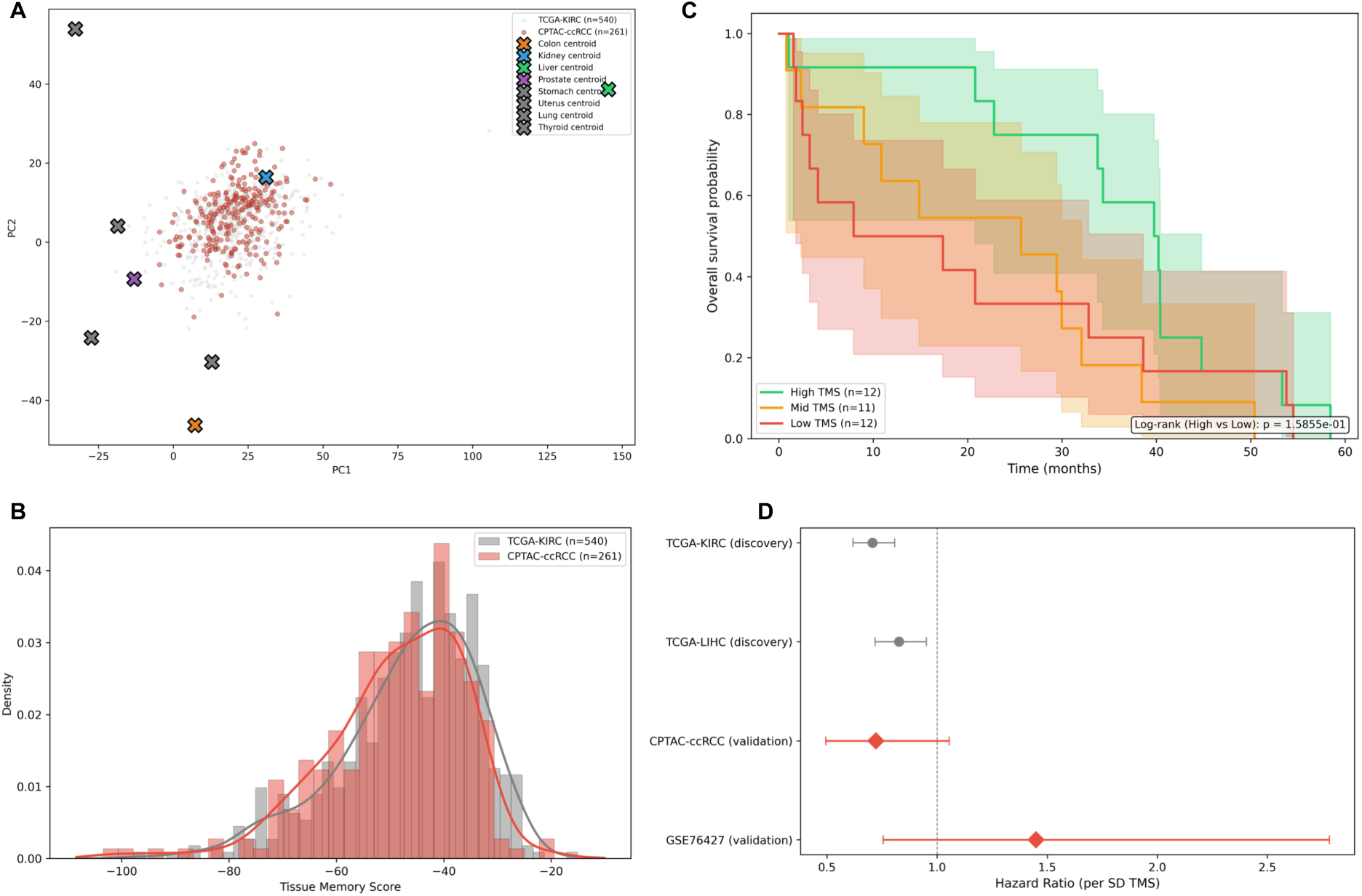
External validation. (A) CPTAC-ccRCC tumors (*n* = 261) projected onto the locked GTEx PCA axes alongside TCGA-KIRC tumors (grey), confirming co-localization near the kidney centroid. (B) TMS density distributions for TCGA-KIRC (discovery) and CPTAC-ccRCC (validation), showing distributional concordance (KS *D* = 0.09). (C) Kaplan–Meier overall survival curves by TMS tertiles in CPTAC-ccRCC (*n* = 35 with survival annotation). (D) Forest plot of univariable TMS hazard ratios for the two discovery cohorts (TCGA-KIRC, TCGA-LIHC) and two validation cohorts (CPTAC-ccRCC, GSE76427), with pooled random-eXects estimate. Diamonds denote validation cohorts.

Among the 35 patients with survival annotation (35 events, 0 censored), univariable Cox regression yielded HR = 0.72 (95% CI 0.49-1.06; *p* = 0.093; Table S23), with concordant HR direction (R1: pass) and overlapping confidence intervals with the discovery HR (R3: pass) but failing the significance criterion (R2: fail). The absence of censored observations reflects the structure of the CPTAC-ccRCC clinical annotation, in which overall survival data were available exclusively for deceased patients at the time of data freeze. This restriction conditions the Cox estimate on the occurrence of death and may attenuate the estimated hazard ratio toward the null; the resulting HR should therefore be interpreted as a lower bound on the true effect magnitude. The replication verdict was therefore PARTIAL REPLICATION (Table S25), consistent with the small survival-annotated sample size limiting statistical power. The CPTAC-ccRCC validation should therefore be interpreted as a distributional and directional concordance assessment of the locked reference, not as a powered replication of the survival association; the decedent-only sampling frame, absence of censored observations, and 35-patient sample size preclude inference on effect magnitude.

Kaplan-Meier analysis by TMS tertiles showed a concordant separation pattern (Figure 7C; Table S24). Fixed-effect meta-analysis pooling TCGA-KIRC and CPTAC-ccRCC yielded a combined HR of 0.71 (*p* = 7.6 × 10⁻⁸; *I*² = 0%), confirming complete absence of heterogeneity between the two ccRCC cohorts (Figure 7D). The statistical power of the pooled estimate derives almost entirely from the TCGA-KIRC discovery cohort; the CPTAC contribution constrains the heterogeneity estimate but does not materially alter the point estimate or its precision.

GSE76427 comprised 115 HCC patients profiled on the Illumina HumanHT-12 v4 microarray. HVG overlap was 88.2%, below the 90% target specified in the design document. TMS distributions were dramatically shifted (validation mean = −140.2, SD = 6.9; discovery mean = −37.2, SD = 23.3; KS *D* = 0.99, *p* < 10⁻¹³⁴), reflecting a fundamental platform incompatibility: microarray intensity scaling compresses the dynamic range of the PCA projection, and the lower HVG overlap further distorts the distance computation. C1 accuracy in GSE76427 was 28.7%, far below the TCGA-LIHC value of 96.0%. The univariable Cox HR was 1.45 (95% CI 0.76-2.78; *p* = 0.26), reversed in direction from the discovery HR. The replication verdict was NON-REPLICATION. Given the platform incompatibility (microarray vs. RNA-seq), distributional shift exceeding 4.4 SD units, and HVG overlap below 90%, this non-replication is attributed to technical rather than biological discordance (Figure S8). An RNA-seq validation cohort would be required to evaluate LIHC TMS replication adequately. LIRI-JP (ICGC) was identified as the most suitable candidate; however, the ICGC Data Portal was decommissioned in June 2024, and access to the archived expression matrices and matched clinical endpoints requires authorization through the ICGC 25K Data Access process, which was not executed within the scope of the present study.

## Discussion

This study establishes a quantitative framework for measuring tissue-of-origin transcriptional memory in solid tumors and evaluates its prognostic relevance in a pre-specified pan-cancer design. Three findings emerge from this analysis. First, tissue identity retention is a near-universal property: six of eight tumor types project closer to their matched healthy tissue centroid than to any alternative, with classification accuracies of 80%-100% against a 12.5% chance baseline. Second, the prognostic informativeness of tissue memory is tissue-dependent: TMS carries independent survival information in ccRCC (multivariable HR = 0.59; Shapley contribution 16.3%) but not, after covariate adjustment, in HCC. Third, the conditions under which tissue memory measurement fails constitute informative boundary conditions, delineating the dependence of the framework on faithful representation of the tumor-origin cellular compartment in the reference atlas.

### Tissue identity retention and boundary conditions

The C1 result corroborates and extends the observation by Hoadley *et al.*^1^ that cell-of-origin patterns dominate the molecular classification of TCGA tumors. Where Hoadley *et al.* demonstrated this principle through unsupervised clustering of multi-platform data from 33 cancer types, the present study operationalizes tissue identity as a continuous, per-sample distance metric referenced to an external healthy atlas. The two approaches address different levels of the same question: the Hoadley classification captures inter-tumor-type separation, while TMS quantifies the degree of identity retention within a single tumor type and links it to clinical covariates.

The near-perfect C1 accuracy in PRAD (100%) and THCA (99.2%) reflects the transcriptional distinctiveness of prostatic and thyroid epithelium, driven by lineage-defining programs such as androgen receptor signaling and thyroglobulin synthesis, respectively. The high but imperfect accuracy in KIRC (95.9%) and LIHC (96.0%) is consistent with the broader transcriptional repertoire of renal proximal tubular and hepatocytic programs, which share metabolic gene expression with other parenchymal organs. LUAD’s lower accuracy (80.2%) is expected given the transcriptional heterogeneity of lung adenocarcinoma and the mixed cellularity of normal lung tissue, which encompasses alveolar, bronchial, and stromal compartments in proportions that may not match the distal airway progenitor origin of LUAD^3^.

The UCEC and STAD C1 failures are not methodological artifacts but biological boundary conditions of the framework. UCEC’s near-complete failure (1.3% accuracy) originates from a fundamental mismatch between the GTEx Uterus reference, which is dominated by myometrial smooth muscle, and the glandular epithelial compartment from which endometrioid carcinomas arise. STAD’s 43.7% accuracy reflects the transcriptional overlap between gastric and colonic mucosa, a consequence of shared embryonic endodermal origin that is sufficient to blur organ-level centroid discrimination in the joint PCA space. These failures highlight a general principle: the informativeness of atlas-based tissue identity quantification is bounded by the compartmental fidelity of the reference. Bulk RNA-seq atlases sample tissue-level transcriptomes, not cell-type-specific programs, and how well the dominant transcriptional signal matches the carcinoma-origin cell type differs by organ. Single-cell atlases with cell-type deconvolution could address this limitation, at the cost of the format compatibility that bulk-to-bulk projection affords.

### Prognostic relevance in ccRCC

The restriction of confirmed C3 results to KIRC is not unexpected given the biology of ccRCC. Clear cell renal cell carcinoma arises from proximal tubular epithelium, and the Fuhrman/WHO-ISUP nuclear grading system^4,5^ captures nuclear features that correlate with progressive loss of proximal tubular epithelial differentiation. The TMS-stage association in KIRC (d = 0.42) and the independent survival association (multivariable HR = 0.59) demonstrate that molecular retention of proximal tubular transcriptional identity, as captured by distance to the GTEx Kidney-Cortex centroid, adds prognostic information to stage, purity, and TMB. The Shapley-attributed 16.3% of explained survival variance positions TMS as the second-most informative predictor in the KIRC multivariable model, behind stage (77.6%) but substantially exceeding purity (4.6%) and TMB (1.5%).

The TMS-survival association held in every pre-specified sensitivity analysis (purity residualization, rank transformation, median dichotomization, stage stratification, and time-varying coefficient models), with no violation of the proportional hazards assumption and a false positive rate of 5.1% in the random-gene negative control (Results, Section 4). The consistency of the association with progression-free interval (PFI multivariable HR = 0.58; *p* = 4.7 × 10⁻⁵; Supplementary Methods S6; Figure S7; Tables S32-S35) supports the conclusion that TMS reflects tumor-intrinsic biology and not general frailty or non-cancer mortality. The PFI model for KIRC detected a PH violation for Stage (*ρ* = 0.273, *p* = 0.008; Table S34); the TMS coefficient itself satisfied the assumption (*ρ* = −0.129, *p* = 0.217).

Per-axis decomposition revealed that the prognostic signal localizes to PC2 (adjusted HR = 0.73; *p* = 0.012; Figure 5D), an axis enriched for renal solute transporter programs (Reactome: Multifunctional Anion Exchangers; Figure S5A; Tables S20, S21). PC1 and PC3 carried no independent survival information (Supplementary Methods S7; Tables S36-S39). This concentration of prognostic content on a single transcriptional axis implies that tissue memory loss in ccRCC is not a diffuse, genome-wide process but a specific erosion of tubular transport identity, consistent with the known histological progression from well-differentiated clear cell morphology toward sarcomatoid dedifferentiation.

These results translate into clinically interpretable risk stratification. Within intermediate-stage ccRCC (stages II-III; *n* = 181), TMS tertiles separated five-year mortality by 17.3 absolute percentage points (low TMS: 46.9%; high TMS: 29.5%), with an adjusted HR of 0.56 (*p* = 2.3 × 10⁻⁵). This absolute risk difference is of a magnitude that may inform post-nephrectomy surveillance intensity or adjuvant therapy candidacy in a multidisciplinary tumor board setting, although comparison with treatment effects from randomized trials is precluded by differences in endpoint definition, time horizon, and study design. The NRI of 0.33 at five years and the positive net benefit across the 5%-40% threshold range on decision curve analysis provide complementary evidence that TMS adds decision-relevant information to stage-based risk models.

The two most widely adopted post-nephrectomy risk stratification instruments for ccRCC are the UCLA Integrated Staging System (UISS)^30^ and the Leibovich score^31^. Neither system incorporates molecular data. TMS occupies a complementary axis: it quantifies the molecular residual of differentiated transcriptional identity, a property related to histological grade but not reducible to it, one of the shared inputs to both scoring systems. The weak TMS-stage correlation in KIRC (Spearman *ρ* = −0.20) and the retained prognostic significance of TMS after stage adjustment (multivariable HR = 0.59; *p* = 6.6 × 10⁻⁵) confirm that TMS captures survival-relevant information orthogonal to the staging and grading components common to UISS and Leibovich. Direct quantification of incremental discrimination relative to these instruments was not feasible because ECOG performance status and tumor necrosis status are not uniformly recorded in the TCGA-KIRC clinical data; an institutional cohort with complete UISS and Leibovich annotation alongside matched RNA-seq would be required.

### The purity confound and its resolution

The TMS-purity correlation in KIRC (*ρ* = 0.71) is the primary methodological obstacle to interpreting the prognostic association. Tumors with low purity (high stromal and immune admixture) contain normal tissue RNA that mechanistically inflates TMS. Three lines of evidence, detailed in Results, argue that the KIRC prognostic signal is not attributable to purity: purity-residualized TMS retains both univariable and multivariable significance with near-identical Shapley variance allocation; Shapley decomposition allocates only 4.6% of survival variance to purity versus 16.3% to TMS; and the multivariable model simultaneously includes purity as a covariate with VIF well below conventional collinearity thresholds. The sign inversion of purity in the multivariable model (univariable HR = 0.45, protective; multivariable HR = 4.83, adverse) is a Simpson’s paradox: after conditioning on TMS and stage, the residual purity variation indexes a distinct biological property, possibly the association between high tumor cellularity and aggressive clear cell histological features, that carries an independent adverse prognostic association.

The converse case is informative: in LUAD, where TMS-purity correlation was *ρ* = −0.63, residualization abolished both the C2 stage association (d = −0.005) and any candidate prognostic signal, confirming that LUAD’s raw TMS was a purity proxy. The differential behavior of KIRC (TMS retains prognostic value after residualization) and LUAD (TMS does not) validates the residualization protocol as an effective discriminator of genuine tissue memory signal from purity artifact.

### LIHC, external validation, and relationship to existing frameworks

LIHC presents a biologically coherent but prognostically non-definitive picture. The univariable TMS-survival association (HR = 0.83; *p* = 0.008) did not survive Bonferroni correction and was attenuated to non-significance in the multivariable model (HR = 0.87; *p* = 0.11). The discrepancy between the Shapley contribution (14.6%, comparable to KIRC) and the multivariable p-value reflects the lower total explained variance in LIHC (*R*² = 0.081 vs. 0.189 in KIRC): TMS accounts for a similar relative share of a smaller total, yielding insufficient absolute effect magnitude for detection at the Bonferroni-corrected threshold. Formal power estimation confirms that the LIHC multivariable model (111 events, observed HR = 0.87) operated at approximately 35% power; confirmation of the LIHC TMS association at conventional thresholds would require a cohort with approximately 350 OS events, a sample size achievable through multi-institutional collaboration or meta-analytic pooling of RNA-seq HCC cohorts.

The Edmondson-Steiner grading system^6^ captures dedifferentiation in HCC along a morphological axis, and the stage-based C2 association (d = 0.50) confirms that TMS tracks this axis. The failure to confirm independent prognostic value is plausibly explained by the fact that dedifferentiation in HCC is prognostically subordinate to vascular invasion and portal hypertension severity, neither of which is captured in the present covariate set.

The CPTAC-ccRCC partial replication (HR = 0.72; *p* = 0.093; direction concordant, CIs overlapping) demonstrates that the locked reference transfers to an independent RNA-seq cohort without re-estimation. The distributional concordance (KS *D* = 0.09; mean shift 0.18 SD units) and preserved C1 accuracy (95.0% vs. 95.9%) confirm platform stability within the RNA-seq modality. The failure to reach statistical significance is attributable to the survival-annotated sample size (*n* = 35), which yields approximately 40% power to detect an HR of 0.72 at α = 0.05. Fixed-effect pooling of the two ccRCC cohorts produced HR = 0.71 (*p* = 7.6 × 10⁻⁸; *I*² = 0%), confirming complete absence of heterogeneity. The GSE76427 non-replication is entirely attributable to platform incompatibility: the 4.4 SD distributional shift, 88.2% HVG overlap, and reversed HR direction demonstrate that microarray intensity scaling is fundamentally incompatible with an RNA-seq-derived PCA reference. RNA-seq HCC cohorts such as LIRI-JP would be required for a valid LIHC replication attempt.

TMS is conceptually distinct from machine-learning-derived stemness indices such as mRNAsi^13^. The mRNAsi score measures resemblance to pluripotent stem cells along a universal stemness axis, while TMS measures retention of a specific differentiated program by projection onto tissue-specific healthy reference axes. The two metrics capture opposing directions of the same phenotypic continuum: mRNAsi quantifies what tumors gain (pluripotency-associated transcription), while TMS quantifies what tumors lose (lineage-specific transcription). Direct comparison confirmed that TMS and mRNAsi are weakly correlated in ccRCC (*ρ* = 0.26) and orthogonal in LIHC (*ρ* = −0.01). Variance inflation factors in the joint four-covariate Cox model confirmed the absence of problematic collinearity (VIF_-./_ = 2.21, 𝑉𝐼𝐹_012345_ = 1.16 in KIRC; 𝑉𝐼𝐹_6./_= 1.19, VIF_712345_ = 1.05 in LIHC; all below the conventional threshold of 5).

In the joint multivariable model, TMS subsumed the prognostic content of mRNAsi in ccRCC (TMS HR = 0.61, *p* = 2.3 × 10⁻⁷; mRNAsi HR = 0.95, *p* = 0.55), while both indices contributed non-redundantly in HCC (TMS HR = 0.83, p = 0.029; mRNAsi HR = 1.28, p = 0.008). These tissue-dependent patterns may reflect the differential relevance of stemness-associated transcription to dedifferentiation trajectories in renal versus hepatic lineages. Similarly, Ben-Porath et al.^14^ established the transcriptional link between stemness and poor outcome in breast cancer. The present work generalizes this concept by replacing a fixed gene signature with a data-driven, atlas-referenced distance metric that adapts to each tissue lineage, because the PCA loadings that define the score differ by tissue.

### Strengths and limitations

The pre-specified decision gate design, with locked analytic thresholds and no post-hoc modifications, distinguishes this study from exploratory pan-cancer biomarker analyses in which positive results may be selected from a larger space of tested models. The decision gate summary reports both passage and failure, documenting all analytic degrees of freedom. Shapley-based variance attribution quantifies the unique survival variance contributed by TMS in the presence of correlated covariates, and the combination with NRI and DCA provides a multi-dimensional assessment of prognostic value at actionable thresholds. The negative control analysis (1,000 random gene-set iterations) provides empirical confirmation that the observed associations are specific to the tissue-identity gene set.

Six limitations apply. First, the C4 assessment includes only two cohorts, precluding formal meta-analytic inference. The *I*² estimate is not meaningfully interpretable with *n* = 2 studies, and the C4 result is framed as a directional consistency assessment. Second, the TMS framework operates on bulk RNA-seq, which captures tissue-level transcriptomes without cell-type resolution. Although purity residualization effectively deconfounds TMS in KIRC, the framework cannot distinguish genuine epithelial-intrinsic tissue memory from stromal contamination at the single-cell level. Single-cell RNA-seq with cell-type deconvolution (e.g., CIBERSORTx) could decompose bulk TMS into epithelial and stromal components. Third, the GTEx reference comprises post-mortem tissue from adult donors, and agonal gene expression changes may alter the reference PCA axes. The bootstrap stability analysis (mean angle 3.30°) mitigates concern about donor-specific outliers but does not specifically address agonal contamination. Fourth, the external validation for KIRC is underpowered (*n* = 35 survival-annotated patients), and the LIHC validation was invalidated by microarray platform incompatibility, and the most suitable RNA-seq alternative (LIRI-JP) was not included because the ICGC Data Portal was decommissioned prior to study execution, with residual access restricted to authorized users. Fifth, overall survival is a composite endpoint influenced by salvage therapies and non-cancer mortality; the PFI sensitivity analysis argues against a non-tumor-driven mechanism but does not fully exclude confounding by general frailty. Sixth, TMS collapses the multi-dimensional structure of tissue identity loss into a scalar. Per-axis decomposition in KIRC demonstrated prognostic concentration on PC2, but whether this generalizes to other tumor types remains to be determined. Alternative distance metrics (e.g., Mahalanobis distance or per-axis weighted composites) could capture axis-dependent prognostic heterogeneity more faithfully.

## Conclusion

This study establishes tissue-of-origin transcriptional memory as a quantifiable property of solid tumors and demonstrates, through a pre-specified multi-stage analytic design with locked decision gates, that the degree of this retention carries independent prognostic value in ccRCC. Six of eight tumor types retained transcriptional identity of their tissue of origin; the prognostic informativeness of this retention proved tissue-dependent. In LIHC, TMS exhibited a directionally concordant but statistically underpowered association that warrants adequately powered confirmation in an independent RNA-seq HCC cohort.

The mechanistic localization of the prognostic signal to PC2 (renal solute transporter programs) implies that tissue memory loss in ccRCC reflects a specific erosion of tubular transport identity, not a diffuse transcriptomic decay. The 17 percentage-point five-year mortality separation between TMS tertiles in intermediate-stage ccRCC positions TMS as a candidate molecular input for post-nephrectomy risk stratification, complementary to existing clinicopathological instruments (UISS, Leibovich) that do not incorporate transcriptomic data. The C1 failures in STAD and UCEC identify reference atlas composition as the principal constraint on framework applicability, establishing that bulk tissue references must faithfully represent the cellular compartment from which carcinomas arise.

Three questions remain open: single-cell decomposition of epithelial-intrinsic tissue memory to resolve the bulk RNA-seq resolution limitation, adequately powered RNA-seq validation in both ccRCC and HCC, and integration with established clinicopathological instruments to quantify incremental discrimination in institutional cohorts with complete staging and grading annotation.

