## Supplementary for "Tissue Memory Score measures transcriptional retention in solid tumors and predicts survival in clear cell renal cell carcinoma"

### Table of Contents

**Supplementary Methods S1** — Tumor purity computation
**Supplementary Methods S2** — Histological grade extraction
**Supplementary Methods S3** — Covariate correlations and purity residualization
**Supplementary Methods S4** — Normal tissue control
**Supplementary Methods S5** — Proportional hazards diagnostics
**Supplementary Methods S6** — Progression-free interval sensitivity analysis
**Supplementary Methods S7** — Per-principal-component prognostic decomposition
**Supplementary Methods S8** — Stemness comparison
**Supplementary Methods S9** — Clinical scenario: intermediate-stage ccRCC
**Supplementary Methods S10** — Robustness and sensitivity analyses
**Supplementary Methods S11** — External validation details

**Supplementary Figure S1** — All eight TCGA tumor cohort projections onto the locked GTEx V8 PCA reference
**Supplementary Figure S2** — Covariate correlations and grade-based C2 testing
**Supplementary Figure S3** — Proportional hazards diagnostics
**Supplementary Figure S4** — Robustness analyses
**Supplementary Figure S5** — Extended robustness and pan-cancer forest
**Supplementary Figure S6** — LIHC detailed results (exploratory)
**Supplementary Figure S7** — Progression-free interval endpoint sensitivity
**Supplementary Figure S8** — GSE76427 validation failure (LIHC)

**Supplementary Table S1** — Univariable Cox models for TMS (all cohorts)
**Supplementary Table S2** — Multivariable Cox models (KIRC, LIHC)
**Supplementary Table S3** — Proportional hazards assumption testing
**Supplementary Table S4** — Variance inflation factor diagnostics
**Supplementary Table S5** — Shapley decomposition bootstrap CIs
**Supplementary Table S6** — Meta-analysis pooled HR
**Supplementary Table S7** — Meta-analysis per-cohort inputs
**Supplementary Table S8** — RMST by TMS tertiles
**Supplementary Table S9** — Stage-stratified Cox regression
**Supplementary Table S10** — Median-dichotomized TMS Cox models
**Supplementary Table S11** — Rank-normalised TMS Cox models
**Supplementary Table S12** — PCA dimensionality sensitivity (k sweep)
**Supplementary Table S13** — Random-gene negative control
**Supplementary Table S14** — GTEx subsampling stability
**Supplementary Table S15** — Leave-one-organ-out analysis
**Supplementary Table S16** — Purity-residualized TMS Cox models (KIRC)
**Supplementary Table S17** — Net reclassification improvement (KIRC)
**Supplementary Table S18** — Decision curve analysis (KIRC, 3yr and 5yr)
**Supplementary Table S19** — Time-varying coefficient Cox models (KIRC)

**Supplementary Table S20** — PCA loading enrichment (Enrichr)

**Supplementary Table S21** — Top PCA loading genes per component

**Supplementary Table S22** — Validation cohort summary

**Supplementary Table S23** — Validation univariable Cox models

**Supplementary Table S24** — Validation multivariable Cox models

**Supplementary Table S25** — Validation replication assessment

**Supplementary Table S26** — Validation distributional concordance

**Supplementary Table S27** — TMS-covariate correlations (all cohorts)

**Supplementary Table S28** — Normal tissue control

**Supplementary Table S29** — Purity residualization diagnostics

**Supplementary Table S30** — Stage-based C2 attenuation effects

**Supplementary Table S31** — Grade-based C2 associations

**Supplementary Table S32** — PFI univariable Cox models

**Supplementary Table S33** — PFI multivariable Cox models

**Supplementary Table S34** — PFI proportional hazards assumption

**Supplementary Table S35** — OS vs PFI endpoint concordance

**Supplementary Table S36** — Per-PC univariable Cox models

**Supplementary Table S37** — Per-PC multivariable Cox models

**Supplementary Table S38** — PC axis inter-variable correlations

**Supplementary Table S39** — Prognostic axis enrichment summary

**Supplementary Table S40** — TMS-mRNAsi correlations

**Supplementary Table S41** — Joint Cox models (TMS + mRNAsi)

**Supplementary Table S42** — Stemness model comparison

**Supplementary Table S43** — Intermediate-stage absolute risk

**Supplementary Table S44** — Intermediate-stage Cox regression

#### Supplementary Methods

##### S1. Tumor purity computation

ESTIMATE tumor purity scores were computed locally from TCGA expression matrices using single-sample gene set enrichment analysis (ssGSEA). The algorithm follows the formulation of Barbie et al. [25] and Yoshihara et al. [26]. For each sample, genes were ranked by descending expression magnitude. Cumulative enrichment scores for the stromal signature (141 genes) and immune signature (141 genes) were computed by weighting hit positions by the absolute expression magnitude raised to the power $\alpha = 0.25$, with normalization of the cumulative hit and miss distributions to yield per-signature enrichment scores. The ESTIMATE score was defined as the sum of the stromal and immune scores and converted to tumor purity via the published cosine transformation:

$$\text{Purity} = \cos(0.6049872018 + 0.0001467884 \times \text{ESTIMATE score})$$

An earlier implementation using positional index weighting (1, 2, 3, …) rather than expression-magnitude weighting produced near-constant purity scores (SD < 0.01 within cohorts). This error was identified by inspecting the output distribution and corrected prior to all downstream analyses. The corrected implementation was verified by comparing output distributions against published ESTIMATE purity values for TCGA cohorts.

##### S2. Histological grade extraction

Histological grade data were extracted from GDC clinical supplements for cohorts with established grading systems. For TCGA-PRAD, ISUP Grade Group was derived from primary and secondary Gleason patterns: Gleason sum 6 (3+3) = Grade Group 1, Gleason 3+4 = Grade Group 2, Gleason 4+3 = Grade Group 3, Gleason 8 = Grade Group 4, and Gleason 9–10 = Grade Group 5 [8,9]. For TCGA-STAD, WHO histological grade (G1, G2, G3) and Lauren classification (intestinal, mixed, diffuse) were extracted [10]. For TCGA-UCEC, FIGO grade (1–3) was extracted from the clinical supplement [11]. Grade data were unavailable for COAD and not applicable to LIHC, LUAD, and THCA, which lack consensus histological grading systems amenable to ordinal encoding. For TCGA-KIRC, WHO/ISUP nuclear grade (G1–G4) was extracted from GDC clinical supplements and used in the stemness comparison analysis (Section S8).

##### S3. Covariate correlations and purity residualization

Spearman rank correlations between TMS and each covariate (ESTIMATE purity, $\log_{10}$ TMB, ordinal stage, immune score) were computed per cohort. Two cohorts exhibited |rho_TMS-purity| > 0.5: KIRC (Spearman $\rho = 0.71$; $p = 7.5 \times 10^{-84}$) and LUAD ($\rho = -0.63$; $p = 1.5 \times 10^{-61}$). For these cohorts, purity-residualized TMS was computed as the residual from an ordinary least squares regression of TMS on ESTIMATE purity. Post-residualization, the TMS-purity correlation was reduced to $\rho = 0.02$ in KIRC and $\rho = -0.02$ in LUAD, rendering the residualized score effectively orthogonal to purity. Residualized TMS was carried forward into sensitivity survival analyses for KIRC. In LUAD, residualization abolished the stage-based C2 effect (Cohen $d = -0.005$; $p = 0.96$), confirming that the raw TMS-stage association was entirely purity-driven. TMS-TMB correlations were weak in all cohorts ($|\rho| \leq 0.23$). TMS-stage correlations were modest, with the strongest in LIHC ($\rho = -0.27$) and KIRC ($\rho = -0.20$).

##### S4. Normal tissue control

Where TCGA-matched normal tissue samples were available, TMS was computed for these normals using the same locked PCA projection. The expected direction (normal TMS > tumor TMS) was evaluated by Cohen d and Welch t-test as an internal validation of the tissue memory construct. TCGA-matched normals exhibited higher TMS than paired primary tumors in seven of eight cohorts. The largest effects were observed in LUAD (Cohen d = 2.50; p = $7.5 \times 10^{-56}$), KIRC (d = 1.69; p = $1.1 \times 10^{-26}$), THCA (d = 1.21; p = $6.7 \times 10^{-12}$), and LIHC (d = 0.96; p = $1.9 \times 10^{-28}$). UCEC showed a particularly large effect (d = 4.11; p = $2.2 \times 10^{-15}$), consistent with the dramatic separation between myometrial normals and endometrial tumors in PCA space. STAD was the sole exception: normal tissue TMS was lower than tumor TMS (d = −0.64; p = 0.025), an inversion likely reflecting the heterogeneous cellular composition of GTEx Stomach relative to the mucosal epithelial origin of gastric adenocarcinoma.

#### S5. Proportional hazards diagnostics

The proportional hazards assumption was tested for each covariate by computing the Spearman correlation between scaled Schoenfeld residuals and ranked event times. In KIRC, no covariate violated the assumption at $\alpha = 0.05$ (TMS: $\rho = -0.19$, p = 0.064; stage: p = 0.28; purity: p = 0.31; TMB: p = 0.23). As a pre-specified sensitivity, time-varying coefficient models were fitted for KIRC using piecewise Cox regression with cutpoints at median follow-up (1,133 days), three years (1,095 days), and five years (1,826 days). In LIHC, all covariates satisfied the assumption (all p > 0.21). The TMS hazard ratio was stable across early and late follow-up periods: at the median cutpoint, early-period HR = 0.73 (p = 0.005) and late-period HR = 0.67 (p = 0.016). At the three-year cutpoint, early HR = 0.71 (p = 0.002) and late HR = 0.73 (p = 0.027). At the five-year cutpoint, early HR = 0.7116 (p = 0.0007) and late HR = 0.6932 (p = 0.155). The consistency of effect estimates across time intervals confirms that the proportional hazards assumption is not materially violated and that the TMS-survival association is not driven by a specific follow-up epoch.

#### S6. Progression-free interval sensitivity analysis

To evaluate whether TMS captures tumor-intrinsic biology rather than general frailty or non-cancer mortality, the primary survival models were refitted using progression-free interval (PFI) from the TCGA Clinical Data Resource [27] as an alternative endpoint. PFI was available for both gate-passing cohorts: KIRC (536 patients, 159 PFI events) and LIHC (366 patients, 180 PFI events).

Univariable and multivariable Cox models were fitted with the same covariate specification as the OS models (TMS_z, ordinal stage, ESTIMATE purity, $\log_{10}$ TMB). Proportional hazards diagnostics were repeated for the PFI models. In KIRC, Stage exhibited a time-varying effect ($\rho = 0.273$, p = 0.0078; Table S34), while TMS ($\rho = -0.129$, p = 0.217), purity, and TMB satisfied the assumption. Because the violation is confined to the Stage covariate and the TMS coefficient is unaffected, the PFI hazard ratio for TMS remains interpretable under the proportional hazards formulation; however, the Stage coefficient should be understood as an averaged effect over follow-up rather than a constant instantaneous hazard ratio. In LIHC, all covariates satisfied the assumption (all p > 0.42). In KIRC, the PFI multivariable HR for TMS was 0.58 (95% CI 0.45–0.76; p = $4.7 \times 10^{-5}$), closely matching the OS estimate (HR = 0.59). In LIHC, the PFI univariable HR was 0.89 (p = 0.07), directionally concordant with the OS result but non-significant. Endpoint concordance was assessed by comparing the direction of the TMS hazard ratio, overlap of 95% confidence intervals, and concordance of statistical significance between OS and PFI models for each cohort. The concordance of HR direction between OS and PFI endpoints, with overlapping confidence intervals and comparable effect magnitudes, argues against a non-tumor-driven mechanism.

#### S7. Per-principal-component prognostic decomposition

To identify which transcriptional program(s) carry the prognostic content of TMS, the composite Euclidean distance was decomposed into signed per-axis displacements. For each tumor sample i and principal component k (k = 1, 2, 3), the signed displacement was computed as:

dik=vik-ckmatched

where vik is the tumor's projection on PCk and ckmatched is the matched centroid coordinate on axis k. Positive values denote displacement in the direction of increasing PC score; negative values denote displacement in the opposite direction. Each displacement was standardized to zero mean and unit variance within its cohort.

Univariable Cox models (OS~dPCk) were fitted per axis per cohort using both signed and absolute-value formulations. A joint multivariable model including all three axis displacements was fitted to assess whether prognostic information was concentrated on a single axis or distributed. An adjusted model additionally included stage and ESTIMATE purity. Spearman correlations among axis displacements, TMS, purity, and stage were computed to characterize the collinearity structure. Prognostically significant axes ($p < 0.05$, signed variant) were annotated with the top enrichment terms from the PCA loading enrichment table (WP5b), linking the active axis to specific biological programs.

In KIRC, the prognostic signal localized to PC2 (signed displacement univariable HR = 0.73; $p = 3.1 \times 10^{-4}$), an axis enriched for renal solute transporter programs (Reactome: Multifunctional Anion Exchangers). PC1 and PC3 carried no independent survival information. In the joint model adjusted for stage and purity, PC2 retained significance (HR = 0.73; $p = 0.012$). In LIHC, all three axes achieved nominal significance in signed univariable models, with PC1 showing the strongest association (HR = 0.82; $p = 0.008$), enriched for hepatocyte-specific metabolic programs.

#### S8. Stemness comparison

To evaluate the relationship between tissue memory and transcriptional stemness, TMS was compared with two stemness metrics: a gene-signature proxy and the published OCLR-derived mRNAsi scores from Malta et al. [13].

The gene-signature proxy was computed from 15 pluripotency-associated genes (POU5F1, SOX2, NANOG, KLF4, MYC, LIN28A, DNMT3B, TDGF1, SALL4, ZFP42, DPPA4, GDF3, TERT, FOXD3, UTF1) identified in TCGA counts matrices by hardcoded Ensembl-to-symbol mapping. For each sample, gene-level read counts were normalized to $\log_2(\text{CPM} + 1)$ and z-scored per gene across the cohort; the mean z-score constituted the stemness proxy. This signature-based proxy is not equivalent to the OCLR-derived mRNAsi but captures a correlated axis of transcriptional stemness.

Per-sample OCLR-derived mRNAsi scores were obtained directly from the supplementary data of Malta et al. [13] for TCGA samples with available identifiers. These scores were matched to the study cohorts by TCGA patient barcode. The OCLR-derived mRNAsi was used as the primary stemness metric in the joint Cox models reported in the Results; the gene-signature proxy served as an internal consistency check. Spearman correlation between the OCLR-derived mRNAsi and the gene-signature proxy was computed per cohort to assess concordance.

Three nested Cox models were fitted in KIRC and LIHC: Model A (TMS_z + stage + purity), Model B (TMS_z + mRNAsi + stage + purity), and Model C (mRNAsi + stage + purity). Comparison of TMS hazard ratios between Models A and B assessed whether TMS retained independent prognostic value after stemness adjustment. AIC differences and concordance index changes between models quantified the incremental information contributed by each metric.

#### S9. Clinical scenario: intermediate-stage ccRCC

To translate TMS into a clinically interpretable risk stratification, KIRC patients were restricted to AJCC pathologic stage II and III (the intermediate-risk subset where post-nephrectomy management decisions are most contested; n = 181, 62 OS events). TMS tertile boundaries were computed within this subset. Kaplan-Meier curves were fitted per tertile with log-rank testing. Absolute survival probabilities at three-year and five-year horizons were extracted from the step-function survival estimates with 95% confidence intervals from the Greenwood variance formula. Absolute risk differences between the low and high TMS tertiles were computed as the difference in cumulative mortality at each horizon. Univariable (OS ~ TMS_z) and adjusted (OS ~ TMS_z + stage + purity) Cox models were fitted within the intermediate-stage subset to verify that TMS provided prognostic discrimination beyond what residual stage heterogeneity (stage II vs. III) and purity contribute.

#### S10. Robustness and sensitivity analyses

**PCA dimensionality sensitivity.** TMS was recomputed for k = 2, 3, 4, 5, 6, 8, and 10 retained PCs. C1 accuracy and univariable Cox HR were evaluated at each k to assess stability of findings with respect to the dimensionality parameter. Non-monotonic C1 accuracy was observed for COAD, which dropped to 1.3% at k = 4 before recovering to 94.8% at k = 5 and 99.4% at k = 8. This behavior arises because the fourth principal component introduces an axis that differentiates colonic from gastric mucosal transcription with sufficient magnitude to shift nearest-centroid boundaries: at k = 4, the majority of COAD tumors are reassigned to the gastric centroid. The effect is transient because additional components (k ≥ 5) restore colon-specific axes that re-establish correct centroid proximity. This sensitivity to centroid boundary topology at specific dimensionalities reinforces the use of the primary k = 8 specification, at which C1 accuracy is stable for all cohorts retaining tissue identity.

**Subsampling stability.** GTEx reference samples were subsampled at 50%, 70%, and 90% fractions (200 iterations per fraction). Pairwise centroid distances between all tissue pairs were recomputed at each subsample to assess stability of the tissue separation structure.

**Leave-one-organ-out analysis.** Each of the eight organs was sequentially removed from the GTEx reference, PCA was refitted on the remaining seven organs, and the subspace angle between the reduced and full reference was computed. This analysis evaluates the sensitivity of the PCA coordinate system to the specific composition of the tissue panel. The maximum subspace angles observed upon removal of Liver (20.4°) and Thyroid (20.6°) exceed the 15° bootstrap stability threshold applied to within-reference resampling (subsampling stability, above). These two stability analyses address distinct questions: bootstrap resampling evaluates the robustness of the PCA subspace to donor-level sampling variation within a fixed tissue composition, while leave-one-organ-out assesses the structural sensitivity of the subspace to removal of an entire tissue class. The 15° threshold is specific to the bootstrap context; the leave-one-organ-out analysis does not impose the same threshold because removal of a complete organ is expected to produce larger angular shifts than resampling within a fixed panel. The moderate angles for Liver and Thyroid reflect the transcriptional distinctiveness of hepatocyte and thyrocyte programs, which contribute substantially to the variance captured by the first four PCs.

**Gene set enrichment of PCA loadings.** The top 50 genes by absolute loading magnitude in each direction (positive, negative) for each of the retained principal components were submitted to Enrichr for functional enrichment against Gene Ontology Biological Process, KEGG, and Reactome pathway databases.

**Random-gene negative control.** One thousand random gene sets, each of size 3,000, were drawn from the expressed genome. For each random set, PCA was refitted on GTEx, TMS was recomputed for all TCGA tumors, and a univariable Cox model was fitted. The false positive rate was defined as the fraction of random iterations yielding a significant ($p < 0.05$) TMS HR. The expected false positive rate under the null is 5%.

**Additional sensitivity models.** The following pre-specified sensitivity models were fitted for eligible cohorts: (i) rank-normalized TMS (within-cohort percentile transformation) in both univariable and multivariable Cox models; (ii) median-dichotomized TMS (above/below median) in univariable and multivariable models; (iii) stage-stratified Cox models (TMS_z + purity + TMB, stratified by early/late stage); (iv) purity-residualized TMS in univariable and multivariable models (KIRC); (v) RMST analysis as a non-parametric complement to Cox regression.

#### S11. External validation details

The locked PCA reference (HVG list, PCA loading matrix, reference mean vector, and organ centroids) was applied to each validation cohort without re-estimation of any parameter. Validation expression matrices were normalized to match the reference: $\log_2$(CPM + 1) for CPTAC-ccRCC RNA-seq; $\log_2$-transformed pre-normalized intensities for GSE76427 microarray. Gene identifiers were mapped to HGNC symbols and intersected with the locked HVG list; overlap was 97.0% for CPTAC-ccRCC and 88.2% for GSE76427 (below the 90% target specified in the design document). TMS was computed by projection and distance to the matched centroid, then standardized within the validation cohort.

The CPTAC-ccRCC survival-annotated subset comprised 35 patients with zero censoring, reflecting the availability of survival timing data exclusively for deceased patients at the CPTAC data freeze. The implications of this event structure for Cox model estimation are noted in the main text.

Distributional concordance between validation and discovery TMS was assessed by Kolmogorov-Smirnov test and by comparing distribution moments (mean, SD). C1 accuracy was computed in the validation cohort. Univariable and multivariable Cox models were fitted using the same covariate specification as the discovery analysis, subject to covariate availability. Kaplan-Meier analysis by TMS tertiles was performed with log-rank testing.

Replication was assessed by three pre-specified criteria: (R1) concordant HR direction, (R2) univariable $p < 0.05$, and (R3) overlapping 95% CIs between discovery and validation HRs. Full replication required all three; partial replication required R1 with failure of R2 or R3. A fixed-effect meta-analysis pooling the discovery and validation cohorts for KIRC was conducted to obtain a combined HR estimate and to quantify between-cohort heterogeneity by $I^2$.

#### Supplementary Figures

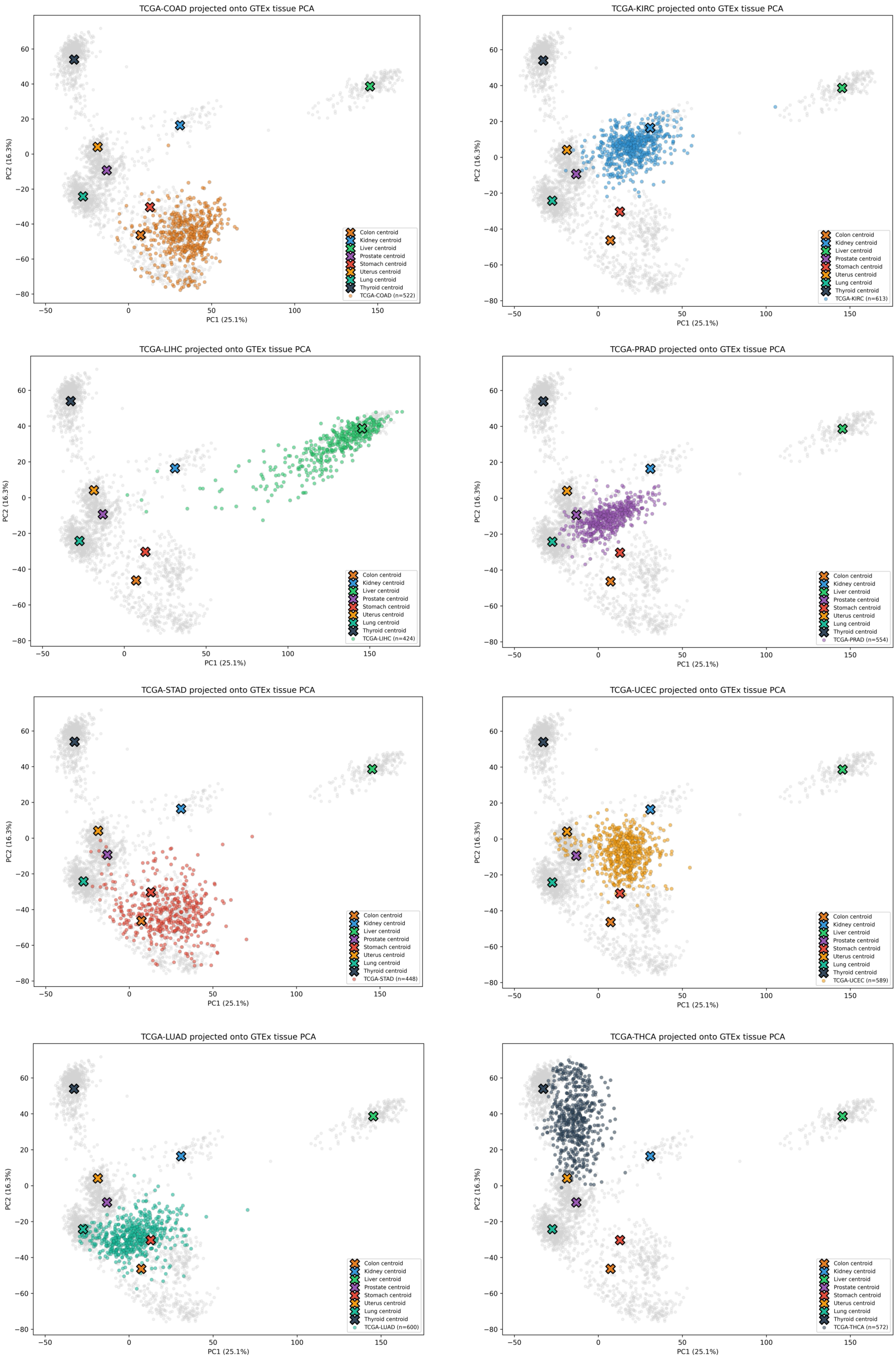


**Supplementary Figure S1.** All eight TCGA tumor cohort projections onto the locked GTEx V8 PCA reference. Each panel shows one TCGA cohort (colored points) overlaid on the GTEx reference (grey) in PC1 vs PC2 space. Crosses denote the eight tissue centroids. Cohorts shown: COAD, KIRC, LIHC, PRAD, STAD, UCEC, LUAD, THCA.

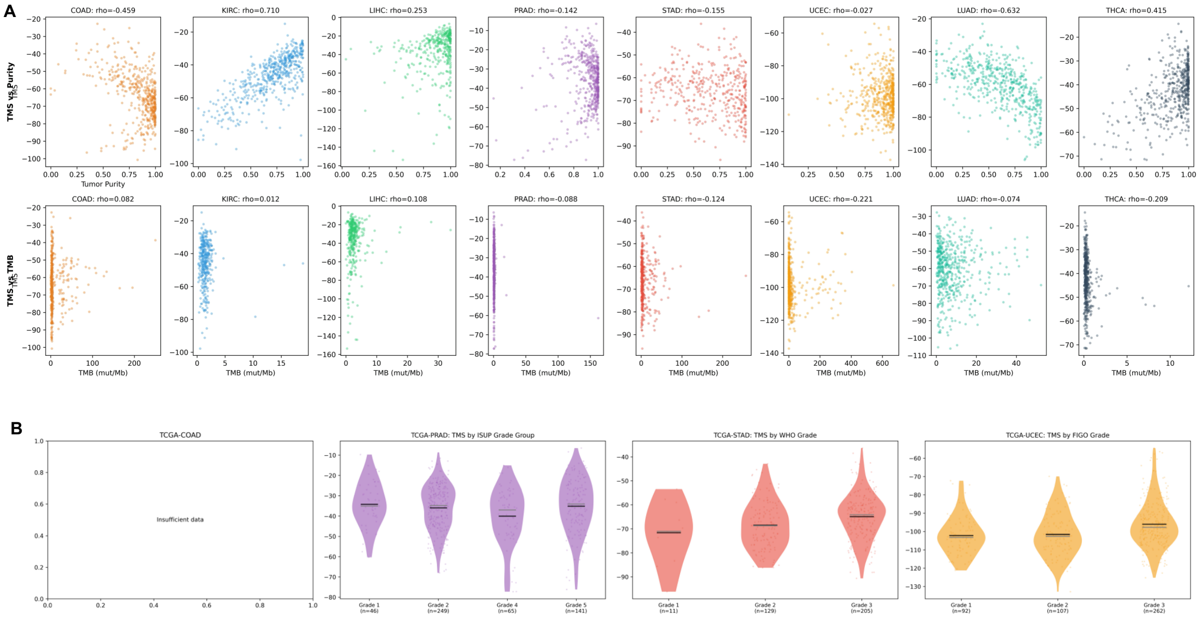

**Supplementary Figure S2.** Covariate correlations and grade-based C2 testing. (A) Scatter plots of TMS versus ESTIMATE tumor purity (top row) and TMS versus tumor mutational burden (bottom row) for all eight cohorts; Spearman ρ values annotated per panel. (B) TMS distributions stratified by histological grade for cohorts with established grading systems: PRAD (ISUP Grade Group), STAD (WHO grade), and UCEC (FIGO grade).

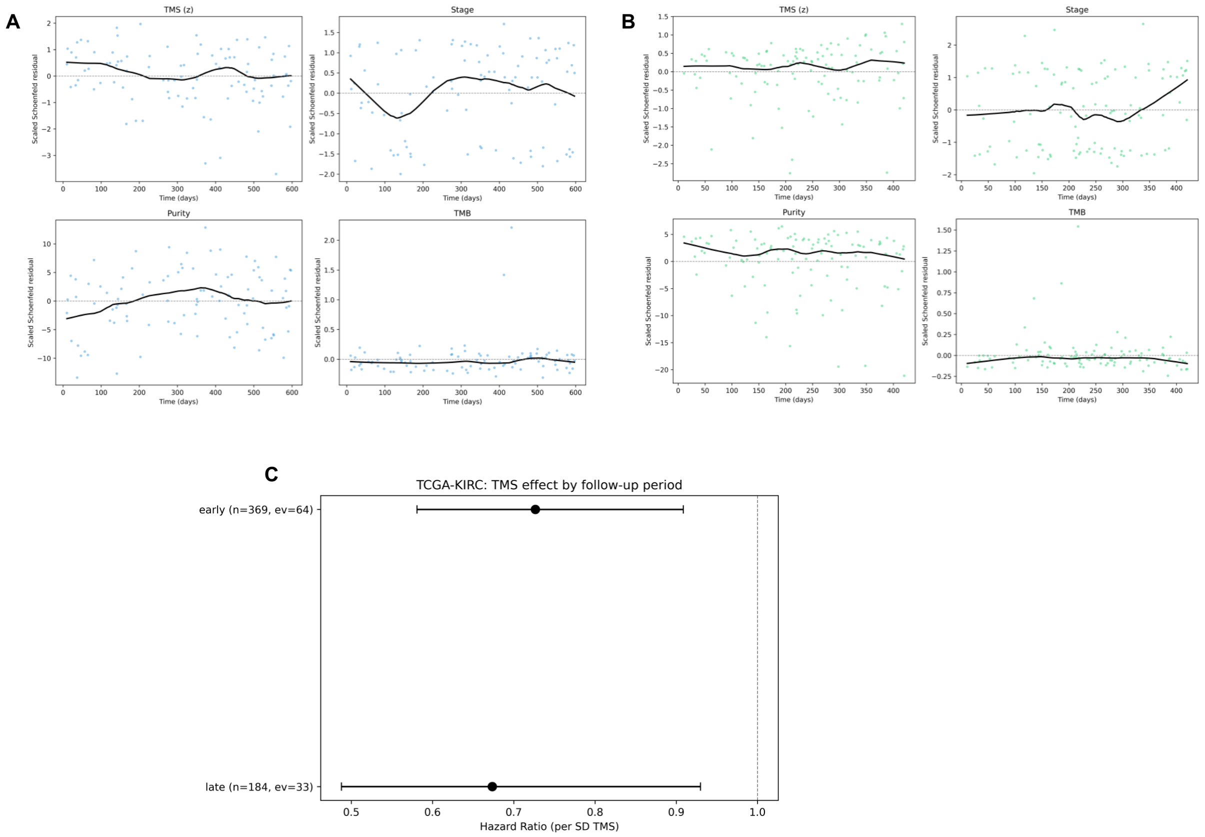


**Supplementary Figure S3.** Proportional hazards diagnostics. (A) Scaled Schoenfeld residuals plotted against ranked event times for each covariate in the KIRC multivariable Cox model, with LOWESS trend lines. (B) Corresponding Schoenfeld residual plots for the LIHC multivariable model. (C) Time-varying coefficient analysis for TMS in KIRC: piecewise Cox hazard ratios for early and late follow-up periods.

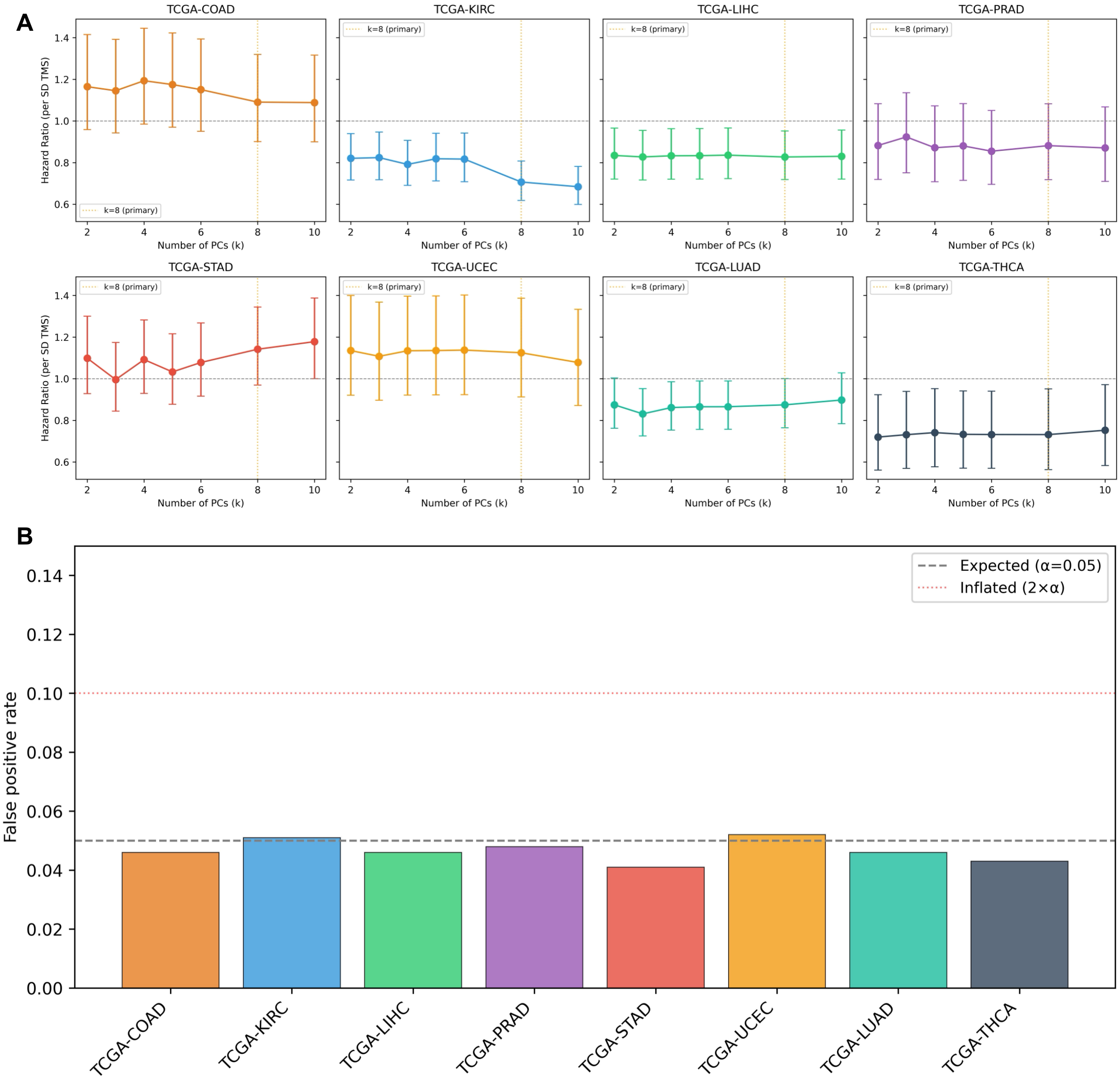


**Supplementary Figure S4.** Robustness analyses. (A) PCA dimensionality sensitivity: univariable TMS hazard ratios as a function of retained principal components (k = 2 to 10) for all eight cohorts; dashed vertical line indicates primary analysis dimensionality (k = 8). (B) Random-gene negative control: false positive rates from 1,000 iterations of random gene-set PCA and univariable Cox modeling; dashed line, expected 5% null rate.

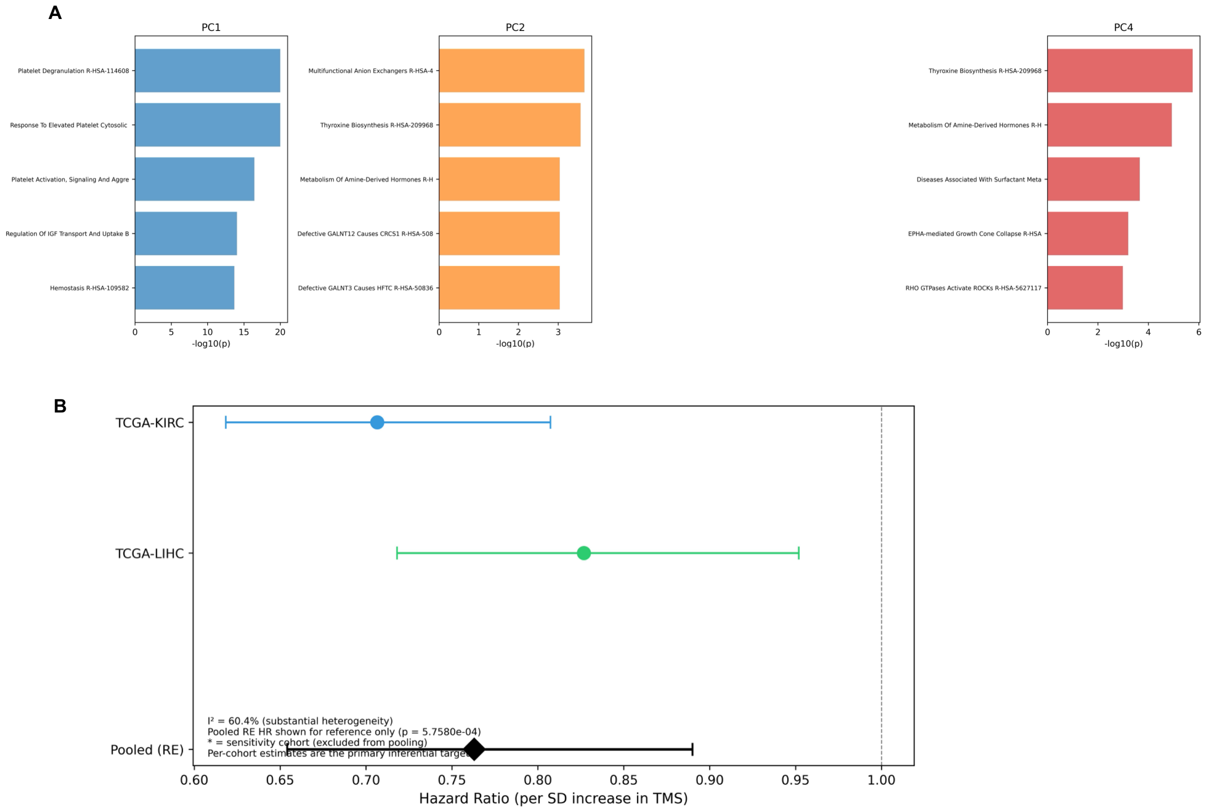


**Supplementary Figure S5.** Extended robustness and pan-cancer forest. (A) Top Reactome enrichment terms for PCA loading genes of PC1, PC2, and PC4, ranked by adjusted p-value. (B) Forest plot of univariable TMS hazard ratios for all eight TCGA cohorts with pooled random-effects estimate.

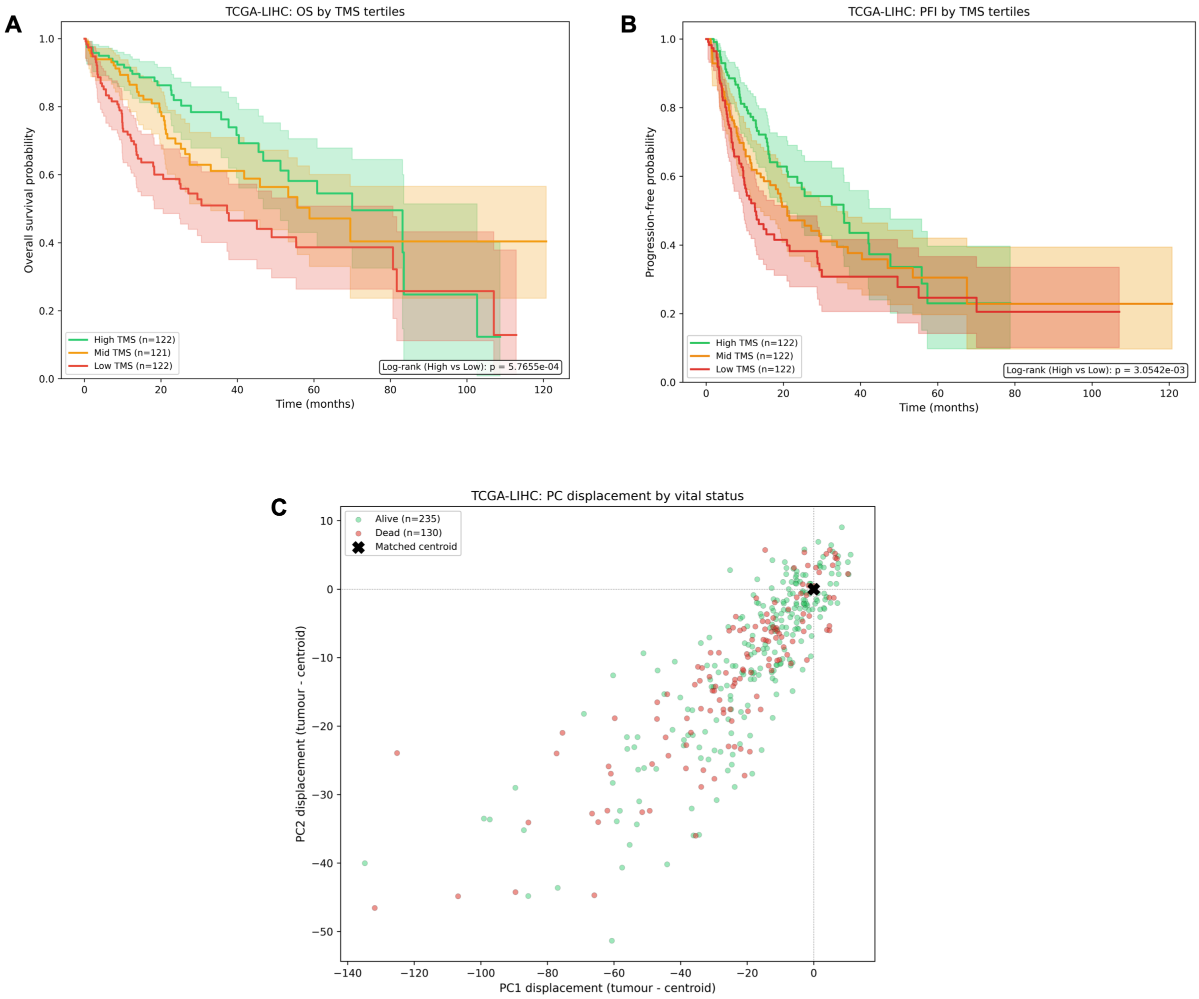


**Supplementary Figure S6.** LIHC detailed results (exploratory). (A) Kaplan–Meier overall survival curves by TMS tertiles in TCGA-LIHC (n = 365; 130 events). (B) Kaplan–Meier progression-free interval curves by TMS tertiles in TCGA-LIHC (n = 366; 180 PFI events). (C) Per-sample displacement from the matched liver centroid in PC1 vs PC2 space, colored by vital status.

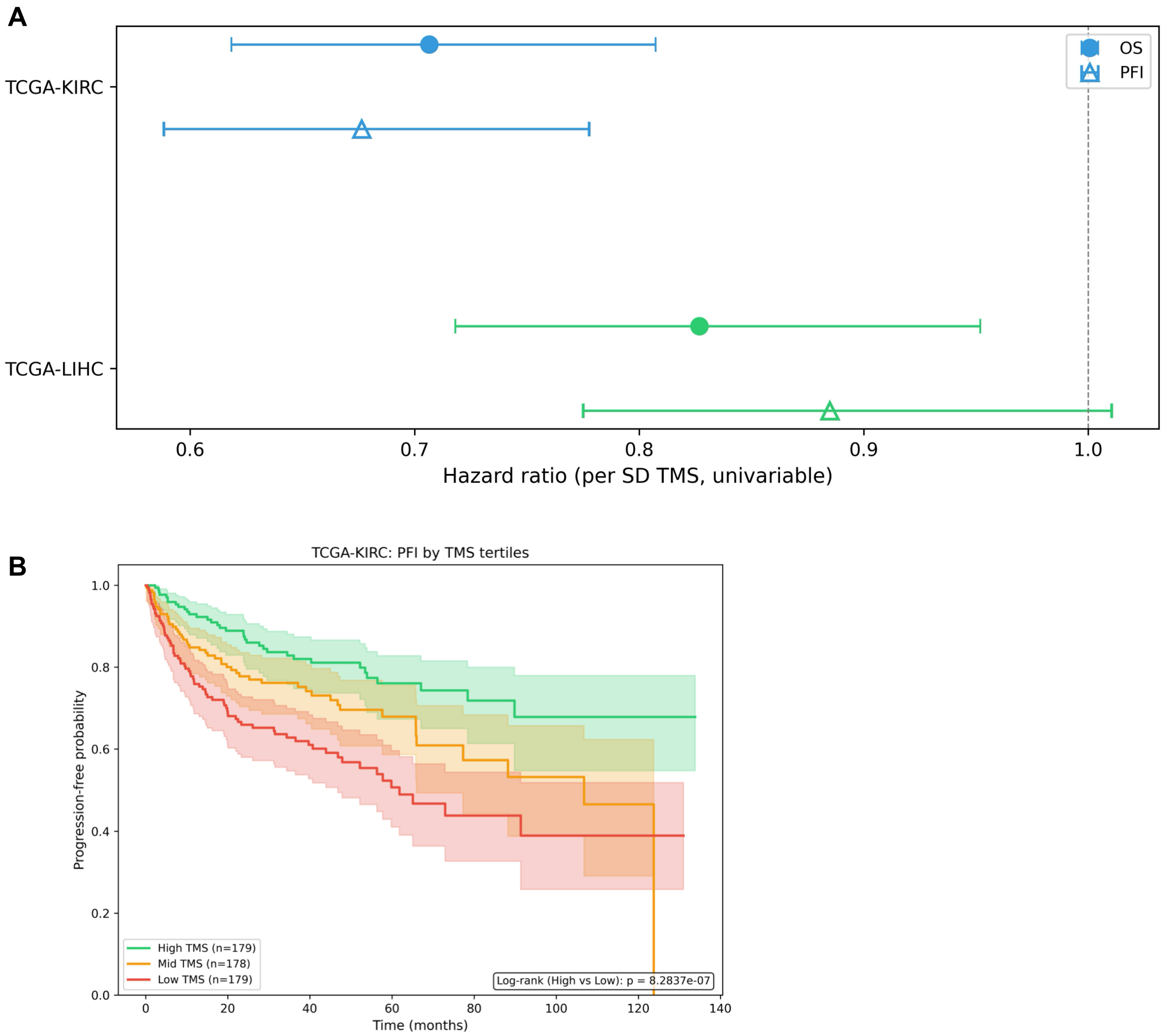


**Supplementary Figure S7.** Progression-free interval endpoint sensitivity. (A) Forest plot comparing univariable TMS hazard ratios for overall survival (filled circles) and progression-free interval (open triangles) in KIRC and LIHC. (B) Kaplan–Meier progression-free interval curves by TMS tertiles in TCGA-KIRC (n = 536; 159 PFI events).

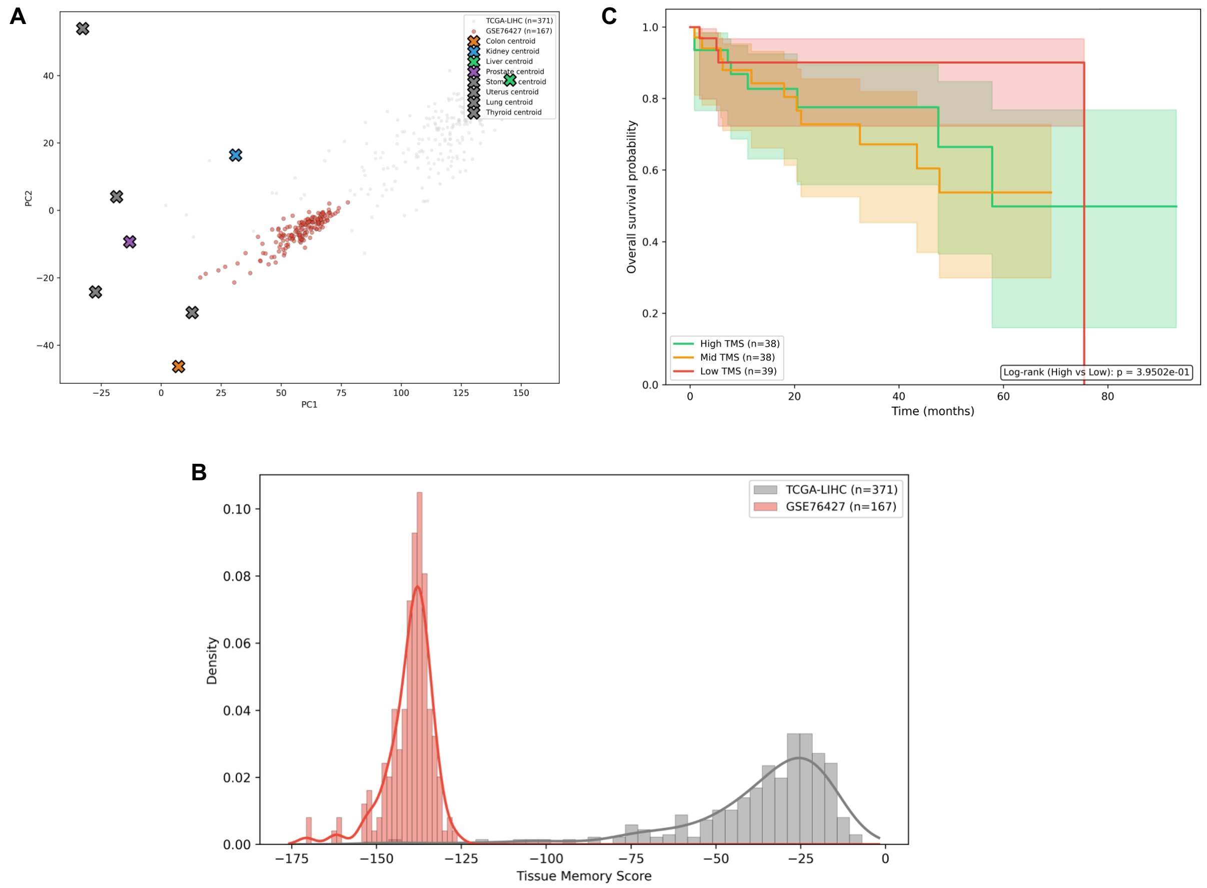


**Supplementary Figure S8.** GSE76427 validation failure (LIHC). (A) GSE76427 HCC tumors (n = 167, Illumina HumanHT-12 v4 microarray) projected onto the locked GTEx PCA axes alongside TCGA-LIHC tumors (grey), showing platform-driven displacement from the liver centroid. (B) TMS density distributions for TCGA-LIHC (discovery) and GSE76427 (validation), demonstrating a distributional shift exceeding 4.4 SD units. (C) Kaplan–Meier overall survival curves by TMS tertiles in GSE76427 (n = 115; 23 events), showing no separation.

### Supplementary Tables

**Supplementary Table S1.** Univariable Cox proportional hazards models for TMS and overall survival across all eight TCGA cohorts. Hazard ratios are per standard deviation increase.

| Cohort | Covariate | n | Events | HR (95% CI) | log(HR) | SE(log HR) | p-value | C-index | Primary |
|---|---|---|---|---|---|---|---|---|---|
| KIRC | TMS | 538 | 175 | 0.71 (0.62–0.81) | -0.3474 | 0.0680 | $3.2 \times 10^{-7}$ | 0.616 | Yes |
| KIRC | Stage | 535 | 174 | 1.89 (1.66–2.16) | 0.6374 | 0.0666 | $1.1 \times 10^{-21}$ | 0.732 | Yes |
| KIRC | Purity | 538 | 175 | 0.45 (0.25–0.83) | -0.7922 | 0.3075 | 0.0100 | 0.548 | Yes |
| KIRC | TMB | 371 | 97 | 1.12 (1.04–1.22) | 0.1176 | 0.0402 | 0.0035 | 0.617 | Yes |
| LIHC | TMS | 365 | 130 | 0.83 (0.72–0.95) | -0.1902 | 0.0719 | 0.0081 | 0.620 | Yes |
| LIHC | Stage | 341 | 116 | 1.66 (1.35–2.04) | 0.5075 | 0.1039 | $1.1 \times 10^{-6}$ | 0.609 | Yes |
| LIHC | Purity | 365 | 130 | 1.01 (0.34–3.00) | 0.0096 | 0.5559 | 0.986 | 0.492 | Yes |
| LIHC | TMB | 352 | 122 | 1.05 (1.01–1.10) | 0.0497 | 0.0226 | 0.028 | 0.589 | Yes |

**Supplementary Table S2.** Multivariable Cox proportional hazards models (TMS, stage, purity, TMB) for gate-passing cohorts (KIRC, LIHC).

| Cohort | Covariate | Coefficient | SE | HR (95% CI) | z | p-value | C-index | AIC | n | Events |
|---|---|---|---|---|---|---|---|---|---|---|
| KIRC | TMS | -0.5195 | 0.1302 | 0.59 (0.46–0.77) | -3.99 | $6.6 \times 10^{-5}$ | 0.755 | 949.36 | 369 | 97 |
| KIRC | Stage | 0.6994 | 0.0946 | 2.01 (1.67–2.42) | 7.39 | $1.4 \times 10^{-13}$ | 0.755 | 949.36 | 369 | 97 |
| KIRC | Purity | 1.5740 | 0.5933 | 4.83 (1.51–15.44) | 2.65 | 0.0080 | 0.755 | 949.36 | 369 | 97 |
| KIRC | TMB | 0.1581 | 0.0429 | 1.17 (1.08–1.27) | 3.69 | $2.3 \times 10^{-4}$ | 0.755 | 949.36 | 369 | 97 |
| LIHC | TMS | -0.1340 | 0.0838 | 0.87 (0.74–1.03) | -1.60 | 0.110 | 0.649 | 1102.27 | 331 | 111 |
| LIHC | Stage | 0.4780 | 0.1082 | 1.61 (1.30–1.99) | 4.42 | $1.0 \times 10^{-5}$ | 0.649 | 1102.27 | 331 | 111 |
| LIHC | Purity | 0.1352 | 0.5729 | 1.14 (0.37–3.52) | 0.24 | 0.813 | 0.649 | 1102.27 | 331 | 111 |
| LIHC | TMB | 0.0432 | 0.0241 | 1.04 (1.00–1.09) | 1.79 | 0.073 | 0.649 | 1102.27 | 331 | 111 |

**Supplementary Table S3.** Proportional hazards assumption testing by Schoenfeld residual correlation for all covariates in the KIRC and LIHC multivariable models.

| Cohort | Covariate | Spearman ρ | p-value | PH violated |
|---|---|---|---|---|
| KIRC | TMS | -0.189 | 0.064 | No |
| KIRC | Stage | 0.111 | 0.279 | No |
| KIRC | Purity | 0.105 | 0.307 | No |
| KIRC | TMB | 0.123 | 0.230 | No |
| LIHC | TMS | 0.113 | 0.238 | No |
| LIHC | Stage | 0.118 | 0.218 | No |
| LIHC | Purity | -0.108 | 0.261 | No |
| LIHC | TMB | -0.070 | 0.467 | No |

**Supplementary Table S4.** Variance inflation factor diagnostics for the KIRC and LIHC multivariable Cox models.

| Cohort | VIF (TMS) | VIF (Stage) | VIF (Purity) | VIF (TMB) |
|---|---|---|---|---|
| KIRC | 2.30 | 1.05 | 2.33 | 1.01 |
| LIHC | 1.15 | 1.09 | 1.06 | 1.02 |

**Supplementary Table S5.** Bootstrap confidence intervals for Shapley decomposition of Cox model pseudo-$R^2$ (500 case-level resamples).

| Cohort | Covariate | Mean | Median | 95% CI Lower | 95% CI Upper | SD | Converged (n) | Convergence rate |
|---|---|---|---|---|---|---|---|---|
| KIRC | TMS | 0.0299 | 0.0278 | 0.0072 | 0.0658 | 0.0151 | 500 | 100.0% |
| KIRC | Stage | 0.1353 | 0.1343 | 0.0706 | 0.2082 | 0.0356 | 500 | 100.0% |
| KIRC | Purity | 0.0105 | 0.0096 | 0.0034 | 0.0221 | 0.0046 | 500 | 100.0% |
| KIRC | TMB | 0.0222 | 0.0190 | 0.0038 | 0.0709 | 0.0168 | 500 | 100.0% |
| LIHC | TMS | 0.0128 | 0.0109 | 5.76e-04 | 0.0433 | 0.0106 | 500 | 100.0% |
| LIHC | Stage | 0.0624 | 0.0603 | 0.0206 | 0.1140 | 0.0250 | 500 | 100.0% |
| LIHC | Purity | 0.0030 | 0.0017 | 8.20e-05 | 0.0156 | 0.0040 | 500 | 100.0% |
| LIHC | TMB | 0.0128 | 0.0092 | 3.13e-04 | 0.0419 | 0.0119 | 500 | 100.0% |

**Supplementary Table S6.** DerSimonian–Laird random-effects and fixed-effect meta-analysis of univariable TMS hazard ratios from eligible cohorts.

| k | Cohorts | RE HR (95% CI) | RE z | RE p-value | Q | Q df | Q p-value | $I^2$ (%) | $\tau^2$ | Direction consistent | C4 pass |
|---|---|---|---|---|---|---|---|---|---|---|---|
| 2 | KIRC,LIHC | 0.76 (0.65–0.89) | -3.44 | $5.8 \times 10^{-4}$ | 2.52 | 1 | 0.112 | 60.37 | 0.0075 | Yes | Yes |

**Supplementary Table S7.** Per-cohort inputs to the meta-analysis: log hazard ratios, standard errors, and random-effects weights.

| Cohort | log(HR) | SE(log HR) | HR (95% CI) | RE weight |
|---|---|---|---|---|
| KIRC | -0.3474 | 0.0680 | 0.71 (0.62–0.81) | 51.100 |
| LIHC | -0.1902 | 0.0719 | 0.83 (0.72–0.95) | 48.900 |

**Supplementary Table S8.** Restricted mean survival time by TMS tertiles for KIRC and LIHC.

| Cohort | TMS group | n | Events | τ (days) | RMST (days) | RMST (months) | RMST (years) |
|---|---|---|---|---|---|---|---|
| COAD | Low | 154 | 38 | 1826.20 | 1449 | 47.60 | 3.97 |
| COAD | Mid | 152 | 25 | 1826.20 | 1519.50 | 49.90 | 4.16 |
| COAD | High | 154 | 36 | 1826.20 | 1450.30 | 47.60 | 3.97 |
| KIRC | Low | 180 | 83 | 1826.20 | 1307.10 | 42.90 | 3.58 |
| KIRC | Mid | 179 | 51 | 1826.20 | 1486.10 | 48.80 | 4.07 |
| KIRC | High | 179 | 41 | 1826.20 | 1605.70 | 52.70 | 4.40 |
| LIHC | Low | 122 | 57 | 1826.20 | 1045 | 34.30 | 2.86 |
| LIHC | Mid | 121 | 40 | 1826.20 | 1269.60 | 41.70 | 3.48 |
| LIHC | High | 122 | 33 | 1826.20 | 1427 | 46.90 | 3.91 |
| PRAD | Low | 167 | 34 | 1826.20 | 1492 | 49 | 4.08 |
| PRAD | Mid | 167 | 30 | 1826.20 | 1537.10 | 50.50 | 4.21 |
| PRAD | High | 167 | 29 | 1826.20 | 1553 | 51 | 4.25 |
| STAD | Low | 130 | 48 | 1826.20 | 1164 | 38.20 | 3.19 |
| STAD | Mid | 129 | 53 | 1826.20 | 1055.20 | 34.70 | 2.89 |
| STAD | High | 129 | 55 | 1826.20 | 969.90 | 31.90 | 2.66 |
| UCEC | Low | 184 | 27 | 1826.20 | 1615.70 | 53.10 | 4.42 |
| UCEC | Mid | 183 | 30 | 1826.20 | 1591.20 | 52.30 | 4.36 |
| UCEC | High | 184 | 36 | 1826.20 | 1540 | 50.60 | 4.22 |
| LUAD | Low | 176 | 78 | 1826.20 | 1188.50 | 39 | 3.25 |
| LUAD | Mid | 175 | 56 | 1826.20 | 1269.40 | 41.70 | 3.48 |
| LUAD | High | 175 | 54 | 1826.20 | 1357.80 | 44.60 | 3.72 |
| THCA | Low | 168 | 22 | 1826.20 | 1611.10 | 52.90 | 4.41 |
| THCA | Mid | 168 | 22 | 1826.20 | 1616.20 | 53.10 | 4.42 |
| THCA | High | 168 | 8 | 1826.20 | 1742.80 | 57.30 | 4.77 |

**Supplementary Table S9.** Stage-stratified Cox regression (TMS + purity + TMB, stratified by early/late stage) for eligible cohorts.

| Cohort | Covariate | HR (95% CI) | p-value | C-index | Model |
|---|---|---|---|---|---|
| COAD | TMS | 0.98 (0.79–1.22) | 0.874 | 0.560 | Stratified early late |
| COAD | Purity | 0.73 (0.23–2.37) | 0.606 | 0.560 | Stratified early late |
| COAD | TMB | 1.01 (1.00–1.02) | 0.061 | 0.560 | Stratified early late |
| KIRC | TMS | 0.60 (0.46–0.77) | $8.0 \times 10^{-5}$ | 0.637 | Stratified early late |
| KIRC | Purity | 5.17 (1.60–16.67) | 0.0059 | 0.637 | Stratified early late |
| KIRC | TMB | 1.17 (1.07–1.28) | $3.8 \times 10^{-4}$ | 0.637 | Stratified early late |
| LIHC | TMS | 0.83 (0.70–0.98) | 0.032 | 0.609 | Stratified early late |
| LIHC | Purity | 1.14 (0.37–3.49) | 0.824 | 0.609 | Stratified early late |
| LIHC | TMB | 1.06 (1.01–1.10) | 0.021 | 0.609 | Stratified early late |
| PRAD | TMS | 0.96 (0.81–1.15) | 0.695 | 0.645 | Penalized ridge |
| PRAD | Stage | 1.85 (1.31–2.61) | $4.3 \times 10^{-4}$ | 0.645 | Penalized ridge |
| PRAD | Purity | 0.82 (0.16–4.29) | 0.813 | 0.645 | Penalized ridge |
| PRAD | TMB | 1.01 (1.00–1.03) | 0.079 | 0.645 | Penalized ridge |
| STAD | TMS | 1.13 (0.95–1.35) | 0.167 | 0.561 | Stratified early late |
| STAD | Purity | 0.76 (0.42–1.36) | 0.347 | 0.561 | Stratified early late |
| STAD | TMB | 0.99 (0.97–1.00) | 0.046 | 0.561 | Stratified early late |
| LUAD | TMS | 0.88 (0.73–1.06) | 0.172 | 0.538 | Stratified early late |
| LUAD | Purity | 0.92 (0.45–1.88) | 0.824 | 0.538 | Stratified early late |
| LUAD | TMB | 1.00 (0.98–1.01) | 0.655 | 0.538 | Stratified early late |
| THCA | TMS | 0.67 (0.49–0.92) | 0.014 | 0.597 | Stratified early late |
| THCA | Purity | 2.65 (0.48–14.83) | 0.266 | 0.597 | Stratified early late |
| THCA | TMB | 0.92 (0.63–1.35) | 0.673 | 0.597 | Stratified early late |

**Supplementary Table S10.** Median-dichotomized TMS Cox models (univariable and multivariable) for all cohorts.

| Cohort | Covariate | HR (95% CI) | p-value | Model |
|---|---|---|---|---|
| COAD | TMS (high vs low) | 0.86 (0.53–1.40) | 0.536 | Binary median |
| COAD | Stage | 2.31 (1.77–3.02) | $6.3 \times 10^{-10}$ | Binary median |
| COAD | Purity | 0.52 (0.16–1.72) | 0.282 | Binary median |
| COAD | TMB | 1.01 (1.00–1.01) | 0.081 | Binary median |
| KIRC | TMS (high vs low) | 0.49 (0.29–0.81) | 0.0053 | Binary median |
| KIRC | Stage | 1.99 (1.66–2.39) | $1.7 \times 10^{-13}$ | Binary median |
| KIRC | Purity | 2.12 (0.75–6.03) | 0.157 | Binary median |
| KIRC | TMB | 1.15 (1.06–1.26) | $9.7 \times 10^{-4}$ | Binary median |
| LIHC | TMS (high vs low) | 0.68 (0.46–1.00) | 0.050 | Binary median |
| LIHC | Stage | 1.62 (1.31–2.00) | $7.7 \times 10^{-6}$ | Binary median |
| LIHC | Purity | 1.22 (0.40–3.76) | 0.729 | Binary median |
| LIHC | TMB | 1.05 (1.00–1.10) | 0.063 | Binary median |
| PRAD | TMS (high vs low) | 0.26 (0.04–1.95) | 0.191 | Binary median |
| PRAD | Stage | 1.04 (0.46–2.38) | 0.919 | Binary median |
| PRAD | Purity | 0.00 (0.00–0.00) † | $2.4 \times 10^{-23}$ | Binary median |
| PRAD | TMB | 1.12 (1.07–1.19) | $1.6 \times 10^{-5}$ | Binary median |
| STAD | TMS (high vs low) | 1.40 (1.00–1.96) | 0.049 | Binary median |
| STAD | Stage | 1.59 (1.29–1.96) | $1.3 \times 10^{-5}$ | Binary median |
| STAD | Purity | 0.80 (0.45–1.44) | 0.457 | Binary median |
| STAD | TMB | 0.99 (0.97–1.00) | 0.031 | Binary median |
| LUAD | TMS (high vs low) | 0.76 (0.53–1.09) | 0.131 | Binary median |
| LUAD | Stage | 1.66 (1.44–1.91) | $1.1 \times 10^{-12}$ | Binary median |
| LUAD | Purity | 0.90 (0.46–1.75) | 0.752 | Binary median |
| LUAD | TMB | 1.00 (0.98–1.02) | 0.809 | Binary median |
| THCA | TMS (high vs low) | 0.57 (0.30–1.05) | 0.073 | Binary median |
| THCA | Stage | 1.58 (1.24–2.02) | $2.5 \times 10^{-4}$ | Binary median |
| THCA | Purity | 1.68 (0.33–8.61) | 0.533 | Binary median |
| THCA | TMB | 0.90 (0.62–1.31) | 0.582 | Binary median |

†PRAD Purity estimates (HR = 0.00) are numerically degenerate, arising from quasi-complete separation of the Purity covariate with respect to the event indicator in a cohort with fewer than 10 multivariable OS events. The maximum partial likelihood estimator for the Purity coefficient diverges under these conditions, producing $\exp(-\infty) \approx 0.00$ and an artificially extreme Wald p-value. PRAD is excluded from primary inference by the pre-specified decision gate (C2 failure); these sensitivity models are provided for completeness.

**Supplementary Table S11.** Rank-normalized TMS Cox models (univariable and multivariable) for all cohorts.

| Cohort | Model | Covariate | HR (95% CI) | p-value | n | Events |
|---|---|---|---|---|---|---|
| COAD | Univariable rank | TMS rank | 1.31 (0.67–2.57) | 0.427 | 460 | 99 |
| COAD | Multivariable rank | TMS rank | 0.72 (0.33–1.57) | 0.404 | 393 | 85 |
| COAD | Multivariable rank | Stage | 2.33 (1.78–3.05) | $6.0 \times 10^{-10}$ | 393 | 85 |
| COAD | Multivariable rank | Purity | 0.49 (0.15–1.61) | 0.240 | 393 | 85 |
| COAD | Multivariable rank | TMB | 1.01 (1.00–1.01) | 0.070 | 393 | 85 |
| KIRC | Univariable rank | TMS rank | 0.26 (0.15–0.44) | $6.5 \times 10^{-7}$ | 538 | 175 |
| KIRC | Multivariable rank | TMS rank | 0.15 (0.05–0.43) | $4.5 \times 10^{-4}$ | 369 | 97 |
| KIRC | Multivariable rank | Stage | 2.00 (1.67–2.41) | $1.3 \times 10^{-13}$ | 369 | 97 |
| KIRC | Multivariable rank | Purity | 5.01 (1.41–17.79) | 0.013 | 369 | 97 |
| KIRC | Multivariable rank | TMB | 1.16 (1.07–1.26) | $4.8 \times 10^{-4}$ | 369 | 97 |
| LIHC | Univariable rank | TMS rank | 0.35 (0.19–0.64) | $7.1 \times 10^{-4}$ | 365 | 130 |
| LIHC | Multivariable rank | TMS rank | 0.42 (0.21–0.84) | 0.014 | 331 | 111 |
| LIHC | Multivariable rank | Stage | 1.58 (1.28–1.96) | $2.5 \times 10^{-5}$ | 331 | 111 |
| LIHC | Multivariable rank | Purity | 1.36 (0.43–4.26) | 0.599 | 331 | 111 |
| LIHC | Multivariable rank | TMB | 1.05 (1.00–1.10) | 0.056 | 331 | 111 |
| PRAD | Univariable rank | TMS rank | 0.75 (0.37–1.52) | 0.422 | 501 | 93 |
| PRAD | Multivariable rank | TMS rank | 0.04 (0.00–0.39) | 0.0056 | 465 | 75 |
| PRAD | Multivariable rank | Stage | 0.47 (0.20–1.12) | 0.088 | 465 | 75 |
| PRAD | Multivariable rank | Purity | 0.00 (0.00–0.00) † | $7.3 \times 10^{-32}$ | 465 | 75 |
| PRAD | Multivariable rank | TMB | 1.02 (0.95–1.10) | 0.578 | 465 | 75 |
| STAD | Univariable rank | TMS rank | 1.60 (0.92–2.78) | 0.093 | 388 | 156 |
| STAD | Multivariable rank | TMS rank | 1.45 (0.82–2.60) | 0.205 | 370 | 144 |
| STAD | Multivariable rank | Stage | 1.60 (1.30–1.97) | $1.1 \times 10^{-5}$ | 370 | 144 |
| STAD | Multivariable rank | Purity | 0.78 (0.43–1.39) | 0.397 | 370 | 144 |
| STAD | Multivariable rank | TMB | 0.98 (0.97–1.00) | 0.032 | 370 | 144 |
| UCEC | Univariable rank | TMS rank | 1.81 (0.88–3.75) | 0.108 | 551 | 93 |
| LUAD | Univariable rank | TMS rank | 0.58 (0.36–0.94) | 0.028 | 526 | 188 |
| LUAD | Multivariable rank | TMS rank | 0.59 (0.31–1.15) | 0.121 | 513 | 184 |
| LUAD | Multivariable rank | Stage | 1.65 (1.44–1.90) | $1.4 \times 10^{-12}$ | 513 | 184 |
| LUAD | Multivariable rank | Purity | 0.84 (0.41–1.72) | 0.638 | 513 | 184 |
| LUAD | Multivariable rank | TMB | 1.00 (0.98–1.02) | 0.771 | 513 | 184 |
| THCA | Univariable rank | TMS rank | 0.28 (0.11–0.74) | 0.011 | 504 | 52 |
| THCA | Multivariable rank | TMS rank | 0.23 (0.07–0.72) | 0.012 | 486 | 49 |
| THCA | Multivariable rank | Stage | 1.59 (1.24–2.02) | $2.1 \times 10^{-4}$ | 486 | 49 |
| THCA | Multivariable rank | Purity | 2.44 (0.46–13.06) | 0.297 | 486 | 49 |
| THCA | Multivariable rank | TMB | 0.89 (0.60–1.32) | 0.562 | 486 | 49 |

†PRAD Purity estimates (HR = 0.00) in the rank-normalized multivariable model are numerically degenerate for the same reason described in Table S10: quasi-complete separation with sparse events. PRAD is excluded from primary inference.

**Supplementary Table S12.** PCA dimensionality sensitivity: C1 accuracy and univariable Cox HR across k = 2 to 10 retained principal components.

| Cohort | k (PCs) | C1 accuracy | HR (95% CI) | p-value |
|---|---|---|---|---|
| COAD | 2 | 46.5% | 1.16 (0.96–1.42) | 0.125 |
| COAD | 3 | 42.0% | 1.15 (0.94–1.39) | 0.173 |
| COAD | 4 | 1.3% | 1.19 (0.99–1.45) | 0.071 |
| COAD | 5 | 94.8% | 1.17 (0.97–1.42) | 0.099 |
| COAD | 6 | 91.7% | 1.15 (0.95–1.39) | 0.151 |
| COAD | 8 | 99.4% | 1.09 (0.90–1.32) | 0.373 |
| COAD | 10 | 99.6% | 1.09 (0.90–1.32) | 0.384 |
| KIRC | 2 | 79.5% | 0.82 (0.72–0.94) | 0.0042 |
| KIRC | 3 | 87.4% | 0.82 (0.72–0.95) | 0.0063 |
| KIRC | 4 | 93.1% | 0.79 (0.69–0.91) | $7.5 \times 10^{-4}$ |
| KIRC | 5 | 96.7% | 0.82 (0.71–0.94) | 0.0049 |
| KIRC | 6 | 98.3% | 0.82 (0.71–0.94) | 0.0054 |
| KIRC | 8 | 95.9% | 0.71 (0.62–0.81) | $3.2 \times 10^{-7}$ |
| KIRC | 10 | 95.2% | 0.68 (0.60–0.78) | $2.4 \times 10^{-8}$ |
| LIHC | 2 | 91.2% | 0.83 (0.72–0.97) | 0.015 |
| LIHC | 3 | 92.0% | 0.83 (0.72–0.95) | 0.0095 |
| LIHC | 4 | 91.5% | 0.83 (0.72–0.96) | 0.014 |
| LIHC | 5 | 91.2% | 0.83 (0.72–0.96) | 0.014 |
| LIHC | 6 | 91.8% | 0.84 (0.72–0.97) | 0.015 |
| LIHC | 8 | 96.2% | 0.83 (0.72–0.95) | 0.0081 |
| LIHC | 10 | 96.4% | 0.83 (0.72–0.96) | 0.0098 |
| PRAD | 2 | 50.1% | 0.88 (0.72–1.08) | 0.231 |
| PRAD | 3 | 47.7% | 0.92 (0.75–1.14) | 0.451 |
| PRAD | 4 | 35.5% | 0.87 (0.71–1.07) | 0.195 |
| PRAD | 5 | 54.9% | 0.88 (0.72–1.08) | 0.230 |
| PRAD | 6 | 97.4% | 0.85 (0.70–1.05) | 0.136 |
| PRAD | 8 | 100.0% | 0.88 (0.72–1.08) | 0.229 |
| PRAD | 10 | 100.0% | 0.87 (0.71–1.07) | 0.184 |
| STAD | 2 | 51.0% | 1.10 (0.93–1.30) | 0.274 |
| STAD | 3 | 71.1% | 1.00 (0.84–1.17) | 0.960 |
| STAD | 4 | 89.7% | 1.09 (0.93–1.28) | 0.284 |
| STAD | 5 | 42.8% | 1.03 (0.88–1.22) | 0.698 |
| STAD | 6 | 33.8% | 1.08 (0.92–1.27) | 0.365 |
| STAD | 8 | 44.3% | 1.14 (0.97–1.34) | 0.112 |
| STAD | 10 | 44.1% | 1.18 (1.00–1.39) | 0.050 |
| UCEC | 2 | 1.1% | 1.14 (0.92–1.40) | 0.236 |
| UCEC | 3 | 1.8% | 1.11 (0.90–1.37) | 0.343 |
| UCEC | 4 | 0.0% | 1.13 (0.92–1.40) | 0.235 |
| UCEC | 5 | 0.0% | 1.14 (0.92–1.40) | 0.232 |
| UCEC | 6 | 0.2% | 1.14 (0.92–1.40) | 0.226 |
| UCEC | 8 | 1.3% | 1.12 (0.91–1.39) | 0.271 |
| UCEC | 10 | 1.5% | 1.08 (0.87–1.33) | 0.489 |
| LUAD | 2 | 8.4% | 0.87 (0.76–1.00) | 0.057 |
| LUAD | 3 | 40.5% | 0.83 (0.73–0.95) | 0.0078 |
| LUAD | 4 | 53.4% | 0.86 (0.75–0.99) | 0.030 |
| LUAD | 5 | 66.7% | 0.87 (0.76–0.99) | 0.034 |
| LUAD | 6 | 68.1% | 0.87 (0.76–0.99) | 0.034 |
| LUAD | 8 | 80.0% | 0.87 (0.76–1.00) | 0.051 |
| LUAD | 10 | 80.6% | 0.90 (0.78–1.03) | 0.119 |
| THCA | 2 | 58.3% | 0.72 (0.56–0.92) | 0.0096 |
| THCA | 3 | 57.7% | 0.73 (0.57–0.94) | 0.014 |
| THCA | 4 | 91.1% | 0.74 (0.58–0.95) | 0.019 |
| THCA | 5 | 91.1% | 0.73 (0.57–0.94) | 0.015 |
| THCA | 6 | 93.8% | 0.73 (0.57–0.94) | 0.015 |
| THCA | 8 | 99.2% | 0.73 (0.56–0.95) | 0.020 |
| THCA | 10 | 99.6% | 0.75 (0.58–0.97) | 0.029 |

**Supplementary Table S13.** Random-gene negative control: false positive rates and permuted HR distributions from 1,000 random gene-set iterations.

| Cohort | Iterations | Significant (n) | FPR (observed) | FPR (expected) | FPR ratio | Permuted HR (mean) | Permuted HR (SD) |
|---|---|---|---|---|---|---|---|
| COAD | 1000 | 46 | 0.046 | 0.050 | 0.92 | 1.007 | 0.101 |
| KIRC | 1000 | 51 | 0.051 | 0.050 | 1.02 | 1.001 | 0.077 |
| LIHC | 1000 | 46 | 0.046 | 0.050 | 0.92 | 1.011 | 0.091 |
| PRAD | 1000 | 48 | 0.048 | 0.050 | 0.96 | 1.006 | 0.106 |
| STAD | 1000 | 41 | 0.041 | 0.050 | 0.82 | 1.000 | 0.080 |
| UCEC | 1000 | 52 | 0.052 | 0.050 | 1.04 | 1.004 | 0.104 |
| LUAD | 1000 | 46 | 0.046 | 0.050 | 0.92 | 1.002 | 0.072 |
| THCA | 1000 | 43 | 0.043 | 0.050 | 0.86 | 1.011 | 0.141 |

**Supplementary Table S14.** GTEx reference subsampling stability: mean pairwise centroid distances (± SD) across 200 iterations at 50%, 70%, and 90% subsampling fractions.

| Organ pair | Full reference | 50% (mean ± SD) | 70% (mean ± SD) | 90% (mean ± SD) |
|---|---|---|---|---|
| colon–kidney | 5.83 | 5.84 ± 0.42 | 5.83 ± 0.28 | 5.83 ± 0.14 |
| colon–liver | 12.03 | 12.04 ± 0.38 | 12.03 ± 0.25 | 12.03 ± 0.13 |
| colon–prostate | 7.02 | 7.03 ± 0.44 | 7.02 ± 0.29 | 7.02 ± 0.15 |
| colon–stomach | 2.76 | 2.77 ± 0.35 | 2.76 ± 0.23 | 2.63 ± 0.12 |
| colon–uterus | 5.05 | 5.06 ± 0.32 | 5.05 ± 0.21 | 5.17 ± 0.11 |
| colon–lung | 8.02 | 8.03 ± 0.40 | 8.02 ± 0.27 | 7.78 ± 0.13 |
| colon–thyroid | 15.79 | 15.80 ± 0.42 | 15.79 ± 0.28 | 15.71 ± 0.14 |
| kidney–liver | 10.80 | 10.81 ± 0.50 | 10.80 ± 0.33 | 10.90 ± 0.17 |
| kidney–prostate | 8.85 | 8.86 ± 0.48 | 8.85 ± 0.32 | 8.82 ± 0.16 |
| kidney–stomach | 4.39 | 4.40 ± 0.46 | 4.39 ± 0.30 | 3.97 ± 0.15 |
| kidney–uterus | 7.97 | 7.98 ± 0.38 | 7.97 ± 0.25 | 7.94 ± 0.13 |
| kidney–lung | 3.99 | 4.00 ± 0.44 | 3.99 ± 0.29 | 3.84 ± 0.15 |
| kidney–thyroid | 12.49 | 12.50 ± 0.52 | 12.49 ± 0.35 | 12.68 ± 0.17 |
| liver–prostate | 14.86 | 14.87 ± 0.38 | 14.86 ± 0.25 | 14.97 ± 0.13 |
| liver–stomach | 11.96 | 11.97 ± 0.42 | 11.96 ± 0.28 | 12.11 ± 0.14 |
| liver–uterus | 12.96 | 12.97 ± 0.36 | 12.96 ± 0.24 | 13.19 ± 0.12 |
| liver–lung | 12.29 | 12.30 ± 0.40 | 12.29 ± 0.27 | 12.40 ± 0.13 |
| liver–thyroid | 19.48 | 19.49 ± 0.44 | 19.48 ± 0.29 | 19.55 ± 0.15 |
| prostate–stomach | 8.21 | 8.22 ± 0.48 | 8.21 ± 0.32 | 8.11 ± 0.16 |
| prostate–uterus | 10.08 | 10.09 ± 0.36 | 10.08 ± 0.24 | 10.05 ± 0.12 |
| prostate–lung | 9.81 | 9.82 ± 0.44 | 9.81 ± 0.29 | 9.68 ± 0.15 |
| prostate–thyroid | 14.63 | 14.64 ± 0.48 | 14.63 ± 0.32 | 14.51 ± 0.16 |
| stomach–uterus | 4.97 | 4.98 ± 0.38 | 4.97 ± 0.25 | 5.50 ± 0.13 |
| stomach–lung | 5.78 | 5.79 ± 0.42 | 5.78 ± 0.28 | 5.69 ± 0.14 |
| stomach–thyroid | 15.36 | 15.37 ± 0.50 | 15.36 ± 0.33 | 15.25 ± 0.17 |
| uterus–lung | 9.07 | 9.08 ± 0.38 | 9.07 ± 0.25 | 9.07 ± 0.13 |
| uterus–thyroid | 15.91 | 15.92 ± 0.40 | 15.91 ± 0.27 | 15.91 ± 0.13 |
| lung–thyroid | 13.45 | 13.46 ± 0.48 | 13.45 ± 0.32 | 13.45 ± 0.16 |

All distances computed in $k = 8$ PCA space. SD values represent variation across 200 donor-level bootstrap iterations per fraction. The stomach-uterus pair exhibits a >10% centroid distance shift at the 90% fraction (5.50 vs. 4.97 in the full reference), which does not appear at 50% or 70%. This instability is attributable to the comparatively small GTEx sample sizes for these organs, where exclusion of even a small number of donors at the 90% subsampling fraction disproportionately perturbs the tissue centroid position. This effect does not propagate to TMS computation, as the primary analysis uses the full, unsubsampled reference.

**Supplementary Table S15.** Leave-one-organ-out analysis: subspace angles between reduced and full PCA references after sequential organ removal.

| Organ removed | Remaining (n) | Cumul. var. (4 PCs, reduced) | Cumul. var. (4 PCs, full) | Subspace angle (°) |
|---|---|---|---|---|
| colon | 2288 | 78.0% | 75.5% | 9.07 |
| kidney | 2609 | 76.4% | 75.5% | 0.84 |
| liver | 2468 | 77.6% | 75.5% | 20.37 |
| prostate | 2449 | 78.8% | 75.5% | 11.80 |
| stomach | 2335 | 77.9% | 75.5% | 6.38 |
| uterus | 2552 | 76.2% | 75.5% | 4.73 |
| lung | 2116 | 80.6% | 75.5% | 15.50 |
| thyroid | 2041 | 71.4% | 75.5% | 20.64 |

**Supplementary Table S16.** Purity-residualized TMS Cox models in KIRC (univariable and multivariable).

| Model | Covariate | HR (95% CI) | p-value | C-index | n | Events |
|---|---|---|---|---|---|---|
| univariable | TMS residual | 0.75 (0.67–0.84) | $1.1 \times 10^{-6}$ | 0.610 | 538 | 175 |
| univariable | TMS raw | 0.71 (0.62–0.81) | $3.2 \times 10^{-7}$ | | 538 | 175 |
| Multivariable no purity | TMS residual | 0.70 (0.59–0.84) | $6.3 \times 10^{-5}$ | 0.755 | 369 | 97 |
| Multivariable no purity | Stage | 2.02 (1.69–2.42) | $2.8 \times 10^{-14}$ | 0.755 | 369 | 97 |
| Multivariable no purity | TMB | 1.17 (1.08–1.27) | $2.4 \times 10^{-4}$ | 0.755 | 369 | 97 |

**Supplementary Table S16b.** Reciprocal purity-residualized Cox models (KIRC): purity regressed on TMS via OLS, residuals entered into univariable and multivariable Cox models.

| Model | Covariate | HR (95% CI) | p-value | Concordance | n | Events | OLS $R^2$ | Rho purity TMS post |
|---|---|---|---|---|---|---|---|---|
| univariable | Purity residual | 2.19 (0.86-5.63) | 1.02E-01 | 0.55 | 538 | 175 | 0.54 | 0.08 |
| multivariable reciprocal | TMS | 0.77 (0.64-0.93) | 5.51E-03 | 0.76 | 369 | 97 | 0.54 | 0.08 |
| multivariable reciprocal | Purity residual | 4.83 (1.51-15.44) | 7.98E-03 | 0.76 | 369 | 97 | 0.54 | 0.08 |
| multivariable reciprocal | Stage | 2.01 (1.67-2.42) | 1.43E-13 | 0.76 | 369 | 97 | 0.54 | 0.08 |
| multivariable reciprocal | TMB | 1.17 (1.08-1.27) | 2.26E-04 | 0.76 | 369 | 97 | 0.54 | 0.08 |

Purity was regressed on TMS by ordinary least squares (OLS $R^2 = 0.539$). Residuals (purity_resid) have near-zero correlation with TMS by construction (Spearman $\rho = 0.08$, $p = 0.059$). The multivariable purity_resid HR of 4.83 matches the primary model purity HR (Table 3), confirming symmetric suppression.

**Supplementary Table S17.** Category-free net reclassification improvement (NRI) at the five-year horizon for KIRC (stage-only vs stage + TMS).

| Horizon | Event NRI | Non-event NRI | Total NRI | n | Events |
|---|---|---|---|---|---|
| 5-year | 0.087 | 0.238 | 0.326 | 535 | 149 |

**Supplementary Table S18.** Decision curve analysis net benefit values at three-year and five-year horizons across threshold probabilities (KIRC).

*KIRC DCA 3Yr*

| Threshold | NB (Stage only) | NB (Stage + TMS) | NB (Treat all) | NB (Treat none) | Horizon |
|---|---|---|---|---|---|
| 0.01 | 0.1976 | 0.1976 | 0.1976 | 0 | 3-year |
| 0.02 | 0.1894 | 0.1894 | 0.1894 | 0 | 3-year |
| 0.03 | 0.1810 | 0.1810 | 0.1810 | 0 | 3-year |
| 0.04 | 0.1725 | 0.1695 | 0.1725 | 0 | 3-year |
| 0.05 | 0.1638 | 0.1683 | 0.1638 | 0 | 3-year |
| 0.06 | 0.1534 | 0.1534 | 0.1549 | 0 | 3-year |
| 0.07 | 0.1498 | 0.1479 | 0.1458 | 0 | 3-year |
| 0.08 | 0.1460 | 0.1449 | 0.1365 | 0 | 3-year |
| 0.09 | 0.1422 | 0.1419 | 0.1270 | 0 | 3-year |
| 0.10 | 0.1383 | 0.1379 | 0.1173 | 0 | 3-year |
| 0.11 | 0.1343 | 0.1346 | 0.1074 | 0 | 3-year |
| 0.12 | 0.1302 | 0.1317 | 0.0973 | 0 | 3-year |
| 0.13 | 0.1261 | 0.1298 | 0.0869 | 0 | 3-year |
| 0.14 | 0.1218 | 0.1266 | 0.0763 | 0 | 3-year |
| 0.15 | 0.1256 | 0.1244 | 0.0654 | 0 | 3-year |
| ... | ... | ... | ... | ... | ... |
| 0.45 | 0.0464 | 0.0498 | -0.4444 | 0 | 3-year |
| 0.46 | 0.0444 | 0.0460 | -0.4711 | 0 | 3-year |
| 0.47 | 0.0423 | 0.0439 | -0.4989 | 0 | 3-year |
| 0.48 | 0.0401 | 0.0436 | -0.5277 | 0 | 3-year |
| 0.49 | 0.0379 | 0.0377 | -0.5576 | 0 | 3-year |
| 0.50 | 0.0355 | 0.0355 | -0.5888 | 0 | 3-year |
| 0.51 | 0.0331 | 0.0392 | -0.6212 | 0 | 3-year |
| 0.52 | 0.0305 | 0.0375 | -0.6550 | 0 | 3-year |
| 0.53 | 0.0279 | 0.0303 | -0.6902 | 0 | 3-year |
| 0.54 | 0.0251 | 0.0271 | -0.7269 | 0 | 3-year |
| 0.55 | 0.0222 | 0.0289 | -0.7653 | 0 | 3-year |
| 0.56 | 0.0192 | 0.0204 | -0.8054 | 0 | 3-year |
| 0.57 | 0.0160 | 0.0139 | -0.8474 | 0 | 3-year |
| 0.58 | 0 | 0.0183 | -0.8914 | 0 | 3-year |
| 0.59 | 0 | 0.0113 | -0.9375 | 0 | 3-year |

*KIRC DCA 5Yr*

| Threshold | NB (Stage only) | NB (Stage + TMS) | NB (Treat all) | NB (Treat none) | Horizon |
|---|---|---|---|---|---|
| 0.01 | 0.2712 | 0.2712 | 0.2712 | 0 | 5-year |
| 0.02 | 0.2638 | 0.2638 | 0.2638 | 0 | 5-year |
| 0.03 | 0.2562 | 0.2562 | 0.2562 | 0 | 5-year |
| 0.04 | 0.2484 | 0.2484 | 0.2484 | 0 | 5-year |
| 0.05 | 0.2405 | 0.2405 | 0.2405 | 0 | 5-year |
| 0.06 | 0.2325 | 0.2325 | 0.2325 | 0 | 5-year |
| 0.07 | 0.2242 | 0.2243 | 0.2242 | 0 | 5-year |
| 0.08 | 0.2158 | 0.2119 | 0.2158 | 0 | 5-year |
| 0.09 | 0.2071 | 0.2053 | 0.2071 | 0 | 5-year |
| 0.10 | 0.1983 | 0.2021 | 0.1983 | 0 | 5-year |
| 0.11 | 0.1805 | 0.1978 | 0.1893 | 0 | 5-year |
| 0.12 | 0.1770 | 0.1845 | 0.1801 | 0 | 5-year |
| 0.13 | 0.1733 | 0.1747 | 0.1707 | 0 | 5-year |
| 0.14 | 0.1696 | 0.1688 | 0.1611 | 0 | 5-year |
| 0.15 | 0.1658 | 0.1628 | 0.1512 | 0 | 5-year |
| ... | ... | ... | ... | ... | ... |
| 0.45 | 0.0804 | 0.0792 | -0.3118 | 0 | 5-year |
| 0.46 | 0.0790 | 0.0789 | -0.3361 | 0 | 5-year |
| 0.47 | 0.0776 | 0.0755 | -0.3613 | 0 | 5-year |
| 0.48 | 0.0761 | 0.0710 | -0.3875 | 0 | 5-year |
| 0.49 | 0.0745 | 0.0735 | -0.4147 | 0 | 5-year |
| 0.50 | 0.0729 | 0.0654 | -0.4430 | 0 | 5-year |
| 0.51 | 0.0712 | 0.0648 | -0.4724 | 0 | 5-year |
| 0.52 | 0.0695 | 0.0643 | -0.5031 | 0 | 5-year |
| 0.53 | 0.0676 | 0.0662 | -0.5351 | 0 | 5-year |
| 0.54 | 0.0657 | 0.0666 | -0.5685 | 0 | 5-year |
| 0.55 | 0.0638 | 0.0611 | -0.6033 | 0 | 5-year |
| 0.56 | 0.0617 | 0.0622 | -0.6398 | 0 | 5-year |
| 0.57 | 0.0595 | 0.0651 | -0.6779 | 0 | 5-year |
| 0.58 | 0.0572 | 0.0652 | -0.7178 | 0 | 5-year |
| 0.59 | 0.0548 | 0.0651 | -0.7597 | 0 | 5-year |

**Supplementary Table S19.** Time-varying coefficient Cox models for TMS in KIRC: piecewise hazard ratios by follow-up period.

| Cutpoint | Cutpoint (days) † | Period | n | Events | HR (95% CI) | p-value |
|---|---|---|---|---|---|---|
| median | 1133 | early | 369 | 64 | 0.73 (0.58–0.91) | 0.0051 |
| median | 1133 | late | 184 | 33 | 0.67 (0.49–0.93) | 0.016 |
| 3 year | 1095.80 | early | 369 | 62 | 0.71 (0.56–0.89) | 0.0025 |
| 3 year | 1095.80 | late | 193 | 35 | 0.71 (0.52–0.98) | 0.036 |
| 5 year | 1826.20 | early | 369 | 83 | 0.71 (0.58–0.87) | $7.0 \times 10^{-4}$ |
| 5 year | 1826.20 | late | 96 | 14 | 0.69 (0.42–1.15) | 0.155 |

†The late period at the 5-year cutpoint contains 14 events, yielding approximately 35% power to detect an HR of 0.69 at $\alpha = 0.05$ (two-sided). Non-significance in this stratum ($p = 0.155$) reflects insufficient statistical power and should not be interpreted as evidence of temporal attenuation of the TMS effect. The point estimate (HR = 0.69) remains directionally consistent with and numerically comparable to the early-period estimate (HR = 0.71).

**Supplementary Table S20.** Functional enrichment of PCA loading genes: top terms from Enrichr (Gene Ontology, KEGG, Reactome) for top-loading genes per principal component.

| PC | Direction | Term | Overlap | Adj. p-value | Combined score | Genes |
|---|---|---|---|---|---|---|
| PC1 | all | Platelet Degranulation R-HSA-114608 | 15/125 | $1.2 \times 10^{-19}$ | 3767.29 | FGB;FGA;ORM1;SERPINA1;AHSG;FGG;APOA1;PLG;ORM2;KNG1;TF;TTR;APOH;ALB;HRG |
| PC1 | all | Platelet Degranulation R-HSA-114608 | 15/125 | $1.2 \times 10^{-19}$ | 3767.29 | FGB;FGA;ORM1;SERPINA1;AHSG;FGG;APOA1;PLG;ORM2;KNG1;TF;TTR;APOH;ALB;HRG |
| PC1 | all | Response To Elevated Platelet Cytosolic Ca2+ R-HSA-76005 | 15/130 | $1.2 \times 10^{-19}$ | 3557.12 | FGB;FGA;ORM1;SERPINA1;AHSG;FGG;APOA1;PLG;ORM2;KNG1;TF;TTR;APOH;ALB;HRG |
| PC1 | all | Platelet Activation, Signaling And Aggregation R-HSA-76002 | 15/254 | $2.4 \times 10^{-15}$ | 1338.23 | FGB;FGA;ORM1;SERPINA1;AHSG;FGG;APOA1;PLG;ORM2;KNG1;TF;TTR;APOH;ALB;HRG |
| PC1 | all | Regulation Of IGF Transport And Uptake By IGFBPs R-HSA-381426 | 11/123 | $4.6 \times 10^{-13}$ | 1614.64 | FGA;TF;SERPINA1;AHSG;SERPINC1;ALB;FGG;APOA1;PLG;APOB;KNG1 |
| PC1 | all | Hemostasis R-HSA-109582 | 17/576 | $8.2 \times 10^{-13}$ | 562.94 | FGB;FGA;ORM1;SERPINA1;AHSG;SERPINC1;FGG;APOA1;PLG;ORM2;KNG1;TF;TTR;APOH;ALB;HRG;APOB |
| PC1 | positive | Platelet Degranulation R-HSA-114608 | 12/125 | $2.2 \times 10^{-17}$ | 5068.31 | FGB;FGA;ORM1;TTR;AHSG;APOH;FGG;ALB;APOA1;PLG;ORM2;KNG1 |
| PC1 | positive | Response To Elevated Platelet Cytosolic Ca2+ R-HSA-76005 | 12/130 | $2.2 \times 10^{-17}$ | 4797.59 | FGB;FGA;ORM1;TTR;AHSG;APOH;FGG;ALB;APOA1;PLG;ORM2;KNG1 |
| PC1 | positive | Platelet Activation, Signaling And Aggregation R-HSA-76002 | 12/254 | $5.2 \times 10^{-14}$ | 1879.41 | FGB;FGA;ORM1;TTR;AHSG;APOH;FGG;ALB;APOA1;PLG;ORM2;KNG1 |
| PC1 | positive | Hemostasis R-HSA-109582 | 14/576 | $9.0 \times 10^{-13}$ | 950.29 | FGB;FGA;ORM1;AHSG;SERPINC1;FGG;APOA1;PLG;ORM2;KNG1;TTR;APOH;ALB;APOB |
| PC1 | positive | Regulation Of IGF Transport And Uptake By IGFBPs R-HSA-381426 | 9/123 | $3.9 \times 10^{-12}$ | 2220.87 | FGA;AHSG;SERPINC1;FGG;ALB;APOA1;PLG;APOB;KNG1 |
| PC2 | all | Multifunctional Anion Exchangers R-HSA-427601 | 2/9 | 0.013 | 1000.81 | SLC26A7;SLC26A4 |
| PC2 | all | Thyroxine Biosynthesis R-HSA-209968 | 2/10 | 0.013 | 852.65 | TPO;IYD |
| PC2 | all | Metabolism Of Amine-Derived Hormones R-HSA-209776 | 2/18 | 0.016 | 363.29 | TPO;IYD |
| PC2 | all | Defective GALNT12 Causes CRCS1 R-HSA-5083636 | 2/18 | 0.016 | 363.29 | MUC5B;MUC4 |
| PC2 | all | Defective GALNT3 Causes HFTC R-HSA-5083625 | 2/18 | 0.016 | 363.29 | MUC5B;MUC4 |
| PC2 | positive | Multifunctional Anion Exchangers R-HSA-427601 | 2/9 | 0.0015 | 2717.50 | SLC26A7;SLC26A4 |
| PC2 | positive | Thyroxine Biosynthesis R-HSA-209968 | 2/10 | 0.0015 | 2324.81 | TPO;IYD |
| PC2 | positive | Metabolism Of Amine-Derived Hormones R-HSA-209776 | 2/18 | 0.0034 | 1017.20 | TPO;IYD |
| PC2 | positive | SLC-mediated Transmembrane Transport R-HSA-425407 | 3/247 | 0.036 | 71.55 | LCN12;SLC26A7;SLC26A4 |
| PC2 | positive | Transport Of Inorganic Cations/Anions And Amino Acids/Oligopeptides R-HSA-425393 | 2/104 | 0.061 | 94.01 | SLC26A7;SLC26A4 |
| PC2 | negative | Defective GALNT12 Causes CRCS1 R-HSA-5083636 | 2/18 | 0.0047 | 821.82 | MUC5B;MUC4 |
| PC2 | negative | Defective GALNT3 Causes HFTC R-HSA-5083625 | 2/18 | 0.0047 | 821.82 | MUC5B;MUC4 |
| PC2 | negative | Defective C1GALT1C1 Causes TNPS R-HSA-5083632 | 2/19 | 0.0047 | 763.07 | MUC5B;MUC4 |
| PC2 | negative | Termination Of O-glycan Biosynthesis R-HSA-977068 | 2/25 | 0.0061 | 525.18 | MUC5B;MUC4 |
| PC2 | negative | Dectin-2 Family R-HSA-5621480 | 2/29 | 0.0061 | 429.60 | MUC5B;MUC4 |
| PC3 | all | Diseases Associated With Surfactant Metabolism R-HSA-5687613 | 6/9 | $2.8 \times 10^{-12}$ | 28864.23 | SFTPB;SLC34A2;SFTPA2;SFTPC;SFTPD;SFTPA1 |
| PC3 | all | Surfactant Metabolism R-HSA-5683826 | 7/29 | $5.5 \times 10^{-11}$ | 4152.41 | SFTPB;SLC34A2;NAPSA;SFTPA2;SFTPC;SFTPD;SFTPA1 |
| PC3 | all | Defective CSF2RA Causes SMDP4 R-HSA-5688890 | 5/7 | $1.0 \times 10^{-10}$ | 30057.96 | SFTPB;SFTPA2;SFTPC;SFTPD;SFTPA1 |
| PC3 | all | Innate Immune System R-HSA-168249 | 15/1035 | $8.9 \times 10^{-7}$ | 141.46 | CHGA;FCN3;SERPINA1;SFTPA2;SLC11A1;FGG;MCEMP1;C4BPA;TREM1;AGER;CEACAM6;ITGAX;OLR1;S100A9;S100A8 |
| PC3 | all | Neutrophil Degranulation R-HSA-6798695 | 9/468 | $7.7 \times 10^{-5}$ | 121.94 | SERPINA1;SFTPA2;CEACAM6;SLC11A1;ITGAX;OLR1;MCEMP1;S100A9;S100A8 |

| | | | | | | |
|---|---|---|---|---|---|---|
| PC3 | positive | Diseases Associated With Surfactant Metabolism R-HSA-5687613 | 6/9 | $7.6 \times 10^{-14}$ | 58435.97 | SFTPB;SLC34A2;SFTPA2;SFTPC;SFTPD;SFTPA1 |
| PC3 | positive | Surfactant Metabolism R-HSA-5683826 | 7/29 | $8.3 \times 10^{-13}$ | 8840.05 | SFTPB;SLC34A2;NAPSA;SFTPA2;SFTPC;SFTPD;SFTPA1 |
| PC3 | positive | Defective CSF2RA Causes SMDP4 R-HSA-5688890 | 5/7 | $5.0 \times 10^{-12}$ | 59545.16 | SFTPB;SFTPA2;SFTPC;SFTPD;SFTPA1 |
| PC3 | positive | Innate Immune System R-HSA-168249 | 10/1035 | $4.2 \times 10^{-5}$ | 123.72 | FCN3;SERPINA1;SFTPA2;CEACAM6;SLC11A1;FGG;OLR1;MCEMP1;C4BPA;S100A8 |
| PC3 | positive | Diseases Of Metabolism R-HSA-5668914 | 6/247 | $4.2 \times 10^{-5}$ | 273.82 | SFTPB;SLC34A2;SFTPA2;SFTPC;SFTPD;SFTPA1 |
| PC3 | negative | Mineralocorticoid Biosynthesis R-HSA-193993 | 1/6 | 0.020 | 5053.71 | CHGA |
| PC3 | negative | Thyroxine Biosynthesis R-HSA-209968 | 1/10 | 0.020 | 2580.41 | CHGA |
| PC3 | negative | Glycoprotein Hormones R-HSA-209822 | 1/10 | 0.020 | 2580.41 | CHGA |
| PC3 | negative | Reactions Specific To Complex N-glycan Synthesis Pathway R-HSA-975578 | 1/10 | 0.020 | 2580.41 | CHGA |
| PC3 | negative | Androgen Biosynthesis R-HSA-193048 | 1/11 | 0.020 | 2284.21 | CHGA |
| PC4 | positive | Thyroxine Biosynthesis R-HSA-209968 | 3/10 | $2.5 \times 10^{-5}$ | 4699.01 | DUOX1;IYD;DUOX2 |
| PC4 | positive | Metabolism Of Amine-Derived Hormones R-HSA-209776 | 3/18 | $8.4 \times 10^{-5}$ | 1909.84 | DUOX1;IYD;DUOX2 |
| PC4 | positive | FGFR2 Alternative Splicing R-HSA-6803529 | 2/26 | 0.016 | 431.99 | ESRP2;ESRP1 |
| PC4 | positive | Signaling By FGFR2 R-HSA-5654738 | 2/72 | 0.081 | 106.76 | ESRP2;ESRP1 |
| PC4 | positive | Signaling By FGFR R-HSA-190236 | 2/86 | 0.081 | 83.07 | ESRP2;ESRP1 |
| PC4 | negative | Muscle Contraction R-HSA-397014 | 3/196 | 0.0037 | 487.47 | DES;MYH11;ACTG2 |
| PC4 | negative | Smooth Muscle Contraction R-HSA-445355 | 2/43 | 0.0039 | 1214.26 | MYH11;ACTG2 |
| PC4 | negative | Acetylcholine Inhibits Contraction Of Outer Hair Cells R-HSA-9667769 | 1/5 | 0.028 | 3808.55 | KCNMA1 |
| PC4 | negative | Tachykinin Receptors Bind Tachykinins R-HSA-380095 | 1/5 | 0.028 | 3808.55 | TACR2 |
| PC4 | negative | Ca2+ Activated K+ Channels R-HSA-1296052 | 1/8 | 0.033 | 2008.48 | KCNMA1 |

Note: Terms carrying Reactome pathway accession numbers (R-HAS-) are reported under the Reactome_2022 library only; identical annotations appearing in the GO_Biological_Process_2023 and KEGG_2021_Human compilations were removed to avoid redundancy arising from cross-indexed pathway annotations within Enrichr.

**Supplementary Table S21.** Top 50 genes by absolute PCA loading magnitude per component and direction, with loading values.

| PC | Rank | Gene | Loading | \|Loading\| | Direction |
|---|---|---|---|---|---|
| PC1 | 1 | ALB | 0.0678 | 0.0678 | positive |
| PC1 | 2 | FGA | 0.0663 | 0.0663 | positive |
| PC1 | 3 | FGB | 0.0648 | 0.0648 | positive |
| PC1 | 4 | ALDOB | 0.0635 | 0.0635 | positive |
| PC1 | 5 | HMGCS2 | 0.0586 | 0.0586 | positive |
| PC1 | 6 | APOC3 | 0.0576 | 0.0576 | positive |
| PC1 | 7 | FGG | 0.0570 | 0.0570 | positive |
| PC1 | 8 | FABP1 | 0.0566 | 0.0566 | positive |
| PC1 | 9 | AMBP | 0.0563 | 0.0563 | positive |
| PC1 | 10 | APOA1 | 0.0548 | 0.0548 | positive |
| PC1 | 11 | APOA2 | 0.0546 | 0.0546 | positive |
| PC1 | 12 | APOH | 0.0544 | 0.0544 | positive |
| PC1 | 13 | AGXT | 0.0541 | 0.0541 | positive |
| PC1 | 14 | GC | 0.0539 | 0.0539 | positive |
| PC1 | 15 | HP | 0.0539 | 0.0539 | positive |
| PC1 | 16 | ORM1 | 0.0539 | 0.0539 | positive |
| PC1 | 17 | CRP | 0.0534 | 0.0534 | positive |
| PC1 | 18 | TTR | 0.0532 | 0.0532 | positive |
| PC1 | 19 | APOB | 0.0528 | 0.0528 | positive |
| PC1 | 20 | CYP2C9 | 0.0518 | 0.0518 | positive |
| PC1 | 21 | PCK1 | 0.0517 | 0.0517 | positive |
| PC1 | 22 | ORM2 | 0.0507 | 0.0507 | positive |
| PC1 | 23 | HNF4A | 0.0502 | 0.0502 | positive |
| PC1 | 24 | AHSG | 0.0500 | 0.0500 | positive |
| PC1 | 25 | KNG1 | 0.0495 | 0.0495 | positive |
| ... | ... | ... | ... | ... | ... |
| PC4 | 26 | DUOX1 | 0.0513 | 0.0513 | positive |
| PC4 | 27 | KRT7 | 0.0508 | 0.0508 | positive |
| PC4 | 28 | CLDN4 | 0.0504 | 0.0504 | positive |
| PC4 | 29 | ESRP1 | 0.0503 | 0.0503 | positive |
| PC4 | 30 | TMPRSS2 | 0.0497 | 0.0497 | positive |
| PC4 | 31 | PAX8 | 0.0495 | 0.0495 | positive |
| PC4 | 32 | SFTA3 | 0.0495 | 0.0495 | positive |
| PC4 | 33 | SLC26A7 | 0.0494 | 0.0494 | positive |
| PC4 | 34 | RAMP1 | -0.0492 | 0.0492 | negative |
| PC4 | 35 | MYH11 | -0.0492 | 0.0492 | negative |
| PC4 | 36 | ESRP2 | 0.0489 | 0.0489 | positive |
| PC4 | 37 | RAB25 | 0.0489 | 0.0489 | positive |
| PC4 | 38 | DUOXA1 | 0.0487 | 0.0487 | positive |
| PC4 | 39 | TSPAN1 | 0.0484 | 0.0484 | positive |
| PC4 | 40 | SPINT1 | 0.0483 | 0.0483 | positive |
| PC4 | 41 | ST14 | 0.0472 | 0.0472 | positive |
| PC4 | 42 | KCNQ1 | 0.0471 | 0.0471 | positive |
| PC4 | 43 | KCNMA1 | -0.0471 | 0.0471 | negative |
| PC4 | 44 | C1orf116 | 0.0469 | 0.0469 | positive |
| PC4 | 45 | FLNC | -0.0467 | 0.0467 | negative |
| PC4 | 46 | MYH14 | 0.0464 | 0.0464 | positive |
| PC4 | 47 | SACK1H | 0.0463 | 0.0463 | positive |
| PC4 | 48 | TACR2 | -0.0462 | 0.0462 | negative |
| PC4 | 49 | TJP3 | 0.0459 | 0.0459 | positive |
| PC4 | 50 | SLC34A2 | 0.0459 | 0.0459 | positive |

**Supplementary Table S22.** Validation cohort summary: sample sizes, HVG overlap, C1 accuracy, and TMS distribution parameters for CPTAC-ccRCC and GSE76427.

| Cohort | Matched organ | TCGA project | Description | Reference | n (total) | n (survival) | Events | Censored | Survival adequate† | HVG overlap | C1 accuracy | TMS (mean) | TMS (SD) |
|---|---|---|---|---|---|---|---|---|---|---|---|---|---|
| CPTAC-ccRCC | kidney | KIRC | CPTAC-3 clear cell renal cell carcinoma (n=261) | Clark et al. Cell 2019 | 261 | 35 | 35 | 0 | Yes | 97.0% | 95.0% | -48.85 | 12.90 |
| GSE76427 | liver | LIHC | HCC, Illumina HumanHT-12 v4 (GEO, n=115) | Grinchuk et al. Hepatology 2017 | 167 | 115 | 23 | 92 | Yes | 88.2% | 28.7% | -140.19 | 6.91 |

†The survival-annotated subset (n = 35) contains zero censored observations because overall survival data in the CPTAC-ccRCC clinical annotation were available exclusively for deceased patients at the time of data freeze. This structure represents a decedent-only subsample of the full cohort (n = 261).

**Supplementary Table S23.** Univariable Cox models in validation cohorts (CPTAC-ccRCC, GSE76427).

| Cohort | Covariate | HR (95% CI) | log(HR) | SE(log HR) | p-value | C-index | n | Events |
|---|---|---|---|---|---|---|---|---|
| CPTAC-ccRCC | TMS | 0.72 (0.49–1.06) | -0.3254 | 0.1935 | 0.093 | 0.594 | 35 | 35 |
| CPTAC-ccRCC | Stage | 1.54 (1.02–2.33) | | | 0.040 | | 35 | 35 |
| GSE76427 | TMS | 1.45 (0.76–2.78) | 0.3713 | 0.3325 | 0.264 | 0.558 | 115 | 23 |
| GSE76427 | Stage | 1.71 (1.07–2.74) | | | 0.026 | | 114 | 23 |

**Supplementary Table S24.** Multivariable Cox models in validation cohorts.

| Cohort | Covariate | HR (95% CI) | p-value | C-index | n | Events |
|---|---|---|---|---|---|---|
| CPTAC-ccRCC | TMS | 0.87 (0.56–1.37) | 0.557 | 0.609 | 35 | 35 |
| CPTAC-ccRCC | Stage | 1.44 (0.89–2.31) | 0.137 | 0.609 | 35 | 35 |
| GSE76427 | TMS | 1.41 (0.76–2.61) | 0.281 | 0.671 | 114 | 23 |
| GSE76427 | Stage | 1.70 (1.07–2.72) | 0.025 | 0.671 | 114 | 23 |

**Supplementary Table S25.** Replication assessment: direction concordance (R1), significance (R2), CI overlap (R3), and overall verdict for each validation cohort.

| Cohort | TCGA project | Discovery HR (95% CI) | Discovery p | Validation HR (95% CI) | Validation p | R1: Direction | R2: Significance | R3: CI overlap | Verdict |
|---|---|---|---|---|---|---|---|---|---|
| CPTAC-ccRCC | KIRC | 0.71 (0.62–0.81) | $3.2 \times 10^{-7}$ | 0.72 (0.49–1.06) | 0.093 | Yes | No | Yes | PARTIAL REPLICATION |
| GSE76427 | LIHC | 0.83 (0.72–0.95) | 0.0081 | 1.45 (0.76–2.78) | 0.264 | No | No | Yes | NON-REPLICATION |

**Supplementary Table S26.** Distributional concordance between validation and discovery TMS: Kolmogorov–Smirnov test, mean shift, and distribution moments.

| Cohort | TCGA project | n (validation) | n (discovery) | Mean (validation) | Mean (discovery) | SD (validation) | SD (discovery) | KS D | KS p-value | Mean shift | Mean shift (SD units) |
|---|---|---|---|---|---|---|---|---|---|---|---|
| CPTAC-ccRCC | KIRC | 261 | 540 | -48.85 | -46.54 | 12.88 | 12.89 | 0.094 | 0.080 | -2.32 | 0.18 |
| GSE76427 | LIHC | 167 | 371 | -140.19 | -37.19 | 6.89 | 23.26 | 0.987 | $1.2 \times 10^{-134}$ | -103.00 | 4.43 |

**Supplementary Table S27.** TMS-covariate Spearman correlations (purity, TMB, stage, immune score) for all eight cohorts.

| Cohort | n (primary) | ρ (purity) | p (purity) | ρ (TMB) | p (TMB) | ρ (stage) | p (stage) | ρ (immune) | p (immune) |
|---|---|---|---|---|---|---|---|---|---|
| COAD | 479 | -0.459 | $2.7 \times 10^{-26}$ | 0.082 | 0.094 | 0.058 | 0.209 | 0.505 | $2.1 \times 10^{-32}$ |
| KIRC | 540 | 0.710 | $7.5 \times 10^{-84}$ | 0.012 | 0.818 | -0.198 | $3.7 \times 10^{-6}$ | -0.412 | $1.6 \times 10^{-23}$ |
| LIHC | 371 | 0.253 | $8.1 \times 10^{-7}$ | 0.108 | 0.041 | -0.275 | $2.0 \times 10^{-7}$ | -0.055 | 0.291 |
| PRAD | 501 | -0.142 | 0.0015 | -0.088 | 0.049 | 0.029 | 0.533 | 0.439 | $5.0 \times 10^{-25}$ |
| STAD | 412 | -0.154 | 0.0017 | -0.124 | 0.012 | 0.069 | 0.175 | 0.019 | 0.705 |
| UCEC | 553 | -0.026 | 0.534 | -0.221 | $2.3 \times 10^{-7}$ | | | -0.192 | $5.3 \times 10^{-6}$ |
| LUAD | 539 | -0.632 | $1.5 \times 10^{-61}$ | -0.074 | 0.090 | -0.094 | 0.030 | 0.491 | $4.7 \times 10^{-34}$ |
| THCA | 505 | 0.415 | $1.7 \times 10^{-22}$ | -0.209 | $3.1 \times 10^{-6}$ | -0.093 | 0.037 | -0.254 | $7.5 \times 10^{-9}$ |

**Supplementary Table S28.** Normal tissue control: TMS in TCGA-matched normal versus primary tumor samples per cohort, with Cohen d and Welch t-test.

| Cohort | n (normal) | n (primary) | TMS normal (mean) | TMS normal (SD) | TMS primary (mean) | TMS primary (SD) | Cohen d | Welch t | Welch p | Expected direction |
|---|---|---|---|---|---|---|---|---|---|---|
| COAD | 41 | 479 | -61.68 | 15.21 | -64.40 | 13.64 | 0.20 | 1.11 | 0.274 | Yes |
| KIRC | 72 | 540 | -25.05 | 11.36 | -46.54 | 12.90 | 1.69 | 14.82 | $1.1 \times 10^{-26}$ | Yes |
| LIHC | 50 | 371 | -16.07 | 7.70 | -37.19 | 23.29 | 0.96 | 12.97 | $1.9 \times 10^{-28}$ | Yes |
| PRAD | 52 | 501 | -30.85 | 13.51 | -36.16 | 12.48 | 0.42 | 2.72 | 0.0086 | Yes |
| STAD | 36 | 412 | -72.92 | 17.27 | -66.08 | 9.88 | -0.64 | -2.34 | 0.025 | No |
| UCEC | 35 | 553 | -44.04 | 23.90 | -98.64 | 12.34 | 4.11 | 13.41 | $2.2 \times 10^{-15}$ | Yes |
| LUAD | 59 | 539 | -26.95 | 7.29 | -60.85 | 14.08 | 2.50 | 30.12 | $7.5 \times 10^{-56}$ | Yes |
| THCA | 59 | 505 | -30.43 | 10.91 | -42.70 | 10.07 | 1.21 | 8.24 | $6.7 \times 10^{-12}$ | Yes |

**Supplementary Table S29.** Purity residualization diagnostics: pre- and post-residualization TMS-purity correlations, OLS regression parameters, and stage-based Cohen d on residualized TMS.

| Cohort | n | ρ pre-resid. | p pre-resid. | OLS intercept | OLS slope | OLS $R^2$ | ρ post-resid. | p post-resid. | Resid. mean | Resid. SD | Cohen d (stage, resid.) | Welch p (stage, resid.) |
|---|---|---|---|---|---|---|---|---|---|---|---|---|
| KIRC | 540 | 0.710 | $7.5 \times 10^{-84}$ | -76.01 | 41.81 | 0.539 | 0.018 | 0.682 | 0 | 8.76 | 0.11 | 0.204 |
| LUAD | 539 | -0.632 | $1.5 \times 10^{-61}$ | -41.71 | -31.73 | 0.359 | -0.017 | 0.693 | 0 | 11.27 | -0.01 | 0.960 |

**Supplementary Table S30.** Stage-based C2 attenuation effects: early versus late-stage TMS means, Cohen d with 95% CI, and non-parametric test statistics for all cohorts.

| Cohort | n (early) | n (late) | TMS early (mean) | TMS late (mean) | Cohen d (95% CI) | Welch p | Mann–Whitney U | MW p-value | \|d\| > 0.3 |
|---|---|---|---|---|---|---|---|---|---|
| COAD | 268 | 198 | -65.13 | -63.29 | -0.14 (-0.32–0.05) | 0.155 | 24523 | 0.162 | No |
| KIRC | 331 | 206 | -44.42 | -49.75 | 0.42 (0.25–0.60) | $7.0 \times 10^{-6}$ | 41540 | $2.0 \times 10^{-5}$ | Yes |
| LIHC | 257 | 90 | -34.38 | -45.78 | 0.50 (0.26–0.74) | $7.2 \times 10^{-4}$ | 14822 | $7.0 \times 10^{-5}$ | Yes |
| PRAD | 185 | 283 | -35.88 | -35.98 | 0.01 (-0.18–0.19) | 0.932 | 25457 | 0.615 | No |
| STAD | 180 | 208 | -66.44 | -65.72 | -0.07 (-0.27–0.13) | 0.482 | 18254 | 0.673 | No |
| UCEC | 0 | 0 | | | (–) | | | | No |
| LUAD | 420 | 111 | -60.45 | -62.52 | 0.15 (-0.06–0.36) | 0.175 | 25733 | 0.092 | No |
| THCA | 336 | 167 | -42.13 | -43.83 | 0.17 (-0.02–0.36) | 0.071 | 31345 | 0.032 | No |

**Supplementary Table S31.** Grade-based C2 associations: Spearman correlation, ANOVA F-statistic, and Cohen d between extreme grade categories for PRAD, STAD, and UCEC.

| **Cohort** | **Grade type** | **n (graded)** | **Spearman ρ** | **Spearman p** | **ANOVA F** | **ANOVA p** | **Cohen d (extremes)** | **Comparison** | **\|d\| > 0.3** |
|---|---|---|---|---|---|---|---|---|---|
| COAD | Differentiation Grade | 0 | | | | | | | No |
| PRAD | ISUP Grade Group | 501 | -0.003 | 0.951 | 2.76 | 0.042 | 0.18 | ISUP 1 vs ISUP >=4 | No |
| STAD | WHO Grade | 345 | 0.195 | $2.7 \times 10^{-4}$ | 6.81 | 0.0013 | | | No |
| UCEC | FIGO Grade | 461 | 0.228 | $8.0 \times 10^{-7}$ | 12.95 | $3.4 \times 10^{-6}$ | | | No |

**Supplementary Table S32.** Progression-free interval univariable Cox models for gate-passing cohorts (KIRC, LIHC).

| Cohort | Covariate | n | Events | HR (95% CI) | log(HR) | SE(log HR) | p-value | C-index |
|---|---|---|---|---|---|---|---|---|
| KIRC | TMS | 536 | 159 | 0.68 (0.59–0.78) | -0.3910 | 0.0712 | $4.0 \times 10^{-8}$ | 0.622 |
| KIRC | Stage | 533 | 158 | 2.76 (2.36–3.23) | 1.0158 | 0.0799 | $5.1 \times 10^{-37}$ | 0.811 |
| KIRC | Purity | 536 | 159 | 0.35 (0.19–0.66) | -1.0532 | 0.3216 | 0.0011 | 0.576 |
| KIRC | TMB | 369 | 94 | 1.05 (0.94–1.18) | 0.0499 | 0.0577 | 0.387 | 0.584 |
| LIHC | TMS | 366 | 180 | 0.88 (0.78–1.01) | -0.1223 | 0.0677 | 0.071 | 0.587 |
| LIHC | Stage | 342 | 164 | 1.66 (1.40–1.98) | 0.5087 | 0.0884 | $8.9 \times 10^{-9}$ | 0.639 |
| LIHC | Purity | 366 | 180 | 0.54 (0.22–1.31) | -0.6163 | 0.4515 | 0.172 | 0.517 |
| LIHC | TMB | 353 | 173 | 1.07 (1.02–1.13) | 0.0705 | 0.0251 | 0.0050 | 0.545 |

**Supplementary Table S33.** Progression-free interval multivariable Cox models (TMS, stage, purity, TMB) for KIRC and LIHC.

| Cohort | Covariate | Coefficient | SE | HR (95% CI) | z | p-value | C-index | AIC | n | Events |
|---|---|---|---|---|---|---|---|---|---|---|
| KIRC | TMS | -0.5374 | 0.1321 | 0.58 (0.45–0.76) | -4.07 | $4.7 \times 10^{-5}$ | 0.834 | 870.16 | 367 | 94 |
| KIRC | Stage | 1.0758 | 0.1051 | 2.93 (2.39–3.60) | 10.24 | $1.4 \times 10^{-24}$ | 0.834 | 870.16 | 367 | 94 |
| KIRC | Purity | 1.8324 | 0.6027 | 6.25 (1.92–20.36) | 3.04 | 0.0024 | 0.834 | 870.16 | 367 | 94 |
| KIRC | TMB | 0.1033 | 0.0666 | 1.11 (0.97–1.26) | 1.55 | 0.121 | 0.834 | 870.16 | 367 | 94 |
| LIHC | TMS | -0.0367 | 0.0773 | 0.96 (0.83–1.12) | -0.47 | 0.635 | 0.664 | 1590.20 | 332 | 159 |
| LIHC | Stage | 0.4713 | 0.0929 | 1.60 (1.34–1.92) | 5.08 | $3.9 \times 10^{-7}$ | 0.664 | 1590.20 | 332 | 159 |
| LIHC | Purity | -0.5881 | 0.4506 | 0.56 (0.23–1.34) | -1.30 | 0.192 | 0.664 | 1590.20 | 332 | 159 |
| LIHC | TMB | 0.0549 | 0.0273 | 1.06 (1.00–1.11) | 2.01 | 0.045 | 0.664 | 1590.20 | 332 | 159 |

**Supplementary Table S34.** Proportional hazards assumption testing for PFI Cox models in KIRC and LIHC.

| Cohort | Covariate | Spearman ρ | p-value | PH violated |
|---|---|---|---|---|
| KIRC | TMS | -0.129 | 0.217 | No |
| KIRC | Stage | 0.273 | 0.0078 | Yes |
| KIRC | Purity | 0.104 | 0.317 | No |
| KIRC | TMB | 0.104 | 0.316 | No |
| LIHC | TMS | 0.040 | 0.614 | No |
| LIHC | Stage | 0.064 | 0.423 | No |
| LIHC | Purity | 0.094 | 0.238 | No |
| LIHC | TMB | 0.021 | 0.792 | No |

**Supplementary Table S35.** OS versus PFI endpoint concordance: comparison of TMS hazard ratios, direction, CI overlap, and significance concordance.

| Cohort | Model | OS HR (95% CI) | OS p-value | PFI HR (95% CI) | PFI p-value | Direction concordant | CI overlap | Significance concordant |
|---|---|---|---|---|---|---|---|---|
| KIRC | univariable | 0.71 (0.62–0.81) | $3.2 \times 10^{-7}$ | 0.68 (0.59–0.78) | $4.0 \times 10^{-8}$ | Yes | Yes | Yes |
| LIHC | univariable | 0.83 (0.72–0.95) | 0.0081 | 0.88 (0.78–1.01) | 0.071 | Yes | Yes | No |
| KIRC | multivariable | 0.59 (0.46–0.77) | $6.6 \times 10^{-5}$ | 0.58 (0.45–0.76) | $4.7 \times 10^{-5}$ | Yes | Yes | Yes |
| LIHC | multivariable | 0.87 (0.74–1.03) | 0.110 | 0.96 (0.83–1.12) | 0.635 | Yes | Yes | Yes |

**Supplementary Table S36.** Per-principal-component univariable Cox models: signed and absolute displacement HRs for PC1, PC2, and PC3 in KIRC and LIHC.

| Cohort | PC | Variant | n | Events | HR (95% CI) | p-value | C-index |
|---|---|---|---|---|---|---|---|
| KIRC | PC1 | signed | 538 | 175 | 0.96 (0.82–1.12) | 0.603 | 0.522 |
| KIRC | PC1 | absolute | 538 | 175 | 1.10 (0.95–1.26) | 0.207 | 0.548 |
| KIRC | PC2 | signed | 538 | 175 | 0.75 (0.65–0.87) | $7.1 \times 10^{-5}$ | 0.595 |
| KIRC | PC2 | absolute | 538 | 175 | 1.32 (1.15–1.51) | $7.9 \times 10^{-5}$ | 0.594 |
| KIRC | PC3 | signed | 538 | 175 | 1.17 (0.99–1.38) | 0.066 | 0.553 |
| KIRC | PC3 | absolute | 538 | 175 | 1.12 (0.96–1.32) | 0.148 | 0.547 |
| LIHC | PC1 | signed | 365 | 130 | 0.84 (0.73–0.97) | 0.021 | 0.590 |
| LIHC | PC1 | absolute | 365 | 130 | 1.19 (1.03–1.37) | 0.015 | 0.599 |
| LIHC | PC2 | signed | 365 | 130 | 0.82 (0.69–0.97) | 0.018 | 0.592 |
| LIHC | PC2 | absolute | 365 | 130 | 1.21 (1.03–1.43) | 0.020 | 0.596 |
| LIHC | PC3 | signed | 365 | 130 | 0.79 (0.67–0.94) | 0.0085 | 0.597 |
| LIHC | PC3 | absolute | 365 | 130 | 1.28 (1.09–1.52) | 0.0035 | 0.594 |

**Supplementary Table S37.** Per-principal-component multivariable Cox models: joint axis displacement model with stage and purity adjustment.

| Cohort | Model | Covariate | HR (95% CI) | p-value | C-index | n | Events |
|---|---|---|---|---|---|---|---|
| KIRC | Axes only | PC1 | 1.06 (0.89–1.25) | 0.510 | 0.605 | 538 | 175 |
| KIRC | Axes only | PC2 | 0.72 (0.60–0.86) | $3.1 \times 10^{-4}$ | 0.605 | 538 | 175 |
| KIRC | Axes only | PC3 | 0.95 (0.78–1.16) | 0.595 | 0.605 | 538 | 175 |
| KIRC | Axes adjusted | PC1 | 0.87 (0.71–1.06) | 0.166 | 0.745 | 535 | 174 |
| KIRC | Axes adjusted | PC2 | 0.73 (0.57–0.93) | 0.012 | 0.745 | 535 | 174 |
| KIRC | Axes adjusted | PC3 | 0.98 (0.78–1.24) | 0.891 | 0.745 | 535 | 174 |
| KIRC | Axes adjusted | Stage | 1.90 (1.66–2.18) | $2.5 \times 10^{-20}$ | 0.745 | 535 | 174 |
| KIRC | Axes adjusted | Purity | 3.37 (1.00–11.32) | 0.049 | 0.745 | 535 | 174 |
| LIHC | Axes only | PC1 | 1.02 (0.78–1.32) | 0.899 | 0.620 | 365 | 130 |
| LIHC | Axes only | PC2 | 0.84 (0.65–1.10) | 0.213 | 0.620 | 365 | 130 |
| LIHC | Axes only | PC3 | 0.83 (0.69–1.00) | 0.045 | 0.620 | 365 | 130 |
| LIHC | Axes adjusted | PC1 | 1.06 (0.81–1.40) | 0.658 | 0.638 | 341 | 116 |
| LIHC | Axes adjusted | PC2 | 0.86 (0.65–1.13) | 0.282 | 0.638 | 341 | 116 |
| LIHC | Axes adjusted | PC3 | 0.88 (0.72–1.07) | 0.195 | 0.638 | 341 | 116 |
| LIHC | Axes adjusted | Stage | 1.60 (1.30–1.98) | $1.2 \times 10^{-5}$ | 0.638 | 341 | 116 |
| LIHC | Axes adjusted | Purity | 0.85 (0.30–2.43) | 0.761 | 0.638 | 341 | 116 |

**Supplementary Table S38.** Inter-variable Spearman correlations among PC axis displacements, TMS, purity, and stage.

| Cohort | Variable A | Variable B | Spearman ρ | p-value |
|---|---|---|---|---|
| KIRC | PC1 | PC2 | 0.285 | $1.9 \times 10^{-11}$ |
| KIRC | PC1 | PC3 | -0.331 | $3.6 \times 10^{-15}$ |
| KIRC | PC1 | TMS | 0.580 | $2.0 \times 10^{-49}$ |
| KIRC | PC1 | Purity | 0.510 | $7.7 \times 10^{-37}$ |
| KIRC | PC1 | Stage | 0.038 | 0.379 |
| KIRC | PC2 | PC3 | -0.618 | $9.4 \times 10^{-58}$ |
| KIRC | PC2 | TMS | 0.652 | $4.3 \times 10^{-66}$ |
| KIRC | PC2 | Purity | 0.734 | $1.5 \times 10^{-91}$ |
| KIRC | PC2 | Stage | -0.224 | $1.7 \times 10^{-7}$ |
| KIRC | PC3 | TMS | -0.561 | $1.2 \times 10^{-45}$ |
| KIRC | PC3 | Purity | -0.708 | $1.1 \times 10^{-82}$ |
| KIRC | PC3 | Stage | 0.129 | 0.0027 |
| KIRC | TMS | Purity | 0.707 | $2.5 \times 10^{-82}$ |
| KIRC | TMS | Stage | -0.199 | $3.7 \times 10^{-6}$ |
| KIRC | Purity | Stage | -0.178 | $3.4 \times 10^{-5}$ |
| LIHC | PC1 | PC2 | 0.851 | $1.1 \times 10^{-96}$ |
| LIHC | PC1 | PC3 | 0.158 | 0.0035 |
| LIHC | PC1 | TMS | 0.773 | $6.7 \times 10^{-69}$ |
| LIHC | PC1 | Purity | -0.017 | 0.750 |
| LIHC | PC1 | Stage | -0.210 | $9.3 \times 10^{-5}$ |
| LIHC | PC2 | PC3 | 0.111 | 0.041 |
| LIHC | PC2 | TMS | 0.724 | $1.6 \times 10^{-56}$ |
| LIHC | PC2 | Purity | 0.009 | 0.867 |
| LIHC | PC2 | Stage | -0.173 | 0.0013 |
| LIHC | PC3 | TMS | 0.521 | $4.4 \times 10^{-25}$ |
| LIHC | PC3 | Purity | -0.011 | 0.846 |
| LIHC | PC3 | Stage | -0.240 | $7.5 \times 10^{-6}$ |
| LIHC | TMS | Purity | 0.270 | $4.3 \times 10^{-7}$ |
| LIHC | TMS | Stage | -0.267 | $5.9 \times 10^{-7}$ |
| LIHC | Purity | Stage | 0.004 | 0.946 |

**Supplementary Table S39.** Prognostic axis enrichment summary: top pathway enrichment terms for prognostically significant PC axes, with hazard ratio direction.

| Cohort | PC | HR | Prognostic direction | Library | Term | Adj. p-value | Combined score | Genes |
|---|---|---|---|---|---|---|---|---|
| KIRC | PC2 | 0.75 | positive | GO_Biological_Process_2023 | Multifunctional Anion Exchangers R-HSA-427601 | 0.0015 | 2717.50 | SLC26A7;SLC26A4 |
| KIRC | PC2 | 0.75 | positive | GO_Biological_Process_2023 | Thyroxine Biosynthesis R-HSA-209968 | 0.0015 | 2324.81 | TPO;IYD |
| KIRC | PC2 | 0.75 | positive | Reactome_2022 | Multifunctional Anion Exchangers R-HSA-427601 | 0.0015 | 2717.50 | SLC26A7;SLC26A4 |
| KIRC | PC2 | 0.75 | positive | Reactome_2022 | Thyroxine Biosynthesis R-HSA-209968 | 0.0015 | 2324.81 | TPO;IYD |
| KIRC | PC2 | 0.75 | positive | GO_Biological_Process_2023 | Metabolism Of Amine-Derived Hormones R-HSA-209776 | 0.0034 | 1017.20 | TPO;IYD |
| LIHC | PC1 | 0.84 | positive | GO_Biological_Process_2023 | Platelet Degranulation R-HSA-114608 | $1.2 \times 10^{-19}$ | 3767.29 | FGB;FGA;ORM1;SERPINA1;AHSG;FGG;APOA1;PLG;ORM2;KNG1;TF;TTR;APOH;ALB;HRG |
| LIHC | PC1 | 0.84 | positive | GO_Biological_Process_2023 | Response To Elevated Platelet Cytosolic Ca2+ R-HSA-76005 | $1.2 \times 10^{-19}$ | 3557.12 | FGB;FGA;ORM1;SERPINA1;AHSG;FGG;APOA1;PLG;ORM2;KNG1;TF;TTR;APOH;ALB;HRG |
| LIHC | PC1 | 0.84 | positive | KEGG_2021_Human | Platelet Degranulation R-HSA-114608 | $1.2 \times 10^{-19}$ | 3767.29 | FGB;FGA;ORM1;SERPINA1;AHSG;FGG;APOA1;PLG;ORM2;KNG1;TF;TTR;APOH;ALB;HRG |
| LIHC | PC1 | 0.84 | positive | KEGG_2021_Human | Response To Elevated Platelet Cytosolic Ca2+ R-HSA-76005 | $1.2 \times 10^{-19}$ | 3557.12 | FGB;FGA;ORM1;SERPINA1;AHSG;FGG;APOA1;PLG;ORM2;KNG1;TF;TTR;APOH;ALB;HRG |
| LIHC | PC1 | 0.84 | positive | Reactome_2022 | Platelet Degranulation R-HSA-114608 | $1.2 \times 10^{-19}$ | 3767.29 | FGB;FGA;ORM1;SERPINA1;AHSG;FGG;APOA1;PLG;ORM2;KNG1;TF;TTR;APOH;ALB;HRG |
| LIHC | PC2 | 0.82 | positive | GO_Biological_Process_2023 | Multifunctional Anion Exchangers R-HSA-427601 | 0.0015 | 2717.50 | SLC26A7;SLC26A4 |
| LIHC | PC2 | 0.82 | positive | GO_Biological_Process_2023 | Thyroxine Biosynthesis R-HSA-209968 | 0.0015 | 2324.81 | TPO;IYD |
| LIHC | PC2 | 0.82 | positive | Reactome_2022 | Multifunctional Anion Exchangers R-HSA-427601 | 0.0015 | 2717.50 | SLC26A7;SLC26A4 |
| LIHC | PC2 | 0.82 | positive | Reactome_2022 | Thyroxine Biosynthesis R-HSA-209968 | 0.0015 | 2324.81 | TPO;IYD |
| LIHC | PC2 | 0.82 | positive | GO_Biological_Process_2023 | Metabolism Of Amine-Derived Hormones R-HSA-209776 | 0.0034 | 1017.20 | TPO;IYD |
| LIHC | PC3 | 0.79 | positive | GO_Biological_Process_2023 | Diseases Associated With Surfactant Metabolism R-HSA-5687613 | $7.5 \times 10^{-14}$ | 58435.97 | SFTPB;SLC34A2;SFTPA2;SFTPC;SFTPD;SFTPA1 |
| LIHC | PC3 | 0.79 | positive | KEGG_2021_Human | Diseases Associated With Surfactant Metabolism R-HSA-5687613 | $7.5 \times 10^{-14}$ | 58435.97 | SFTPB;SLC34A2;SFTPA2;SFTPC;SFTPD;SFTPA1 |
| LIHC | PC3 | 0.79 | positive | Reactome_2022 | Diseases Associated With Surfactant Metabolism R-HSA-5687613 | $7.5 \times 10^{-14}$ | 58435.97 | SFTPB;SLC34A2;SFTPA2;SFTPC;SFTPD;SFTPA1 |
| LIHC | PC3 | 0.79 | positive | GO_Biological_Process_2023 | Surfactant Metabolism R-HSA-5683826 | $8.3 \times 10^{-13}$ | 8840.05 | SFTPB;SLC34A2;NAPSA;SFTPA2;SFTPC;SFTPD;SFTPA1 |
| LIHC | PC3 | 0.79 | positive | KEGG_2021_Human | Surfactant Metabolism R-HSA-5683826 | $8.3 \times 10^{-13}$ | 8840.05 | SFTPB;SLC34A2;NAPSA;SFTPA2;SFTPC;SFTPD;SFTPA1 |

**Supplementary Table S40.** TMS-mRNAsi Spearman correlations in KIRC and LIHC (OCLR-derived mRNAsi from Malta et al.).

| Cohort | Source | Comparison | Spearman ρ | p-value | n |
|---|---|---|---|---|---|
| KIRC | pancanatlas | TMS vs mRNAsi | 0.263 | $1.2 \times 10^{-9}$ | 521 |
| LIHC | pancanatlas | TMS vs mRNAsi | -0.014 | 0.797 | 362 |

**Supplementary Table S41.** Joint Cox models (TMS + mRNAsi + stage + purity): Models A, B, and C hazard ratios, concordance, and AIC for KIRC and LIHC.

| Cohort | Model | Covariate | HR (95% CI) | p-value | C-index | AIC | n | Events |
|---|---|---|---|---|---|---|---|---|
| KIRC | A: TMS stage purity | TMS | 0.62 (0.52–0.74) | $1.8 \times 10^{-7}$ | 0.752 | 1852.35 | 535 | 174 |
| KIRC | A: TMS stage purity | Stage | 1.92 (1.68–2.20) | $3.1 \times 10^{-21}$ | 0.752 | 1852.35 | 535 | 174 |
| KIRC | A: TMS stage purity | Purity | 4.51 (1.98–10.27) | $3.3 \times 10^{-4}$ | 0.752 | 1852.35 | 535 | 174 |
| KIRC | B: TMS mRNAsi stage purity | TMS | 0.61 (0.51–0.74) | $2.2 \times 10^{-7}$ | 0.760 | 1774.96 | 518 | 168 |
| KIRC | B: TMS mRNAsi stage purity | mRNAsi | 0.95 (0.81–1.12) | 0.546 | 0.760 | 1774.96 | 518 | 168 |
| KIRC | B: TMS mRNAsi stage purity | Stage | 1.99 (1.73–2.29) | $6.2 \times 10^{-22}$ | 0.760 | 1774.96 | 518 | 168 |
| KIRC | B: TMS mRNAsi stage purity | Purity | 5.36 (2.19–13.12) | $2.4 \times 10^{-4}$ | 0.760 | 1774.96 | 518 | 168 |
| KIRC | C: mRNAsi stage purity | mRNAsi | 1.00 (0.83–1.19) | 0.964 | 0.741 | 1794.90 | 518 | 168 |
| KIRC | C: mRNAsi stage purity | Stage | 1.94 (1.69–2.23) | $2.9 \times 10^{-21}$ | 0.741 | 1794.90 | 518 | 168 |
| KIRC | C: mRNAsi stage purity | Purity | 0.98 (0.52–1.88) | 0.963 | 0.741 | 1794.90 | 518 | 168 |
| LIHC | A: TMS stage purity | TMS | 0.89 (0.76–1.04) | 0.139 | 0.638 | 1160.04 | 341 | 116 |
| LIHC | A: TMS stage purity | Stage | 1.63 (1.32–2.00) | $4.4 \times 10^{-6}$ | 0.638 | 1160.04 | 341 | 116 |
| LIHC | A: TMS stage purity | Purity | 1.05 (0.35–3.10) | 0.934 | 0.638 | 1160.04 | 341 | 116 |
| LIHC | B: TMS mRNAsi stage purity | TMS | 0.83 (0.69–0.98) | 0.029 | 0.659 | 1153.81 | 339 | 116 |
| LIHC | B: TMS mRNAsi stage purity | mRNAsi | 1.28 (1.07–1.54) | 0.0080 | 0.659 | 1153.81 | 339 | 116 |
| LIHC | B: TMS mRNAsi stage purity | Stage | 1.56 (1.26–1.93) | $4.1 \times 10^{-5}$ | 0.659 | 1153.81 | 339 | 116 |
| LIHC | B: TMS mRNAsi stage purity | Purity | 1.14 (0.40–3.24) | 0.811 | 0.659 | 1153.81 | 339 | 116 |
| LIHC | C: mRNAsi stage purity | mRNAsi | 1.22 (1.02–1.46) | 0.034 | 0.642 | 1156.13 | 339 | 116 |
| LIHC | C: mRNAsi stage purity | Stage | 1.62 (1.32–2.00) | $4.2 \times 10^{-6}$ | 0.642 | 1156.13 | 339 | 116 |
| LIHC | C: mRNAsi stage purity | Purity | 0.92 (0.33–2.56) | 0.875 | 0.642 | 1156.13 | 339 | 116 |

Variance inflation factors for Model B: KIRC — VIF_TMS = 2.21, $\mathrm{VIF}_{mRNAsi}$= 1.16, VIF_Stage = 1.07, $VIF_{Purity}$ = 2.45; LIHC — $VIF_{TMS}$ = 1.19, $VIF_{mRNAsi}$ = 1.05, $VIF_{Stage}$ = 1.09, $VIF_{Purity}$ = 1.07. All values fall below the conventional threshold of 5, confirming that the subsumption of mRNAsi by TMS in KIRC is not attributable to collinearity-induced instability.

**Supplementary Table S42.** Model comparison summary: concordance and AIC for TMS-only, joint (TMS + mRNAsi), and mRNAsi-only models.

| Cohort | Model | C-index | AIC | n | Events |
|---|---|---|---|---|---|
| KIRC | A: TMS stage purity | 0.752 | 1852.35 | 535 | 174 |
| KIRC | B: TMS mRNAsi stage purity | 0.760 | 1774.96 | 518 | 168 |
| KIRC | C: mRNAsi stage purity | 0.741 | 1794.90 | 518 | 168 |
| LIHC | A: TMS stage purity | 0.638 | 1160.04 | 341 | 116 |
| LIHC | B: TMS mRNAsi stage purity | 0.659 | 1153.81 | 339 | 116 |
| LIHC | C: mRNAsi stage purity | 0.642 | 1156.13 | 339 | 116 |

**Supplementary Table S43.** Absolute survival probabilities by TMS tertile within intermediate-stage ccRCC (stages II–III) at three-year and five-year horizons.

| TMS tertile | Horizon | n | Events | Survival (95% CI) | Mortality | ARD vs High |
|---|---|---|---|---|---|---|
| Low | 3-year | 61 | 28 | 64.2% (49.6–75.6) | 35.8% | 0.1584 |
| Low | 5-year | 61 | 28 | 53.1% (37.4–66.6) | 46.9% | 0.1732 |
| Mid | 3-year | 60 | 17 | 76.2% (61.7–85.8) | 23.8% | |
| Mid | 5-year | 60 | 17 | 65.0% (48.7–77.3) | 35.0% | |
| High | 3-year | 60 | 17 | 80.0% (66.8–88.4) | 20.0% | |
| High | 5-year | 60 | 17 | 70.5% (55.4–81.3) | 29.5% | |

**Supplementary Table S44.** Cox regression within the intermediate-stage ccRCC subset: univariable and adjusted (TMS + stage + purity) models.

| Model | Covariate | HR (95% CI) | p-value | C-index | n | Events |
|---|---|---|---|---|---|---|
| univariable | TMS | 0.78 (0.62–0.99) | 0.039 | 0.563 | 181 | 62 |
| adjusted | TMS | 0.56 (0.43–0.73) | $2.3 \times 10^{-5}$ | 0.680 | 181 | 62 |
| adjusted | Stage | 2.52 (1.36–4.68) | 0.0034 | 0.680 | 181 | 62 |
| adjusted | Purity | 9.72 (2.81–33.64) | $3.3 \times 10^{-4}$ | 0.680 | 181 | 62 |
